# Investigating Potential Microbial Contributors to Enhanced Metabolite Production under Oxygen Perturbations: An Integrated Metagenomic and Metabolomic Approach

**DOI:** 10.1101/2024.09.21.614222

**Authors:** Xueyang Zhou, Bharat Manna, Boyu Lyu, Naresh Singhal

## Abstract

The specific impacts of oxygen perturbation on microbial communities and their synthesis of metabolites remain unclear. We systematically explored how oxygen perturbations alter microbial growth, subsequently affecting the abundance of functional genes and promoting the synthesis of metabolites such as amino acids. Analysis of microbial community structure indicates dynamic stability under oxygen perturbations, with only a fraction of the microbial abundance being altered. By comparing the abundance of functional genes with metabolic features, we revealed how changes in the microbial community impact the overall system performance. Amino acid biosynthesis has an enhanced potential under conditions of oxygen perturbation. Through biological and statistical correlation analyses, we identified microbial species significantly correlated with the efficiency of target metabolic reactions under oxygen perturbations. *Mycolicibacterium madagascariense*, *Mycolicibacterium fortuitum*, and *Burkholderia pseudomallei* displayed strong associations with proline and tryptophan synthesis reactions. Moreover, the abundance of microbial genera including *Labrys*, *Actinomyces*, and *Nitrosopumilus* exhibited a highlysignificant positive correlation with metabolite abundance in enriched metabolic pathways under oxygen perturbations. These results suggest that microbial systems might achieve dynamic stability in community structure under oxygen perturbations, while exhibiting slightly differential metabolic potentials. Notably, enhanced efficiency in amino acid biosynthesis could help to assimilate more carbon and nitrogen resources in activated sludge during wastewater treatment.

**SYNOPSIS:** We provide an in-depth investigation into the impact of oxygen perturbations on microbial growth in activated sludge systems and the subsequent variations in the abundance of metabolic genes. Our research highlights the potential for subtle changes in amino acid biosynthesis and identifies key microbial species associated with synthesis efficiency. These insights into the dynamic stability and differentiated metabolic potentials of microbial communities during oxygen perturbations can inform the development of more effective resource recovery strategies.

## 1. INTRODUCTION

Excess Activated Sludge (EAS) poses a universal challenge in biological wastewater treatment worldwide. The European Union generates about 10.9 million tons of dry solid EAS each year^1^. Untreated EAS could cause severe environmental harm^2,3^. However, recent research has demonstrated that, through proper processing and recycling, EAS can be transformed into potential resource recovery opportunities^4^. In the process of treating and recycling EAS, its components can be converted into a variety of valuable products. For example, the organic compounds amino acids, proteins, and lipids can be transformed into economically effective materials for biofuels, material protection, and industrial pickling^5–10^. Some research efforts have focused on enhancing the assimilation of nutrients in wastewater by microorganisms to synthesize more organic substances like amino acids and proteins^11^. However, these strategies typically emphasize improving the quality of carbon sources in the wastewater^12,13^. This is unfavorable for some wastewater conditions with a low carbon-nitrogen ratio. Thus, how to improve microbial assimilation efficiency through economical and convenient process adjustment has become a current research focus^14^.

Activated sludge consists of a highly complex microbial community comprised of eukaryotes, bacteria, archaea, and viruses, among which bacteria play a dominant role and have important functions in the carbon and nitrogen conversion^15–17^. Dissolved oxygen (DO) is an important parameter in biological wastewater treatment processes, typically fluctuating with the seasons and varying with water temperature and altitude ^18^. Different levels of DO not only affect nitrogen removal efficiency but also influence the composition, behavior and activity of microorganisms, thus leading to changes in microbial community structure^19^. The intrinsic mechanism of DO-driven changes in microbial community structure is that different microorganisms have different oxygen affinities^20,21^. A wealth of research has explored the differences in dominant functional bacteria under different aeration conditions^22–25^. but little attention has been paid to the metabolic function of valuable metabolite synthesis.

Our study found that using oxygen perturbation strategies to treat activated sludge can improve the synthesis efficiency of metabolites such as amino acids. Metagenomic analysis shows that oxygen perturbations can impact the community structure of microorganisms, thereby improving their related metabolic potential and biosynthesis efficiency.

## 2. MATERIALS AND METHODS

### 2.1 Experimental Setup and Operation

We collected activated sludge inoculated from the Māngere Wastewater Treatment Plant in Auckland, New Zealand (date). Before each batch test, we constructed reaction systems including active sludge with 2.75g/L dry weight concentration, supplemented with inorganic carbon (3840 mg NaHCO_3_), and 1 mL of trace element solution. Additional sources of organic carbon (methanol), inorganic nitrogen (ammonia nitrogen), and phosphorus were injected into the reaction system at an even rate to achieve total concentrations of 1600 mg COD, 320 mg NH ^+^-N for IP or 400 mg NH_4_ ^+^-N for CP & CA, and 40 mg PO ^3-^-P using a syringe pump over 48 hours. The experiment was completed at a temperature of 20±1, and automatically run according to the set aeration mode for 48 hours.

The reaction system was exposed to three different aeration conditions included Continuous Aeration (CA) (stable at 2.0±0.2 mg/L DO), Continuous Perturbation (CP) (DO varying between 0.1 and 2.0 mg/L in ∼3 min cycles), and Intermittent Perturbation (IP) (DO varying between 0 and 2.0 mg/L; aerobic time: anoxic time = 3 min: 3 min) (Figure S2).

### 2.2 Determination of Organic Matter in Biomass

To identify fluorescent organic substances in sludge biomass, an Excitation-emission matrix (EEM) analysis was performed using an Aqualog A-TEEM Spectrometer (Horiba, Japan). The process involved using a 10 mL activated sludge sample, which was then purified by centrifugation twice at 6,113 x g for 10 minutes at a temperature of 4°C and was later reconstituted in a 0.85% NaCl solution. The sludge sample was then converted to pellet form, dissolved in 10 mL of 2% EDTA solution, and allowed to sit at 4°C for a period of three hours. The sample was then subjected to sonication for 20 cycles, each with a 15-second on and off duration on ice. After undergoing centrifugation at 17,467 x g for 20 minutes, the resultant supernatant was passed through a 0.2 μm cellulose acetate filter for EEM analysis.

For the EEM analysis, the excitation wavelengths were set between 240 to 600 nm (3 nm gaps), and the emission wavelengths were set between 212.70 to 622.21 nm (3.28 nm gaps). Fluorescence and absorbance measurements were interpreted using the PARAFAC model by staRdom to evaluate the relative quantities of organic matter in the biomass^26^. The Openfluor database was employed to identify the properties of the components^27^.

### 2.3 Metagenomics Assay

The procedure for DNA extraction started with collecting 10 mL sludge samples. Biological duplicates were acquired from the 24-hour and 48-hour samples across all experimental conditions. The extraction followed the method provided by the DNeasy PowerSoil Kit (Qiagen, Germany), and the resultant DNA was preserved at −20°C for further analysis.

The DNA samples were then sent off to the Auckland Genomics Centre (Auckland, NZ) to conduct metagenomic studies. This process comprised several steps, which included the concentration of sample DNA, quality control assessment, and preparation for Illumina sequencing. This preparation encompassed polymerase chain reaction (PCR) and indexing. Post library QC, normalization was carried out, followed by a final pool check using a bioanalyzer. Lastly, sequencing was carried out on a HiSeq platform, resulting in approximately 400 million 2×150bp paired-end reads.

The raw sequences obtained from the sequencing facility underwent preprocessing steps to eliminate adapters and low-quality sequences. Trimmomatic was used for this purpose^28^, applying specific quality thresholds: LEADING:3, TRAILING:3, SLIDINGWINDOW:10:15, and MINLEN:50 with the following settings: TruSeq3-PE-2.fa:2:30:10:2:keepBothReads^29^. For metagenomic taxonomic and functional profiling, SqueezeMeta v1.5.2 pipeline was employed with default settings^30^. Co-assembly was performed using Megahit^31^, and short contigs (<200 bps) were filtered out using prinseq^32^. Within the SqueezeMeta pipeline, Barrnap tool was used for predicting RNAs^33^, while Prodigal was utilized for predicting ORFs^34^. Similarity searches against NCBI GenBank nr database^35^ and Kyoto Encyclopedia of Genes and Genomes^36^, were performed for taxonomic and functional assignments, respectively, using Diamond^37^. Bowtie2 was used for read mapping against the contigs^38^. Finally, the metagenomics data was analyzed using the SQMtools R package^39^.

### 2.4 Metabolomics Assay

A GC/MS platform was used to ascertain the relative abundance of intracellular metabolites in activated sludge^40^. Three biological replicates along with two technical replicates were investigated for each set of conditions. The sludge samples (10 mL) were spun in a centrifuge, flash-frozen in liquid nitrogen, and preserved at −80°C. Metabolites were extracted by combining the sludge pellets with 2.5 mL of a chilled methanol-water mixture and 0.2 µmol of an internal standard, 2,3,3,3-d4-alanine. The metabolites were released through a procedure of freeze-thawing three times, followed by vigorous shaking, and the supernatant was obtained by centrifuging at 22,707 x g for 15 minutes at −20°C. An additional 2.5 mL of cold methanol-water solution was added to further extract the metabolites from the sludge pellet.

The metabolite extract was then dissolved in 400 µL of a 1 M sodium hydroxide solution, enhanced with 68 µL of pyridine and 334 µL of methanol. 40 µL of MCF was added twice, with each addition followed by 30 seconds of intense vortexing. This solution was then combined with 400 µL of chloroform and mixed for an additional 10 seconds. This mixture was then added with 800 µL of a bicarbonate solution (50 mM), stirred, and spun in a centrifuge. The aqueous layer was removed, and any remaining water in the chloroform phase was taken out using anhydrous sodium sulfate.

The derivatized sample was then assessed using a GC-MS system (Agilent GC7890 attached to an MSD597 unit) outfitted with a ZB-1701 GC capillary column (30 m × 250 µm (id) × 0.15 µm film thickness) and a 5 m guard column (Phenomenex, Torrance, CA, USA). AMDIS software (NIST, Boulder, CO, USA) was used to deconvolute the resulting GC-MS chromatograms. The mass spectra were compared with the in-house MCF MS library to identify the metabolites. The “MassOmics” R package was utilized for data filtering, and the metabolite abundance values were normalized using the internal standard^41^.

### 2.5 Statistical Analysis of Omics Data

In this study, the analysis of microbial diversity and microbial community structure was carried out using MicrobiomeAnalyst^42^. Indices such as Observed, Chao1, and Shannon were utilized for the evaluation of Alpha diversity. The principal coordinate analysis (PCoA) of microbial community structure was based on the Bray–Curtis distance. The Marker Data Profiling function was employed for the grouping and visualization of microbial taxonomic information in the samples. A metagenomeSeq_0-inflated test was used for single-factor statistical comparisons of microbial genera and species, with an adjusted p-value of less than 0.01 considered indicative of different microbial species.

For functional genes analysis, normalized counts (Transcripts Per Million, TPM values) were used. After the genes were screened and categorized based on the Kyoto Encyclopedia of Genes and Genomes (KEGG) database, an ANOVA analysis was conducted for the differential abundance analysis of genes, with an adjusted p-value of less than 0.05 serving as the significance threshold.

Metabolomics analysis was performed using MetaboAnalyst^43^. ANOVA analysis (adjusted p-value < 0.05) was applied to select differential metabolites. Pathway enrichment analysis was conducted by selecting CP vs. CA and IP vs. CA as the grouping to complete the task.

The microbiota traceability analysis for metabolites was conducted using the Metorigin platform^44^. The KEGG database was leveraged to search for microbial groups that could potentially be involved in related metabolic reactions, and correlation analysis was conducted with the metabolites. Spearman analysis was used as the statistical method for analyzing the correlation between differential microbes and differential metabolites. Sankey diagrams were used to display the biological and statistical connections between microbial groups and differential metabolites.

In addition, the heatmaps in this study were visualized using TBtools^45^.

## 3. RESULTS

### 3.1 Functional Potential Shifts under Oxygen Perturbations

#### 3.1.1 Bacterial Community Structure Analysis

There was no discernible difference in the microbial diversity and richness of the activated sludge subjected to either perturbation or non-perturbation of oxygen for a duration of 48 hours. Alpha diversity analysis executed at the species level on the three sample groups suggested that the microbial diversity was not significantly divergent (Figure S3). The Principal Coordinate Analysis (PCoA), utilizing the Bray-Curtis distance, likewise did not exhibit any substantial separation of populations among the three distinct aeration conditions.

Following the analysis of metagenomic sequencing outcomes using SqueezeMeta, a total of 5908 species were identified (of which 5557 were annotated) from the samples (Figure S10). These species were distributed across four kingdoms, including 134 Archaea (mean ± SD, 0.010± 0.001%), 5275 Bacteria (97.104± 0.113%), 140 Eukaryota (0.006±0.001%), and 8 Viruses (0.001± 0.000%). At the phylum level within bacteria, the most prominent were Proteobacteria (47.912±3.073%), Actinobacteria (14.043±1.385%), Bacteroidetes (10.562±0.972%), Chloroflexi (8.242±0.956%), and Nitrospirae (4.843±1.183%) (Figure S11). At the Genus level, the bacteria were predominantly characterized by *Hyphomicrobium* (16.449±1.395%), *Nitrospira* (2.466±0.563%), and *Methylobacillus* (1.836±0.567%) (Figure 1a).

**Figure 1.**
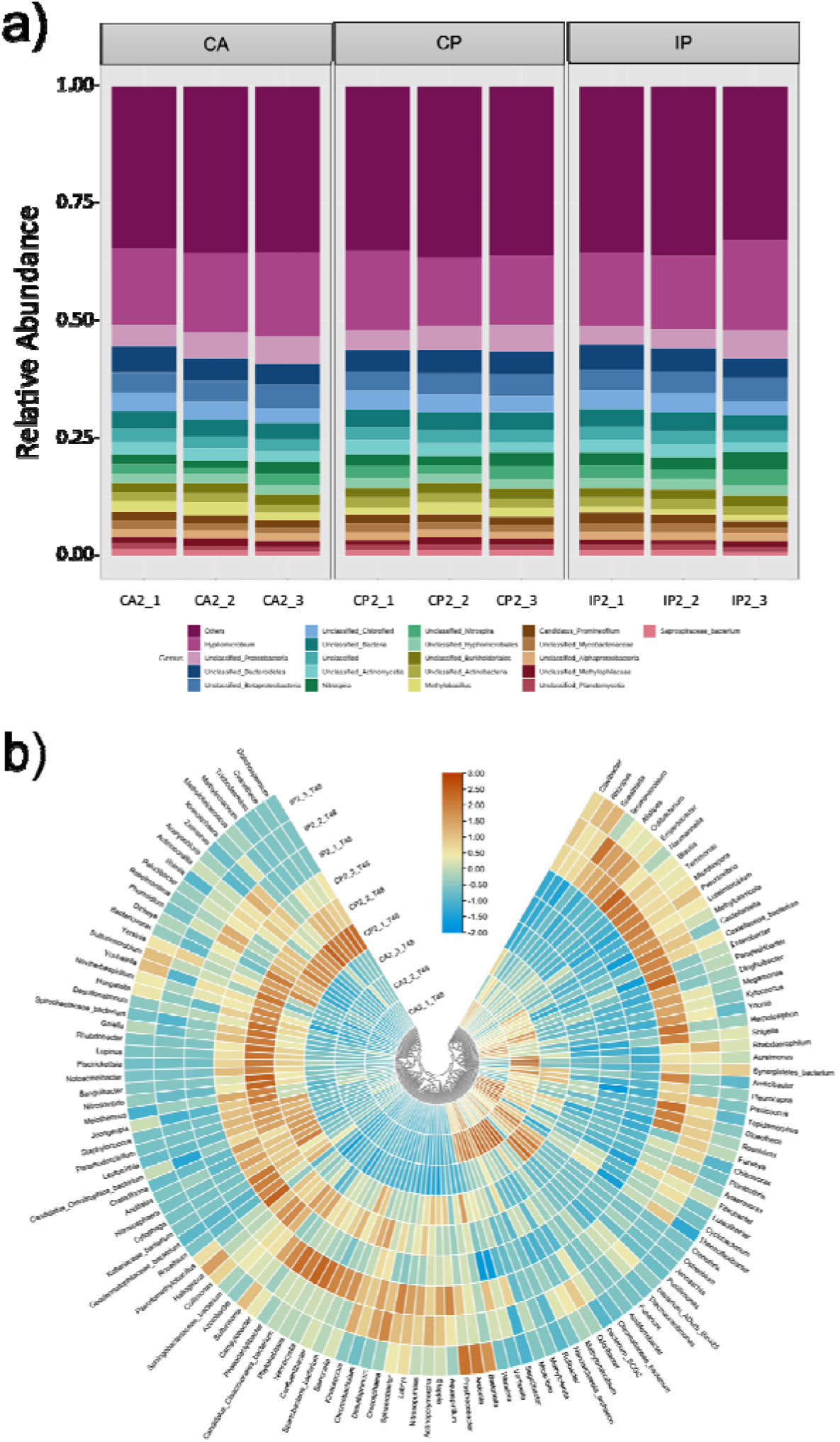
Differences in the taxonomic composition of the microbiota in activated sludge under different aeration conditions: (a) Relative abundance of microbial taxonomic groups at the genus (top 20 genera and others) level; (b) Classification of genera annotated with significant differences (metagenomeSeq_0-inflated test, P<0.01) in abundance among different aeration conditions.

Analysis of significant differences was performed in the 3602 genera across the samples under the three aeration conditions using the metagenomeSeq zero-inflated test (*P*<0.01). As many as 210 genera showed significant differences in abundance across the three conditions (Figure 1b). Genera with markedly higher abundance under CA and CP conditions showed no overlap, suggesting that the dominant genera under these conditions might have specialized survival and reproductive needs, thereby making them the leading genera under specific aeration scenarios. Conversely, the genera displaying higher abundance under IP conditions also exhibited a reasonable abundance under other conditions (CA or CP). This suggests these genera may possess strong adaptability to environmental changes, enabling them to survive under multiple conditions. However, their abundance may decrease under CA and CP conditions due to the higher adaptability of other genera to these conditions.

#### 3.1.2 KEGG Functional Potential Analysis

An analysis of the abundance of 9861 genes obtained from metagenomic sequencing under different aeration conditions was conducted to determine the functional potential of the microbial community in the activated sludge system. Based on the PCA analysis, we observed separation in the relative abundance of KO genes between the three aeration conditions (Figure S20). Upon analyzing the abundance of genes corresponding to KEGG level 2 functional pathways, we found that CP showed relatively higher abundance across 11 metabolic pathways (Figure S21, S22).

Further mapping of the genes to KEGG level 3 pathways revealed subtle differences in abundance patterns between different pathways. For energy metabolism (Figure S24) and nucleotide metabolism (Figure S25), both of which are closely related to microbial growth and proliferation, IP was at a disadvantage in terms of abundance compared to the other two conditions. This differential abundance might suggest a potential weakening of the growth and reproduction capabilities of the microbes under the IP condition. This observation aligns with our findings from the determination of fluorescent organic matter content in biomass (Figure S19), where the lowest value was recorded under the IP condition.

Overall, no obvious difference in the abundance of genes involved in lipid metabolism was observed under different aeration conditions (Figure S26). That is, oxygen perturbation or not did not cause the microbial community to show a clear advantage or disadvantage in terms of lipid metabolism potential. However, there seemed to be a tendency to show certain abundance in different level 3 pathways, possibly indicating that different microbes in the community may have specific advantages in specific sub-pathways of lipid metabolism. Meanwhile, the influence of these three aeration conditions on the humic-like component in biomass was relatively minor, implying that while these conditions may affect some specific functions of the microbial community, their overall impact on lipid metabolism is weak.

Differences in the content of protein-like substances under different aeration conditions show that oxygen perturbations might impact microbial protein synthesis and degradation metabolism. Interestingly, we found that this difference aligns with the gene abundance trend in amino acid metabolism (Figure 2a) and metabolism of other amino acids (Figure S27), that is, the abundance decreases in the order of CP, CA, and IP. This potential might be attributed to the characteristics of the dominant microbes under CP conditions, or it could be a result of the stimulation of certain metabolic activities by the environmental condition.

**Figure 2.**
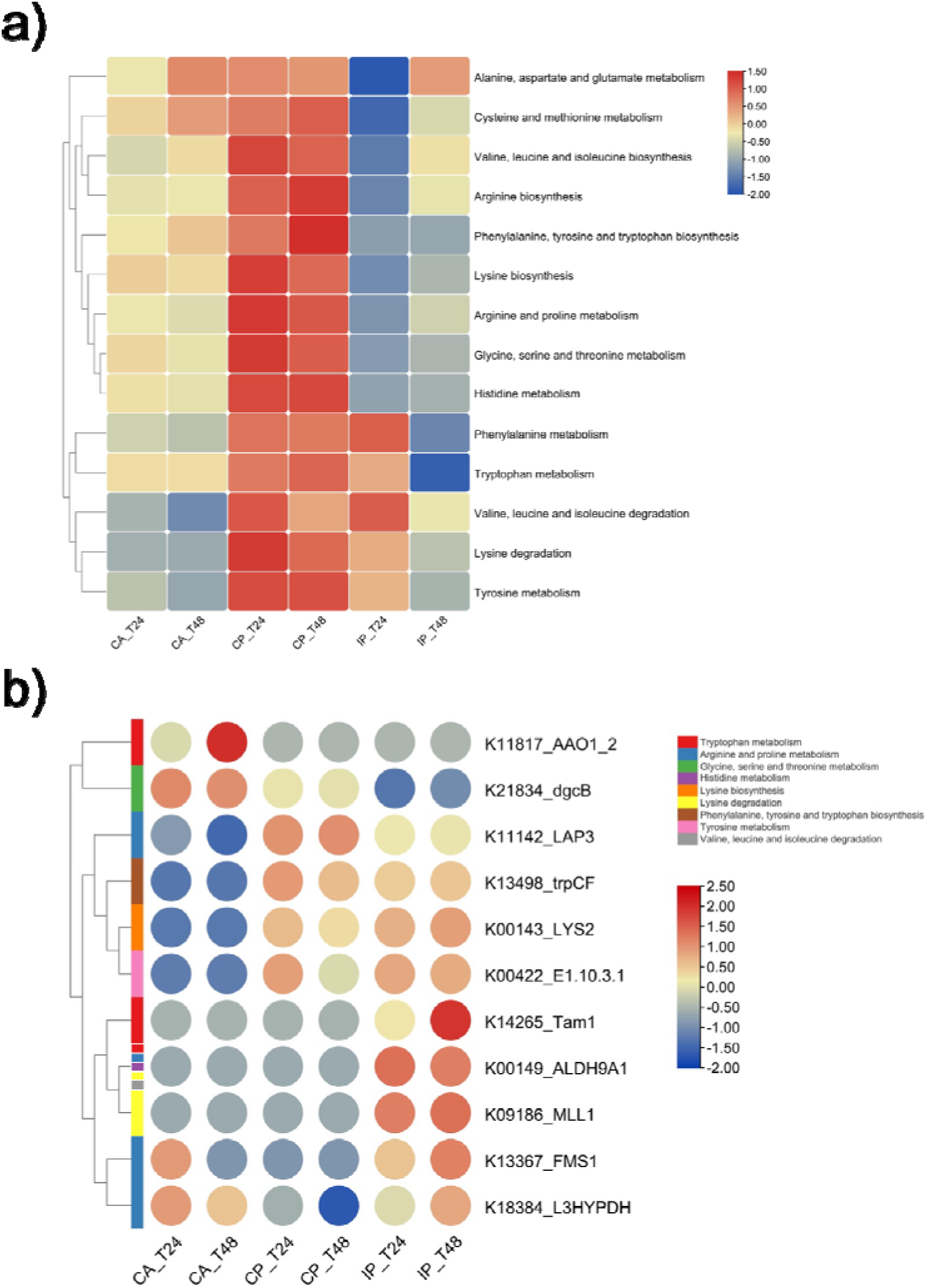
Amino acid metabolic potential of activated sludge samples after 24 and 48 hours of exposure to different aeration conditions: (a) Abundance of genes associated with the KEGG level3 pathways under amino acid metabolism; (b) Genes with significantly different abundance in amino acid metabolism (ANOVA test, *P*<0.05). The heatmap shows the average value of the same condition. Row clustering according to ‘Ward’ (biological triplicates; n=3).

### 3.2 Amino Acid Synthesis Potentially Affected by Gene Abundance

#### 3.2.1 Significant Abundance of Genes in Amino Acid Metabolism

After observing that differences in the abundance of functional genes under different aeration conditions might affect biosynthesis, we investigated functional genes with significant abundance differences in the amino acid metabolism pathway. In parallel, we integrated metabolomics to analyze which amino acid synthesis was affected by differences in microbial growth by influencing the expression potential of specific metabolic pathways.

Through a significant difference analysis of 601 genes related to amino acid metabolism, we found that the abundance of 11 genes significantly differed under different aeration conditions (ANOVA, *P*<0.05, Figure S34). Overall, genes with higher abundance were primarily observed in samples under oxygen perturbation conditions (CP and IP) (Figure 2b). Under CP conditions, the abundance of four genes (K11142, K13498, K00143, K00422) significantly surpassed those under CA conditions. These four genes each belong to different amino acid metabolic pathways: arginine and proline metabolism (ko00330), phenylalanine, tyrosine, and tryptophan biosynthesis (ko00400), lysine biosynthesis (ko00300), and tyrosine metabolism (ko00350). They are more associated with the synthesis process in the overall function of amino acid metabolism.

Similarly, under IP conditions, in addition to those four genes also showing higher abundance, another four genes (K14265, K00149, K09186, and K13367) demonstrated higher abundance compared to CA. Among these, K00149 is involved in multiple metabolic pathways, while the other three genes are respectively located in tryptophan metabolism (ko00380), lysine degradation (ko00310), and arginine and proline metabolism (ko00330), and they are more inclined to participate in the modification process of amino acids.

The only gene that showed significantly higher abundance under CA conditions was K11817, which belongs to tryptophan metabolism (ko00380), and encodes indole-3-acetaldehyde oxidase. This enzyme mainly acts on aldehyde or carbonyl donors and uses oxygen as an electron acceptor, a process that mainly involves the degradation and modification of amino acids.

These results suggest that oxygen perturbations may alter the functional potential of specific amino acid metabolic pathways by affecting the growth of different microbial species, leading to variations in amino acid abundance throughout the system.

#### 3.2.2 Pathway Enrichment Analysis of Metabolites

To verify whether oxygen perturbations lead to an increase in gene abundance and affect the concentration of metabolites in the corresponding pathways, we used GC-MS semi-targeted metabolomic technology to quantitatively analyze the abundance of metabolites in activated sludge samples. Metabolites analysed included 16 amino acids, 23 fatty acids, 18 kinds of acids, 2 kinds of peptides (names?), 1amine (just name it), and 3 other metabolites (name them, otherwise too vague) (Figure S35). Significant difference analysis based on ANOVA found that metabolites show high abundance under oxygen perturbation conditions, including 15 amino acids, 10 fatty acids, 1 peptide, and 4 acids (Figure 3a).

**Figure 3.**
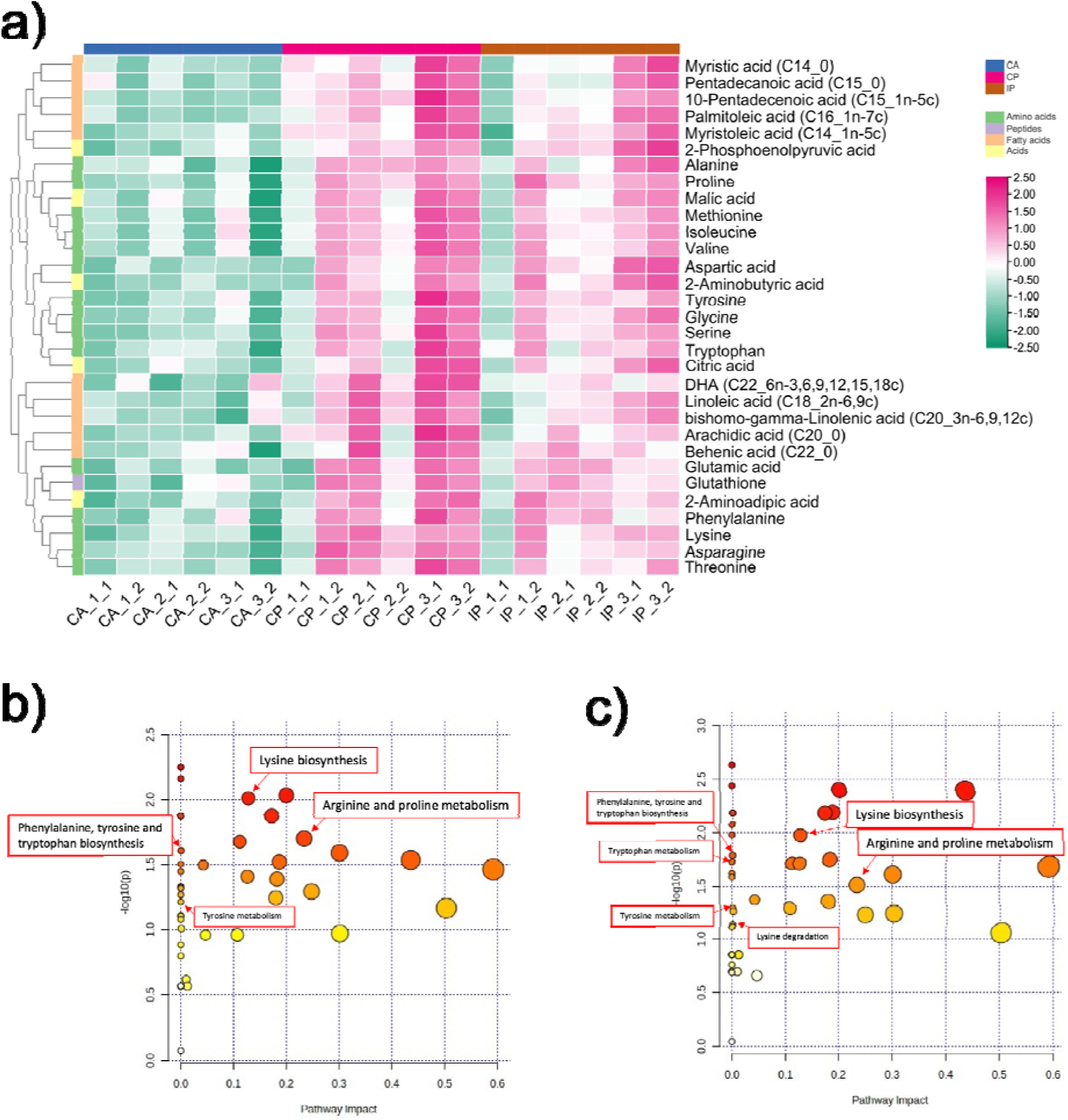
Differences in metabolite abundance in activated sludge biomass samples after 48 h of exposure to different aeration strategies: (a) Significant differences in relative abundance of metabolites (ANOVA test, *P*<0.05). The data is normalized by internal standard, Log10 transformation, and Pareto scaling. Row clustering according to ‘Ward’ (technical duplication results for biological triplicates; n=6); (b, c) Pathway enrichment analysis based on metabolomics (CP vs. CA; IP vs. CA). The marked pathways related to amino acid metabolism are those where metabolites can match the reactions in which the significant genes are involved.

We matched these significantly different metabolites with the differential genes of the microbial community screened through metagenomic analysis. We found that, under the CP condition, compared to CA, four genes with significantly higher abundance could be matched to the same metabolic reactions with metabolites of high differential abundance. Specifically, K11142 and proline correspond to R00135 in ko00330, K13498 and tryptophan correspond to R00674 in ko00400, K00143 and 2-Aminoadipic acid correspond to R03098 in ko00300, K00422 and tyrosine correspond to R00031 in ko00350. Meanwhile, pathway enrichment analysis based on metabolomic data indicates that ko00300, ko00330, and ko00400 are also the three amino acid metabolic pathways with the most significant differences under CP compared to CA (Figure 3b).

Since these 4 significant genes also show high abundance under IP, the corresponding proline and 2-Aminoadipic acid both show significantly higher abundance compared to CA. However, the relative abundance of tryptophan and tyrosine is not as significant as that in CP, which we speculate may be related to the increased potential of the amino acid degradation pathway. This is because the gene (K14265) responsible for converting tryptophan only shows significantly high abundance under IP.

Overall, the significance of metabolite abundance matching with functional gene abundance proves that the increase in functional gene abundance caused by changes in community structure is indeed an important reason for the increase in metabolite abundance under oxygen perturbations. This finding provides valuable clues for us to understand how oxygen perturbations affect microbial metabolism, and also urges us to continue exploring how changes in the abundance of functional genes are caused by changes in community structure.

### 3.3 Contribution of Perturbation-responsive Microbes to Amino Acid Metabolism

#### 3.3.1 Biological Correlation between Microbial Species and Significant Metabolic Reaction

The Biology-Sankey (BIO-Sankey) analysis function of the MetOrigin platform allows us to find the types of microbes in the samples that perform specific metabolic reactions by comparing the microbial information in the samples with the database. We performed BIO-Sankey network map analysis on the metabolic reactions that match significant genes and significant metabolites in amino acid metabolism under oxygen perturbations and obtained the correlation between microbial communities and metabolic reactions at different microbial taxonomic ranks (Figure S41∼S53).

First, we analyzed all microbial species that can perform R00135 (proline synthesis reaction) (Figure S41, S42). Among the types of microbes that have significantly higher abundance compared to CA and are positively correlated with proline abundance, there are 7 species under CP and 10 species under IP. Among these, *Mycolicibacterium madagascariense* and *Aerococcus urinae* under CP, and *Mycolicibacterium litorale* and *Mycolicibacterium madagascariense* under IP, meet the standard of significant positive correlation (Spearman correlation test; R>0, *P*<0.05).

For R00674 (tryptophan synthesis reaction), we found that there are 11 microbial species under CP and 14 microbial species under IP that are positively correlated with tryptophan abundance and their abundance is significantly higher than CA (Figure S43, S44). Among these species, *Klebsiella pneumoniae*, *Azospirillum brasikense*, *Mycolicibacterium madagascariense*, *Pirellula staleyi*, *Streptococcus pneumonia*, and *Aerococcus urinae* under CP meet the criteria of significant positive correlation.

Figure 4 integrates our BIO-Sankey analysis results of R00135 and R00674 at the species level, including network association information of all potential functional species that are positively correlated with metabolites and have significantly higher abundance. Of note, *Mycolicibacterium madagascariense*, *Mycolicibacterium fortuitum*, and *Burkholderia pseudomallei* show relevance under both reactions and two oxygen perturbations, suggesting they may have significant growth advantages under CP or IP and can contribute to amino acid metabolism.

**Figure 4.**
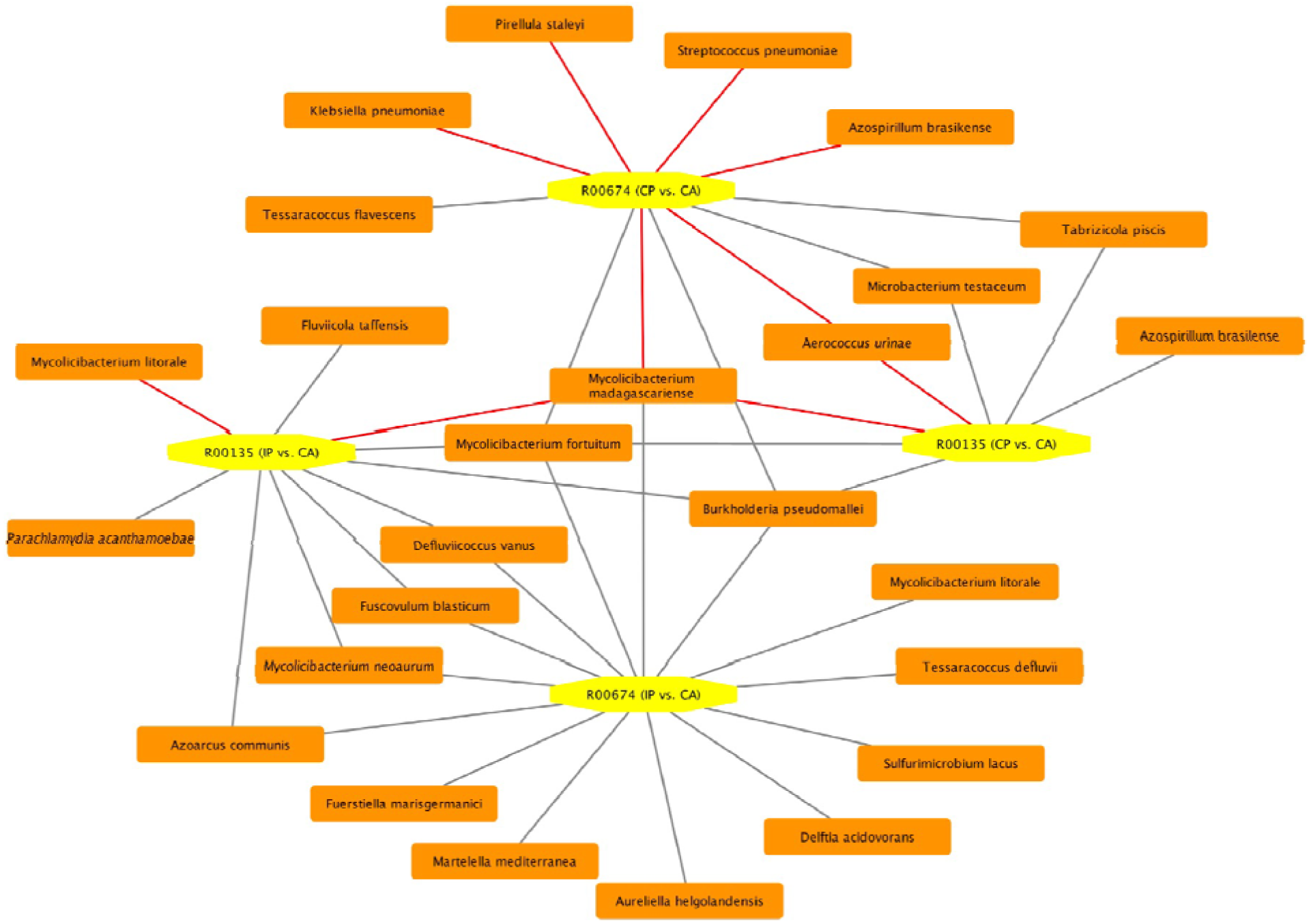
Based on the BIO-Sankey analysis results that biologically significant species capable of performing R00135 (proline synthesis reaction) and R00674 (tryptophan synthesis reaction) in activated sludge systems, the network diagram screened for species showing significant growth in response to oxygen perturbations and showing a positive correlation with the abundance of metabolites synthesised. Red lines indicate species that showed a significant positive correlation with the abundance of anabolic metabolites (Spearman correlation test; R>0, *P*<0.05).

However, for R03098 in lysine biosynthesis and R00031 in tyrosine metabolism, due to the limited species related to these two reactions revealed by the alignment results of the biological database, we cannot filter out significant information.

#### 3.3.2 Statistical Correlation between Microbial Genera and Differential Metabolites

To compensate for possible information loss in the BIO-Sankey analysis caused by the biological database matching process, we also used the Statistics-Sankey (STA-Sankey) analysis to further reveal the potential statistical correlation between microbial communities and metabolites in targeted metabolic pathways (Figure S54∼S66). We particularly focus on metabolic pathways where the function genes with significant differences under oxygen perturbations (CP and IP) are located, and the statistical association between microbial abundance and metabolite abundance, with the network map displaying very significant association results (T test, *P*<0.01).

For CP, the abundance of 10 genera is very significantly positively correlated with the abundance of metabolites, while only 3 genera show a very significant negative correlation (Figure 5a). For IP, the abundance of 16 genera is very significantly positively correlated with metabolite abundance, and 6 genera show a very significant negative correlation (Figure 5b). At the same time, *Labrys*, *Actinomyces*, and *Nitrosopumilus* all show a very significant positive correlation under both CP and IP. This suggests that under oxygen perturbations, these microbial groups may play a key role in the process of producing more amino acids in activated sludge.

**Figure 5.**
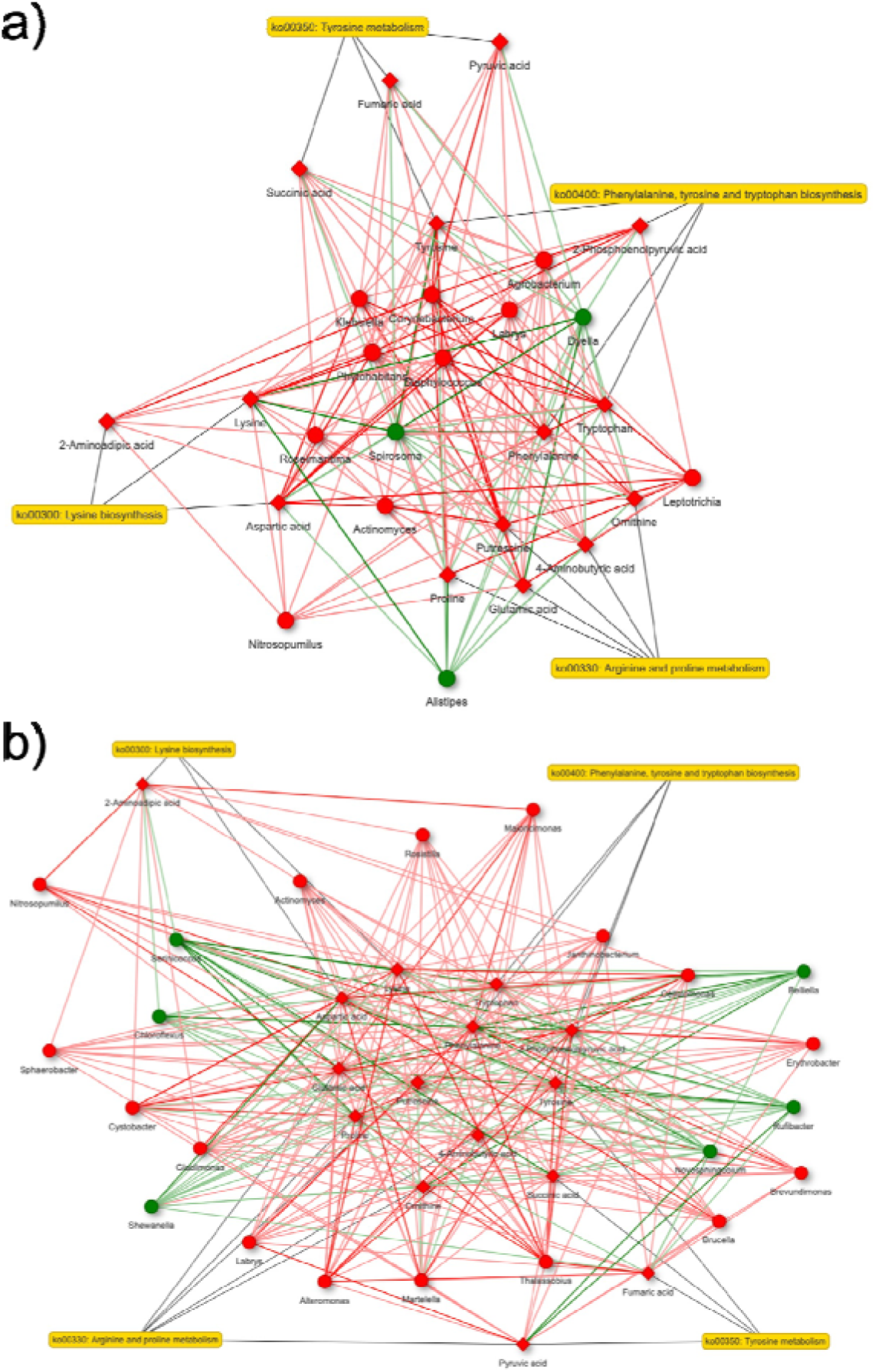
Network diagrams of statistically significant (T test, *P*<0.01) genera for metabolites related to arginine and proline metabolism (ko00330), phenylalanine, tyrosine, and tryptophan biosynthesis (ko00400), lysine biosynthesis (ko00300), and tyrosine metabolism (ko00350), and tyrosine metabolism (ko00350): (a) CP vs. CA; (b) IP vs. CA. Diamonds and nodes indicate associated metabolites and microorganisms, respectively. Red (or green) nodes indicate genera with significantly higher (or lower) oxygen perturbation. Red (or green) connecting lines indicate a positive (or negative) correlation between genera and metabolites.

In addition, we used the Spearman correlation method to perform correlation analysis between all detected intracellular metabolites and microbial species, finding that some genera are correlated with multiple metabolites (Figure S67, S68). This shows that in the process of oxygen perturbation promoting the synthesis of metabolites in the activated sludge system, there may be other mechanisms of response to microbial growth contributing to it.

## 4. DISCUSSION

In this study, we explored how oxygen perturbations affect microbial growth, thereby changing the abundance of functional genes and promoting the synthesis of metabolites such as amino acids. In the activated sludge system, microbes are often affected by the switching between aerobic and anaerobic conditions, which changes their growth rates. Different microbes have different requirements for the oxygen environment, such as aerobic microbes, anaerobic microbes, and facultative microbes. Even bacteria capable of aerobic respiration, due to the differences in oxygen affinity, will have different metabolism and growth rates. When microbial communities are long-term in stable aerobic, low oxygen, or anaerobic environments, microbial structure may tend to highlight the dominant species of a certain type^46^. When the oxygen environment dynamically changes, microbes need to adjust their structure and function to adapt to this change^47^. Especially when environmental changes are too drastic, it may affect the stability and efficiency of the system. However, the CP and IP used in this study did not have a negative impact on the stability of the system’s transformation of organic matter and nitrogen. On the contrary, the results of microbial diversity analysis showed that there was no significant difference in microbial diversity and richness under oxygen perturbation and non-perturbation conditions. The stability of the abundance of most bacterial species ensures the stability of biosystem functioning. However, there are still some species with particularly high or low oxygen demand or tolerance, and their overgrowth or massive death may lead to changes in their abundance in the population. This process often accompanies changes in the interaction relationship between communities, thus enabling the system to achieve a shift of metabolic function in response to the driving effect of environmental factors on it^48^.

In the activated sludge system, we found that changes in the oxygen environment affected the functional gene abundance, which further mapped onto the metabolic characteristics of the system. This phenomenon can be attributed to the influence of environmental parameters on the structure of microbial communities, thereby indirectly affecting enzyme production^49^. The abundance of functional genes not only reflects changes in community structure but also predicts the possibility of enzyme expression^50^. At the KEGG level 3 pathway level, we found that the gene abundance of energy metabolism and nucleotide metabolism can reflect the growth rate of biomass and its organic matter, while the gene abundance of amino acid metabolism and the metabolism of other amino acids reflects the synthesis of protein-like substances in biomass. At the same time, lipid metabolism is related to the humic-like substances in biomass. Although we previously started with redox regulation to explore the impact of oxygen perturbation on microbial system functions, this impact is not yet fully understood. In this study, we further explained the changes in microbial performance from the perspectives of community structure and gene function. In particular, under oxygen perturbations, the activity of metabolic pathways such as arginine and proline metabolism, lysine biosynthesis, phenylalanine, tyrosine, and tryptophan biosynthesis significantly increased. However, we also noted that in the process of using multi-omics joint analysis, in addition to the changes in gene abundance, factors such as enzyme activity, energy utilization efficiency, the abundance of cofactors and co-factors, and post-regulation of gene expression should be considered. These factors may produce effects individually or synergistically, and are worth further research^51,52^.

To identify microbes that play a decisive role in changes in metabolic pathway efficiency, we performed biological and statistical correlation analyses. We used the MetOrigin platform to compare the community information of the samples with databases, precisely screening out microbes that can participate in target metabolic reactions, and further locating microbial species that can grow significantly under oxygen perturbations and are positively correlated with the efficiency of proline and tryptophan synthesis reactions. Among them, *Mycolicibacterium madagascariense*, *Mycolicibacterium fortuitum*, and *Burkholderia pseudomallei* have the strongest correlations in the network. It is worth mentioning that the six species appearing in the network all belong to the genus *Mycolicibacterium*, which is known to growing rapidly in the environment and resistance to environmental changes^53^, and have the function of synthesizing steroidal compounds^54,55^. They not only play biological roles in processes such as cell membrane stabilization and cell proliferation but also provide a carbon source for other bacteria and participate in various cell signaling mechanisms^56,57^.

Through statistical correlation analysis for the microbial and metabolite abundance data, we can find correlated microbes that change consistently with metabolite abundance under oxygen perturbation. In the analysis, the *P*-value threshold was set to 0.01 to screen out microbial genera that were significantly positively correlated with the metabolites in the enriched metabolic pathways. We found that *Labrys*, *Actinomyces*, and *Nitrosopumilus* exist in the network under both CP and IP conditions. As a major genus of ammonia-oxidizing archaea, *Nitrosopumilus* usually has a higher oxygen affinity than ammonia-oxidizing bacteria^58^, and can carry out ammonia oxidation under anaerobic conditions^59^. Recent research shows that under anaerobic conditions, it can produce oxygen for ammonia oxidation, and can also reduce nitrite to gaseous nitrogen^60^. This may mean that in the activated sludge system under oxygen perturbations, it may play an important role in the nitrogen transformation process.

In summary, this study revealed the impact of microbial community growth under oxygen perturbations on amino acid metabolism. We aligned functional genes with metabolic features to determine the impact of microbial community changes on system performance; and screened for microbes that may function under oxygen perturbations through biological and statistical correlation analysis, providing directions for exploring the microbial response to oxygen perturbations in activated sludge system.

## ASSOCIATED CONTENT

### Supporting Information

Detailed materials and methods, microbial diversity, microbial community structure, 3D-EEM spectral characteristics, functional genes, metabolomics analysis, BIO-Sankey network, and STA-Sankey network (PDF) are described in the supporting information.

## ACKNOWLEDGMENT

We thank Nikki Freed for the assistance with metagenomic sequencing, and Saras Green and Alastair Harris for their help with the metabolomics analysis. We also thank Watercare Services Limited for providing the activated sludge culture from the Mangere Wastewater Treatment Plant. The authors acknowledge the use of New Zealand eScience Infrastructure (NeSI) high performance computing facilities, consulting support and/or training services as part of this research. The authors also acknowledge the Centre for eResearch at the University of Auckland for their help in facilitating this research.

