## Supplementary for "Investigating Potential Microbial Contributors to Enhanced Metabolite Production under Oxygen Perturbations: An Integrated Metagenomic and Metabolomic Approach"

### for

*Xueyang Zhou[1,2], Bharat Manna[1,2], Boyu Lyu[1,2], and Naresh Singhal[1,2]* *
[1]Department of Civil and Environmental Engineering, University of Auckland, Auckland 1010, New Zealand
[2]Water Research Centre, University of Auckland, Auckland 1010, New Zealand

#### S1. Methods

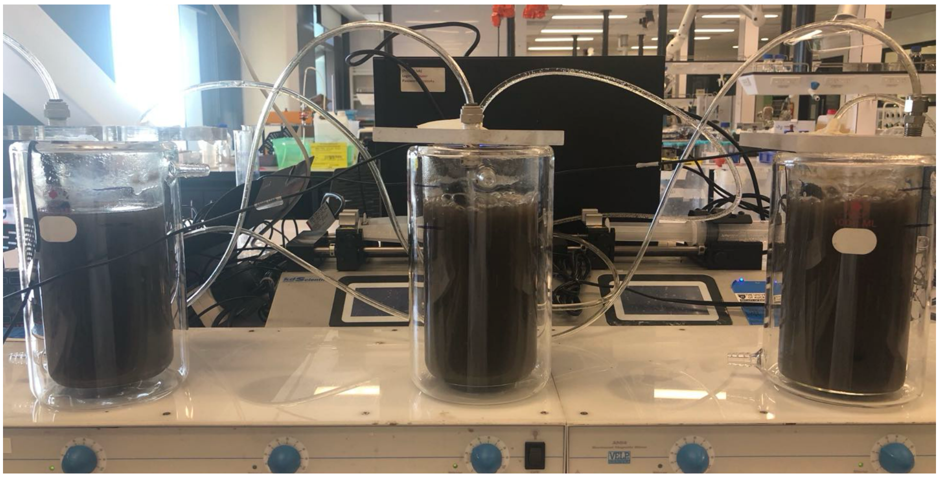

**Figure S1.** The laboratory scale bioreactors with activated sludge

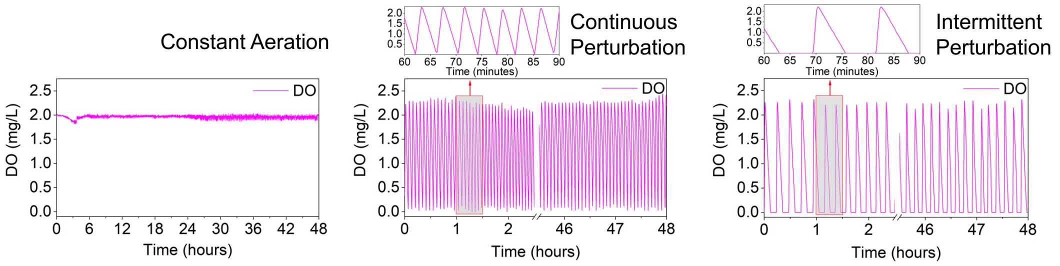


**Figure S2.** Different aeration patterns including constant aeration, continuous perturbation, and intermittent perturbation

#### S2. Results

##### S2.1. Microbial diversity analysis

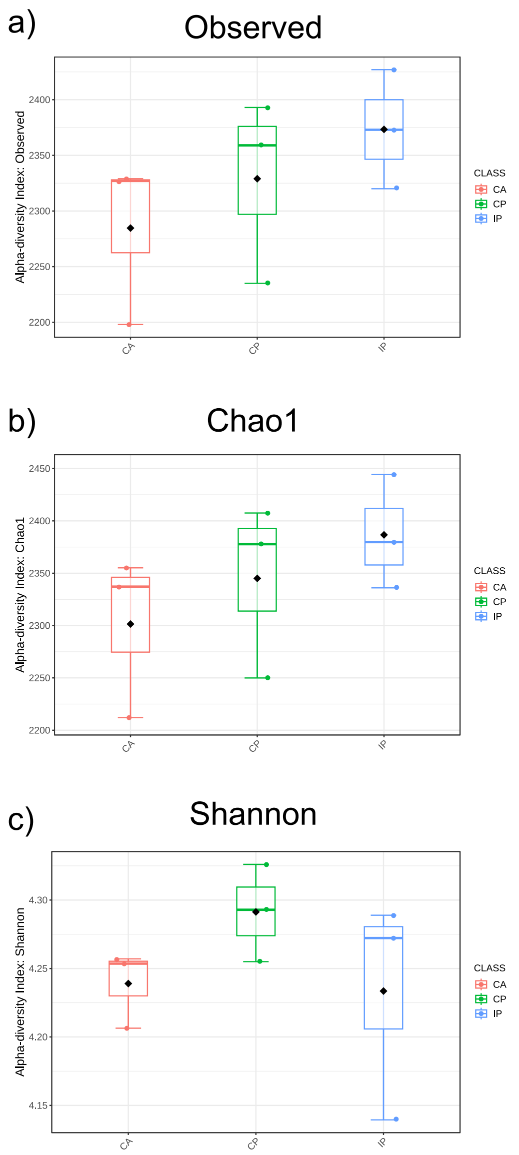


**Figure S3.** Box plots of alpha diversity indices including richness, Chao1, and Shannon (biological triplicates; n=3)

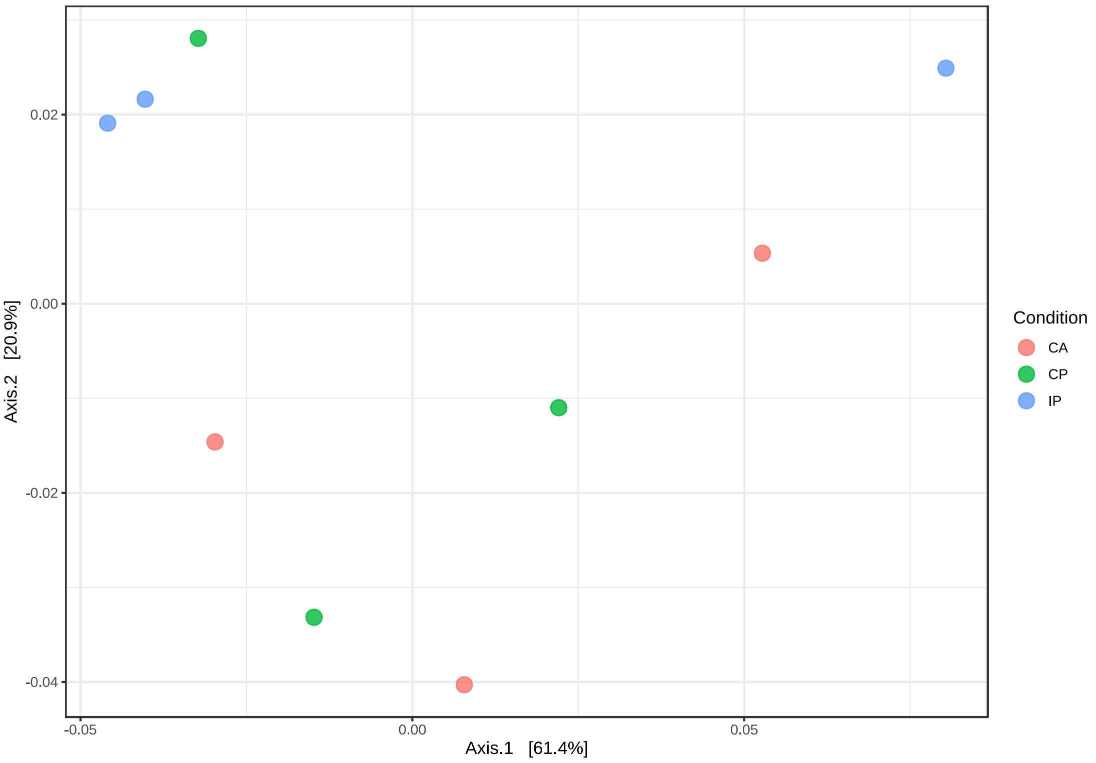


**Figure S4.** The principal coordinate analysis (PCoA) based on Bray–Curtis distance (biological triplicates; n=3)

#### S2.2. Microbial community structure analysis

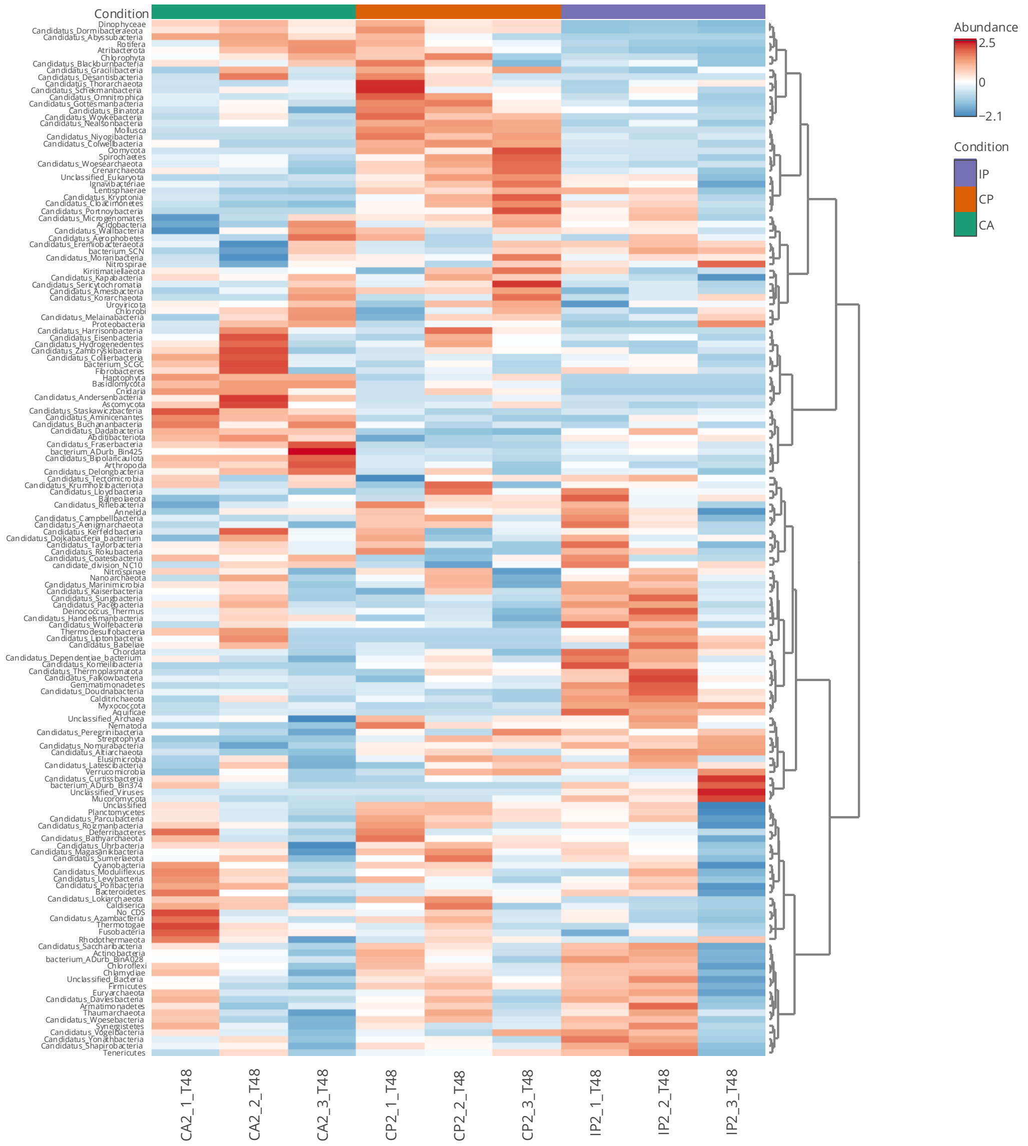


**Figure S5.** The heatmap of taxonomy analysis at the phylum level. The data is filtering based on the ‘Low count filter’ to a minimum count of 4 and 20% prevalence in samples and the ‘percentage to remove’ option under ‘Low variance filter’ set to 10% based on the interquantile range; and normalized by total sum scaling. Row clustering according to ‘Ward’ (biological triplicates; n=3)

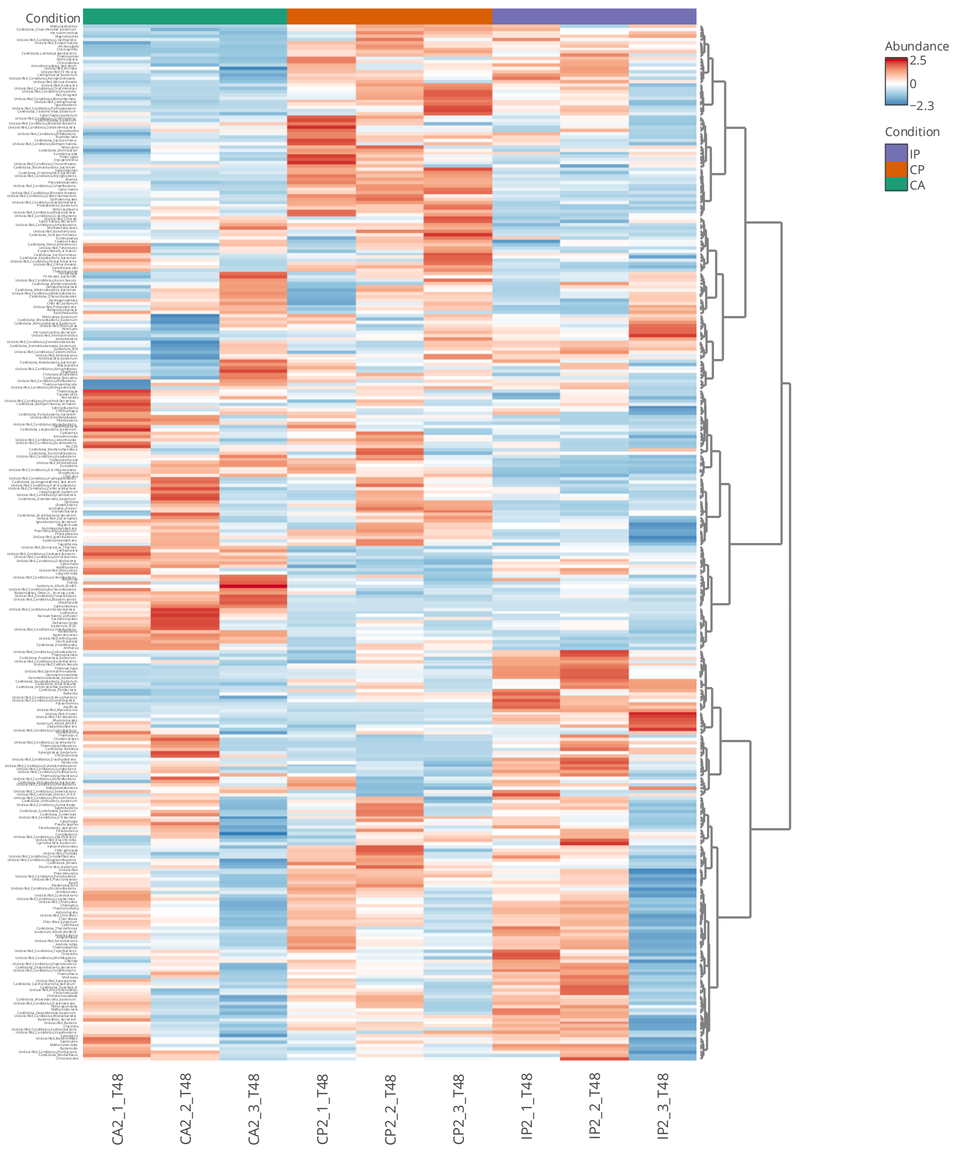


**Figure S6.** The heatmap of taxonomy analysis at the class level. The data is filtering based on the 'Low count filter' to a minimum count of 4 and 20% prevalence in samples and the 'percentage to remove' option under 'Low variance filter' set to 10% based on the interquantile range; and normalized by total sum scaling. Row clustering according to 'Ward' (biological triplicates; n=3)

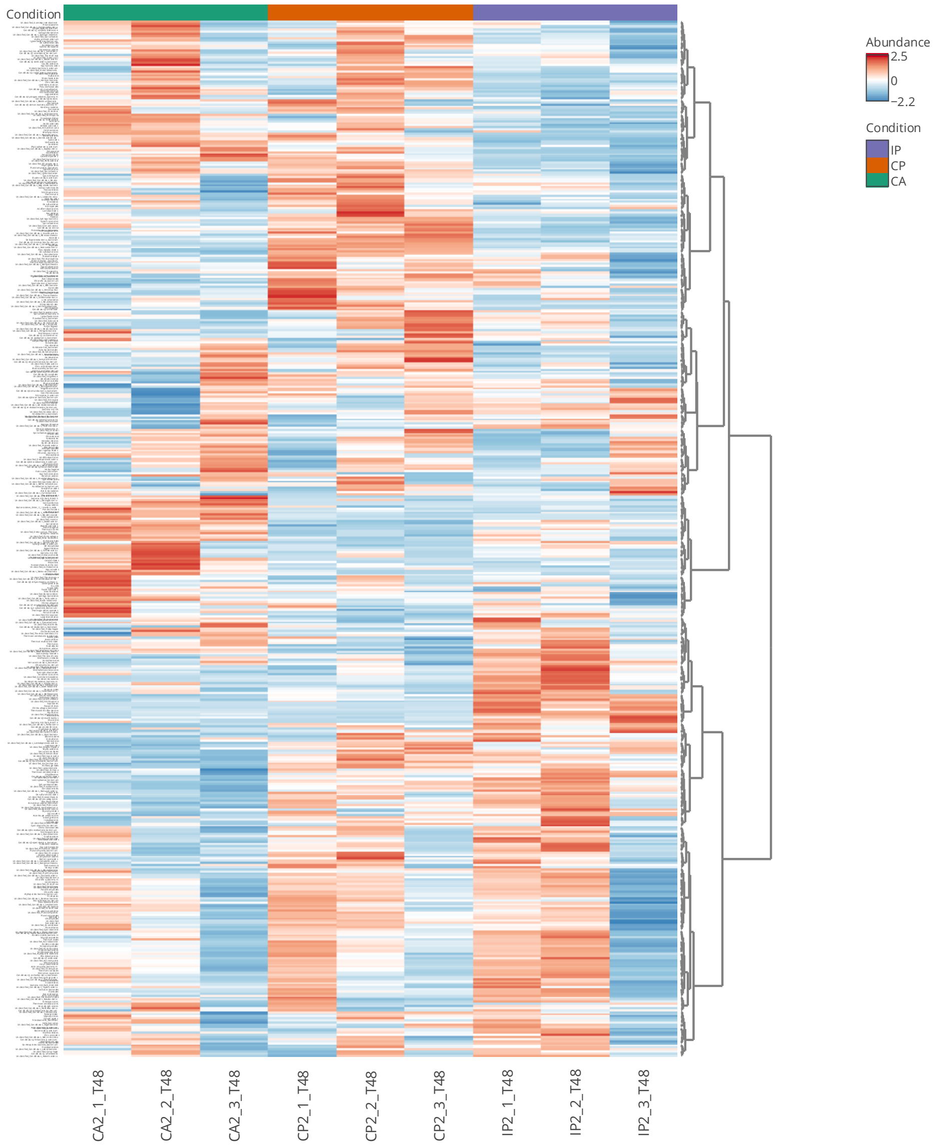


**Figure S7.** The heatmap of taxonomy analysis at the order level. The data is filtering based on the 'Low count filter' to a minimum count of 4 and 20% prevalence in samples and the 'percentage to remove' option under 'Low variance filter' set to 10% based on the interquantile range; and normalized by total sum scaling. Row clustering according to 'Ward' (biological triplicates; n=3)

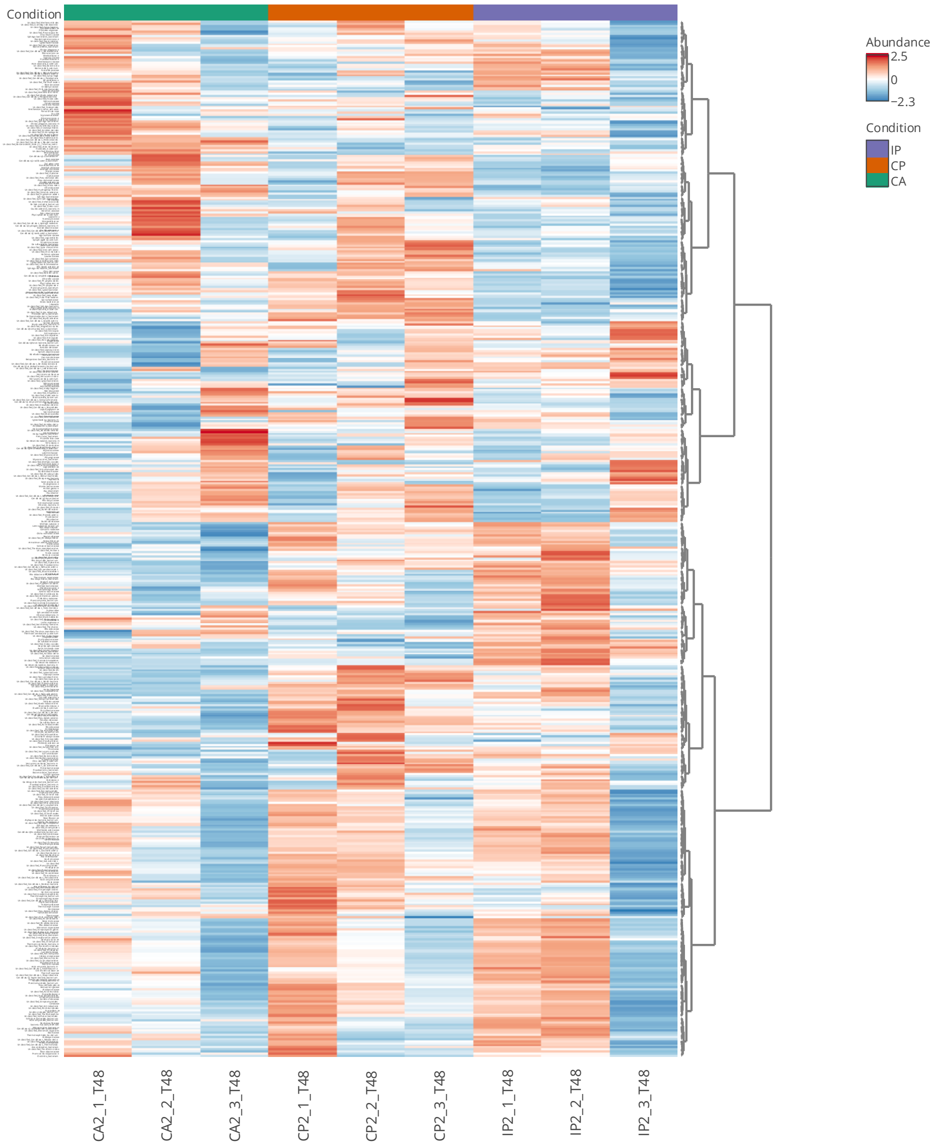


**Figure S8.** The heatmap of taxonomy analysis at the family level. The data is filtering based on the 'Low count filter' to a minimum count of 4 and 20% prevalence in samples and the 'percentage to remove' option under 'Low variance filter' set to 10% based on the interquantile range; and normalized by total sum scaling. Row clustering according to 'Ward' (biological triplicates; n=3)

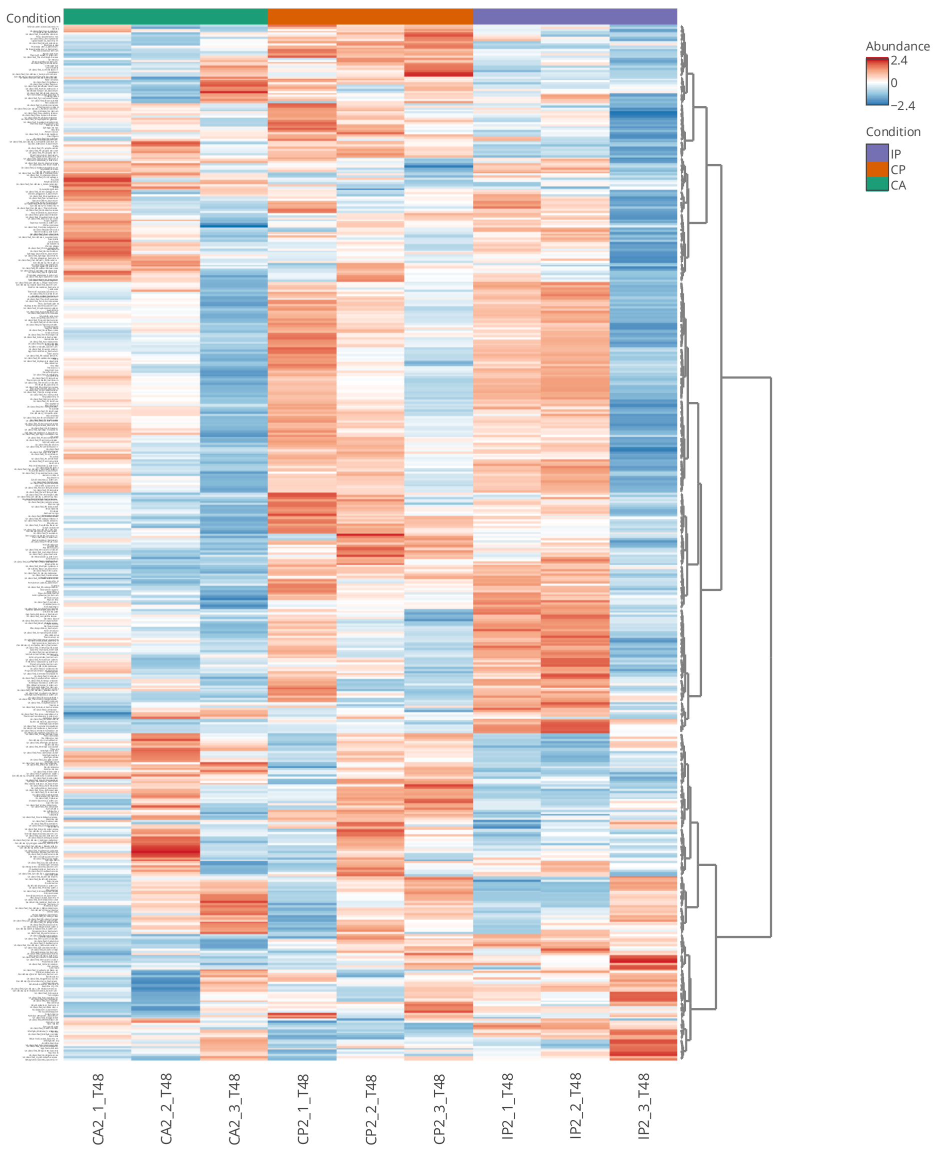


**Figure S9.** The heatmap of taxonomy analysis at the genus level. The data is filtering based on the 'Low count filter' to a minimum count of 4 and 20% prevalence in samples and the 'percentage to remove' option under 'Low variance filter' set to 10% based on the interquantile range; and normalized by total sum scaling. Row clustering according to 'Ward' (biological triplicates; n=3)

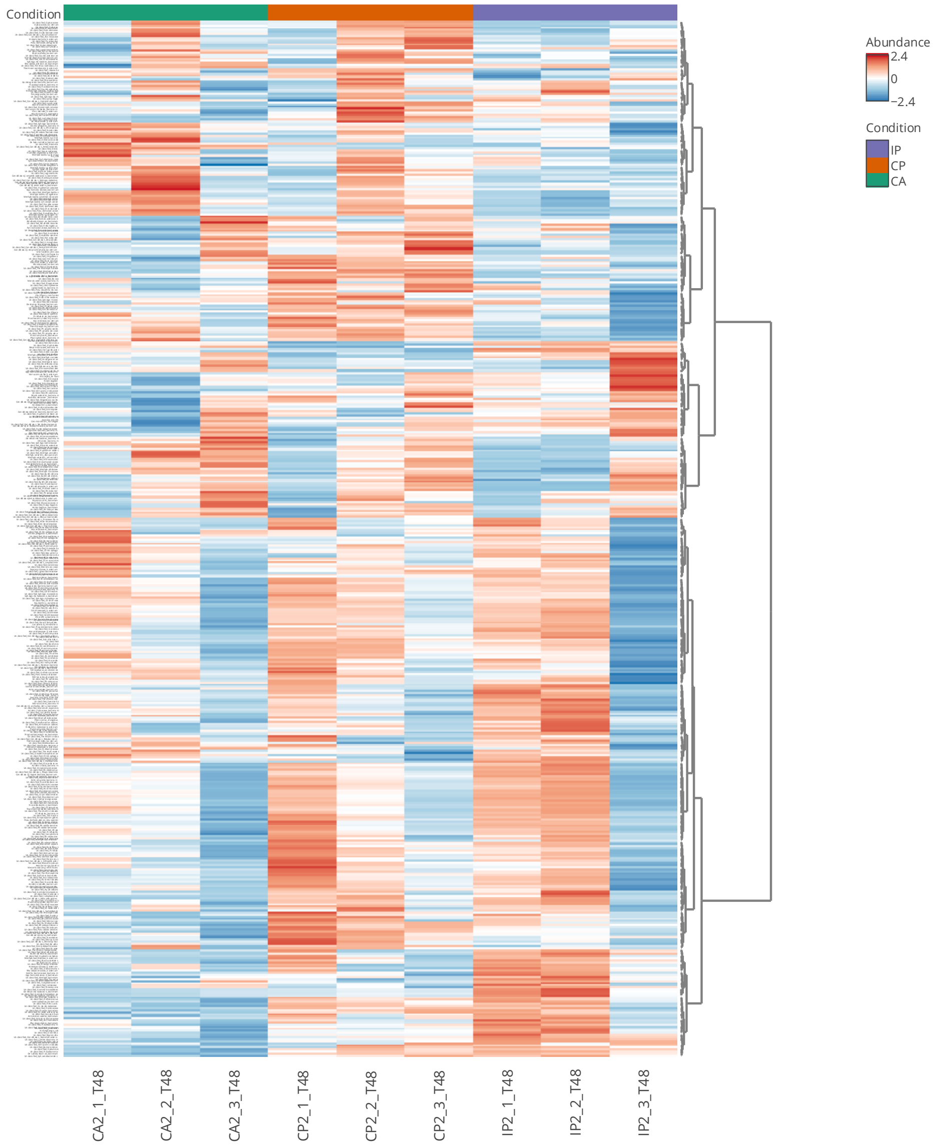


**Figure S10.** The heatmap of taxonomy analysis at the species level. The data is filtering based on the 'Low count filter' to a minimum count of 4 and 20% prevalence in samples and the 'percentage to remove' option under 'Low variance filter' set to 10% based on the interquantile range; and normalized by total sum scaling. Row clustering according to 'Ward' (biological triplicates; n=3)

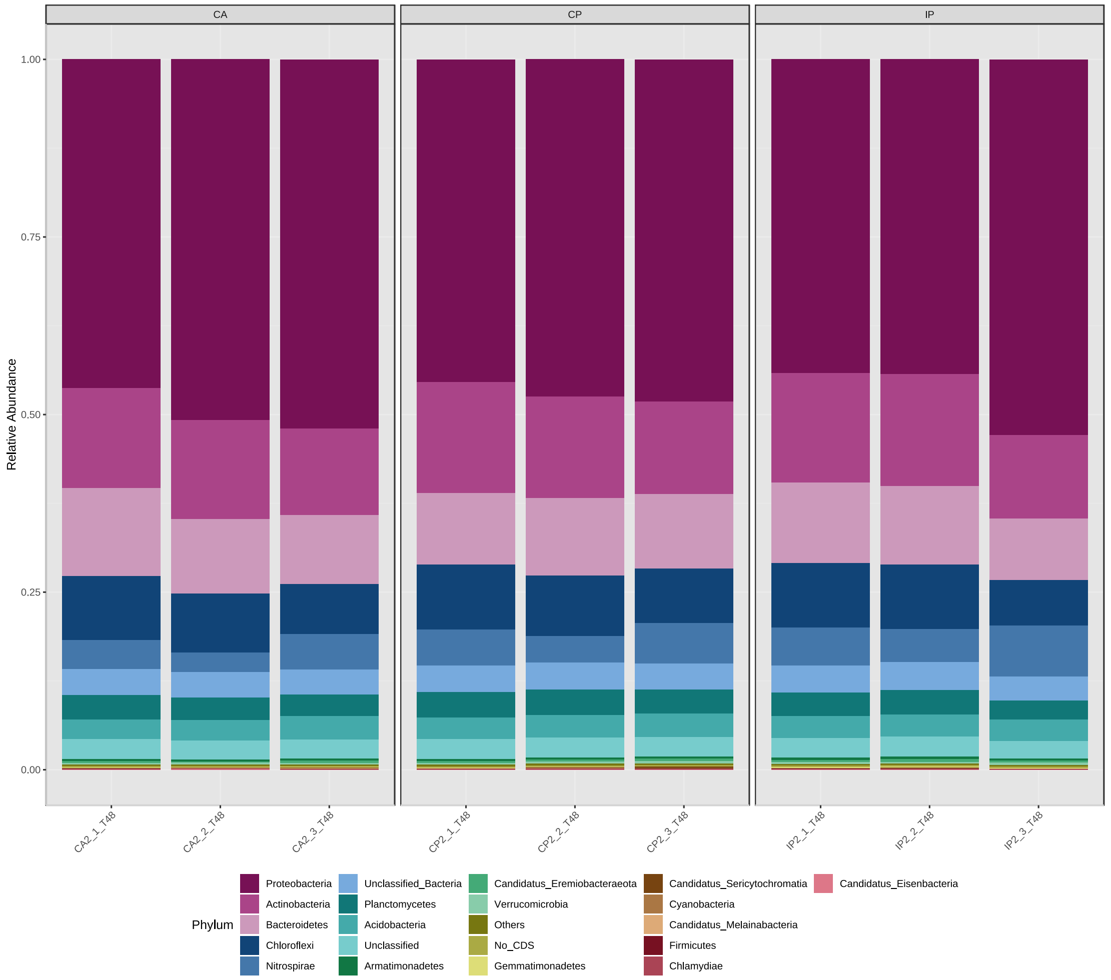


**Figure S11.** Relative abundance of microbial taxonomic groups at the phylum (top 20 phyla and others) level

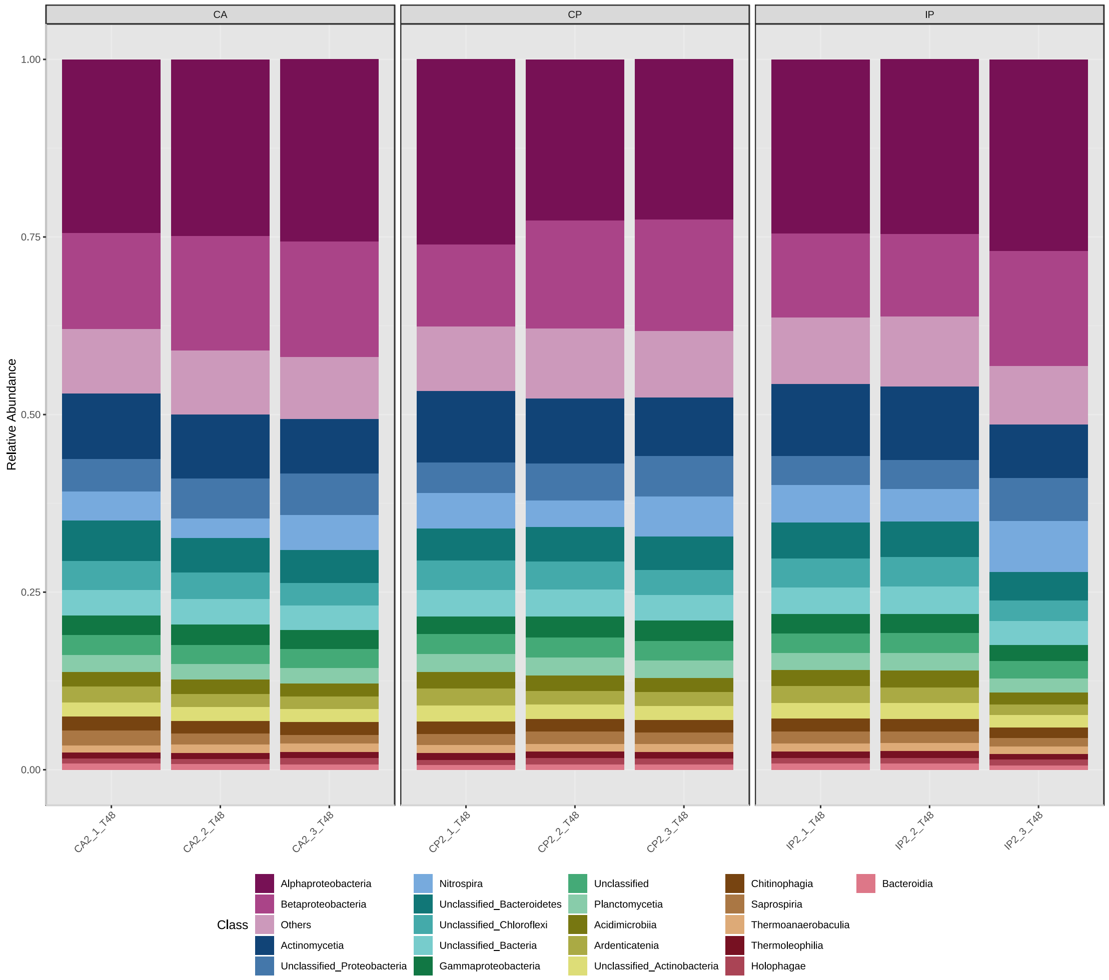


**Figure S12.** Relative abundance of microbial taxonomic groups at the class (top 20 classes and others) level

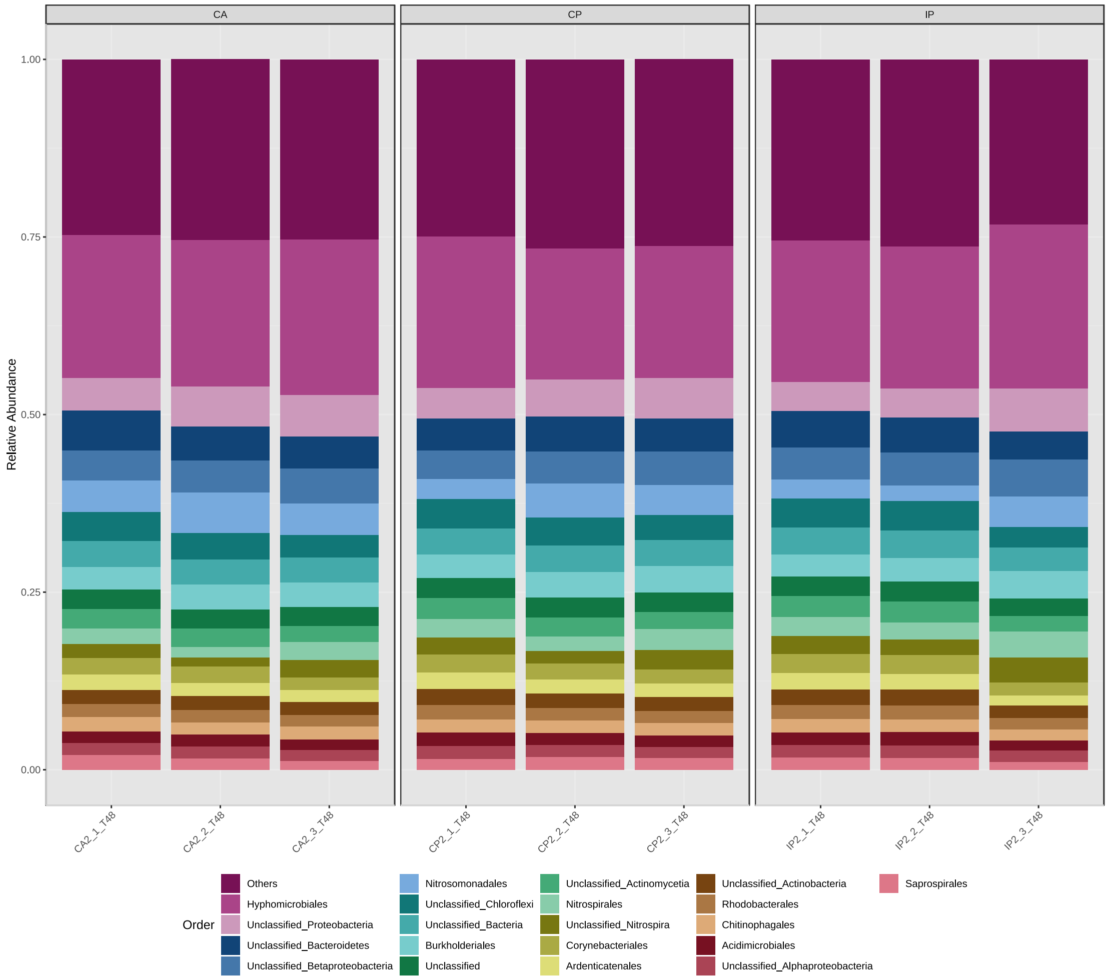


**Figure S13.** Relative abundance of microbial taxonomic groups at the order (top 20 orders and others) level

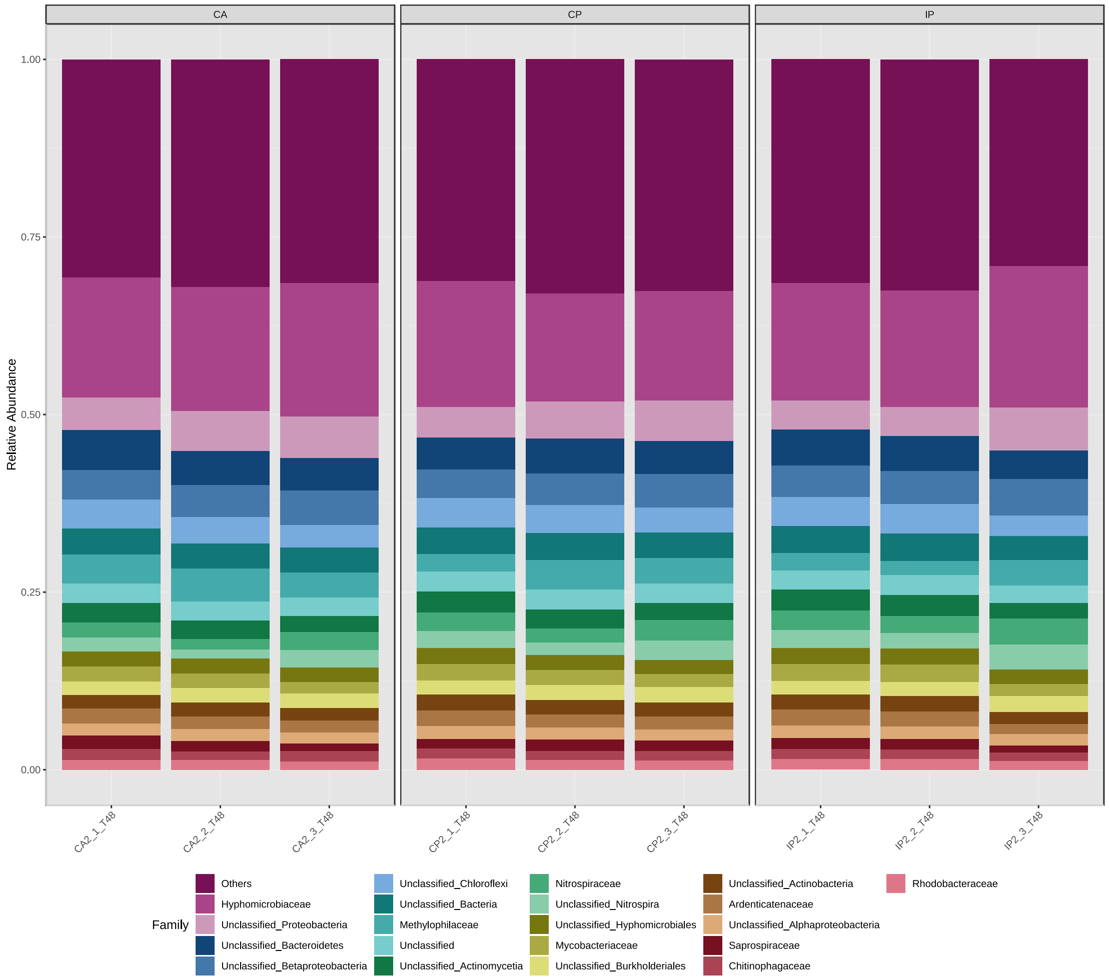


**Figure S14.** Relative abundance of microbial taxonomic groups at the family (top 20 families and others) level

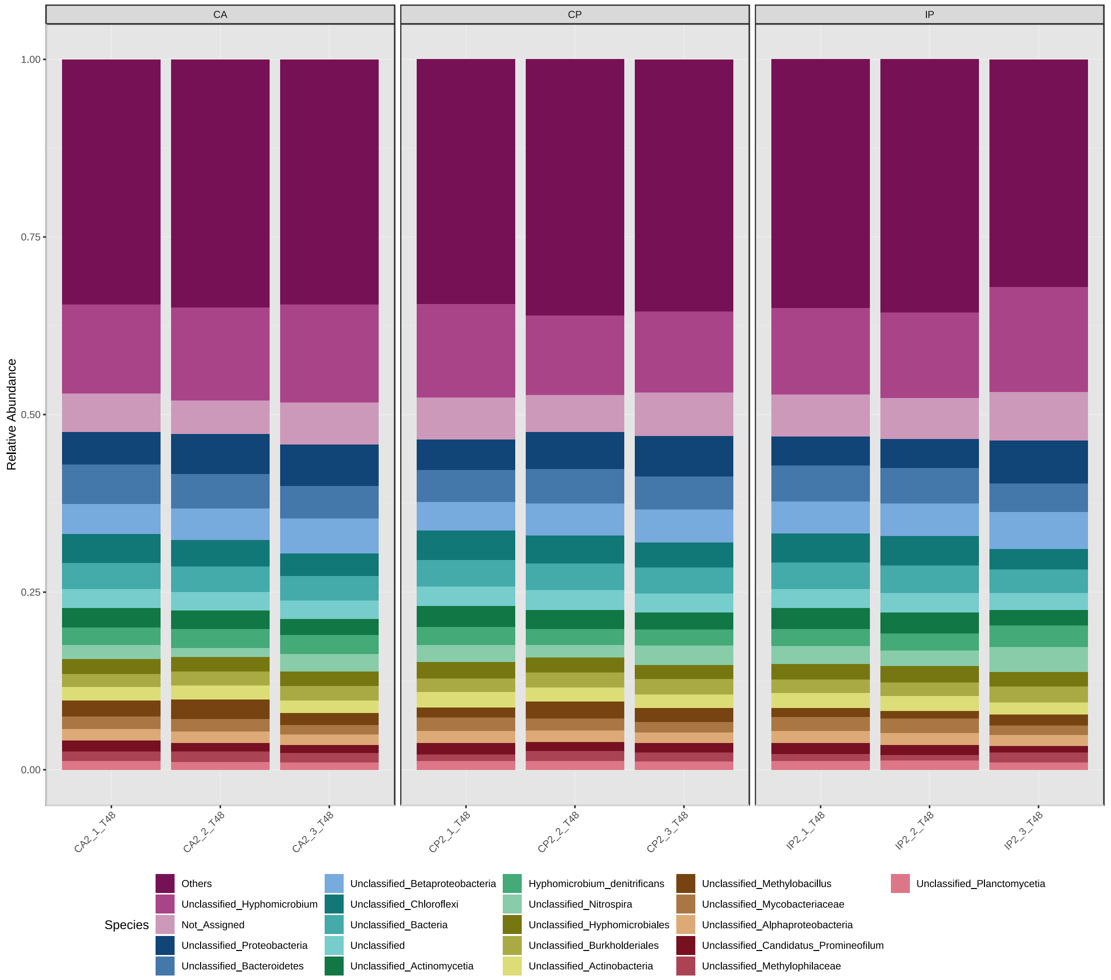


**Figure S15.** Relative abundance of microbial taxonomic groups at the species (top 20 species and others) level

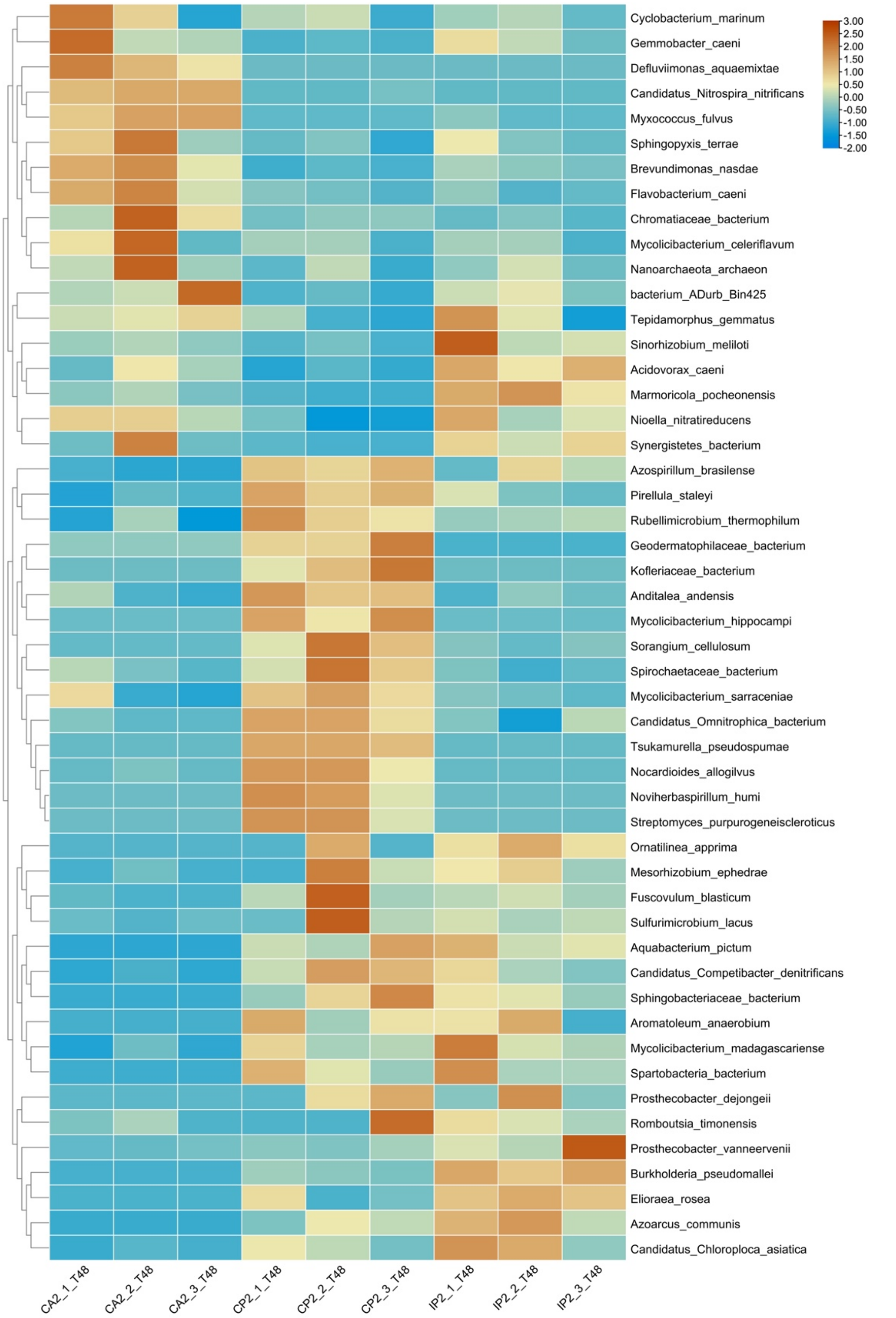


**Figure S16.** Classification of species annotated with significant differences (metagenomeSeq_0-inflated test, *P*<0.01) in abundance among different aeration conditions. Row clustering according to 'Ward'

##### S2.3. Analysis of organic matter in biomass

**Table S1.** The properties of 4 components identified by Openfluor

| Component | Excitation maximum | Emission maximum | Assignment | References |
|---|---|---|---|---|
| Component 1 | 291 nm | 339 nm | Protein-like compound | (Han et al., 2022) |
| Component 2 | 369 nm | 467 nm | Humic-like compound | (Gao & Guéguen, 2017) |
| Component 3 | 468 nm | 534 nm | No matches | |
| Component 4 | 324 nm | 418 nm | Humic-like compound | (Chen et al., 2018) |

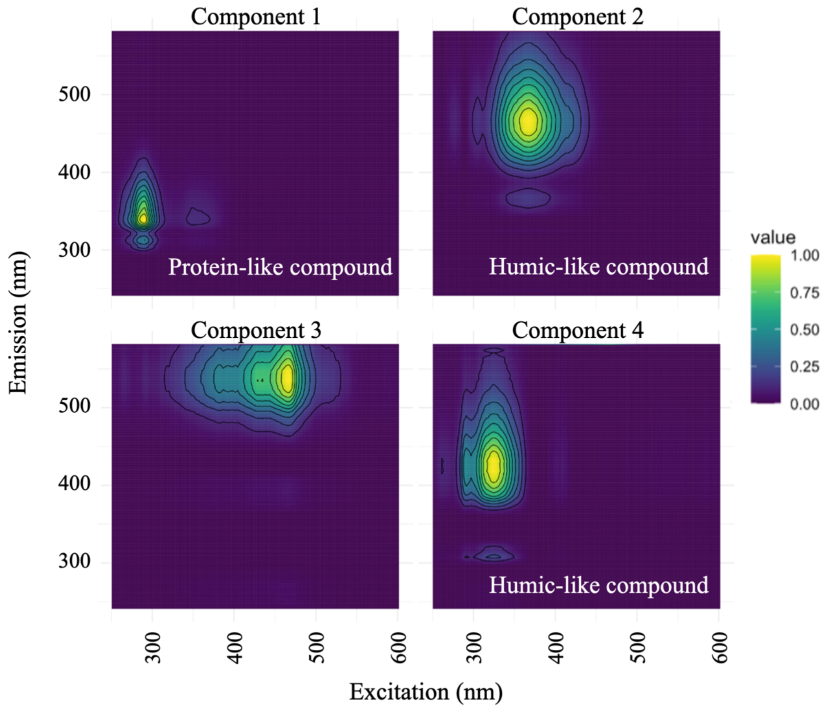


**Figure S17.** fDOM component spectral characteristics were identified by the PARAFAC model

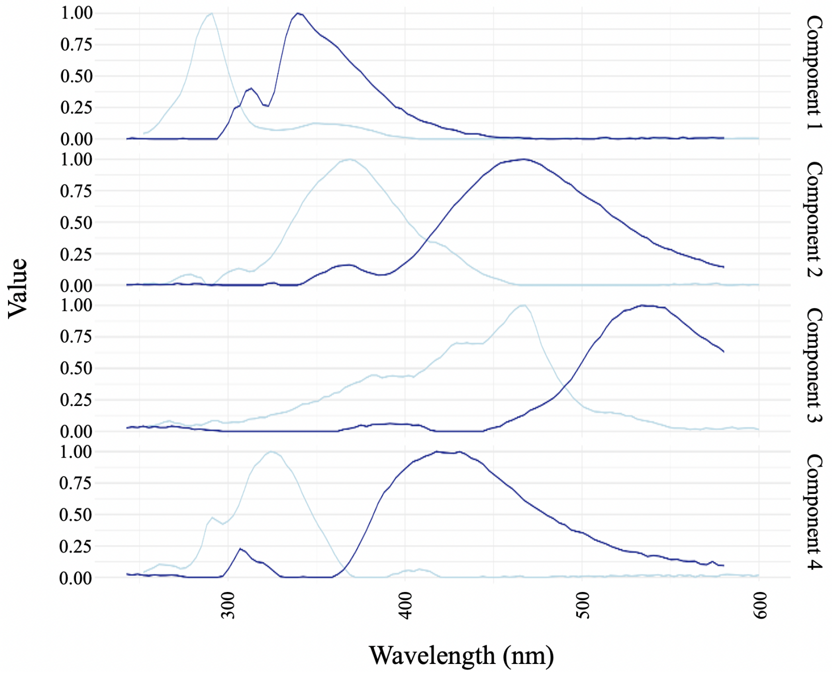


**Figure S18.** Excitation (light Blue) and emission (dark blue) curves of the fDOM components

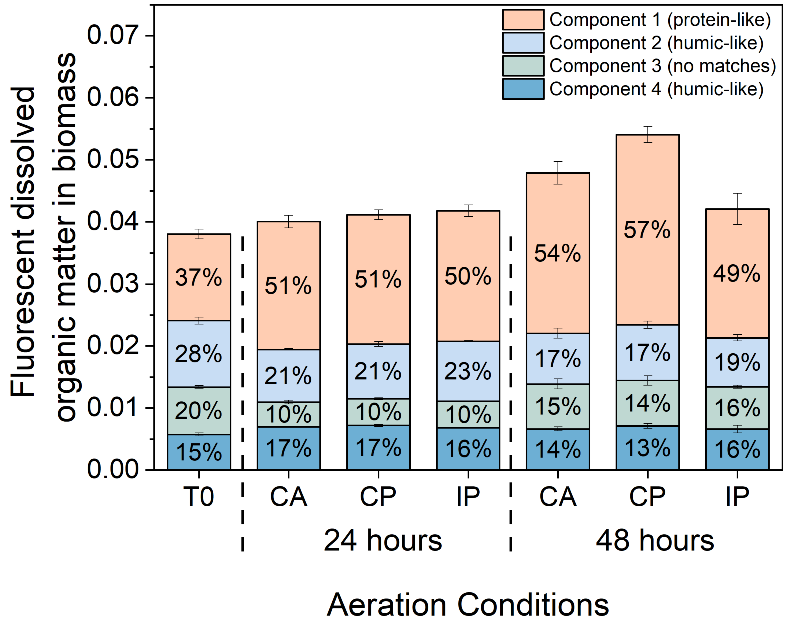


**Figure S19.** Fluorescent dissolved organic matter contents in biomass samples from activated sludge system exposed to different aeration conditions. Error bars represent standard deviations (biological triplicates; n=3).

#### S2.4. Functional genes analysis

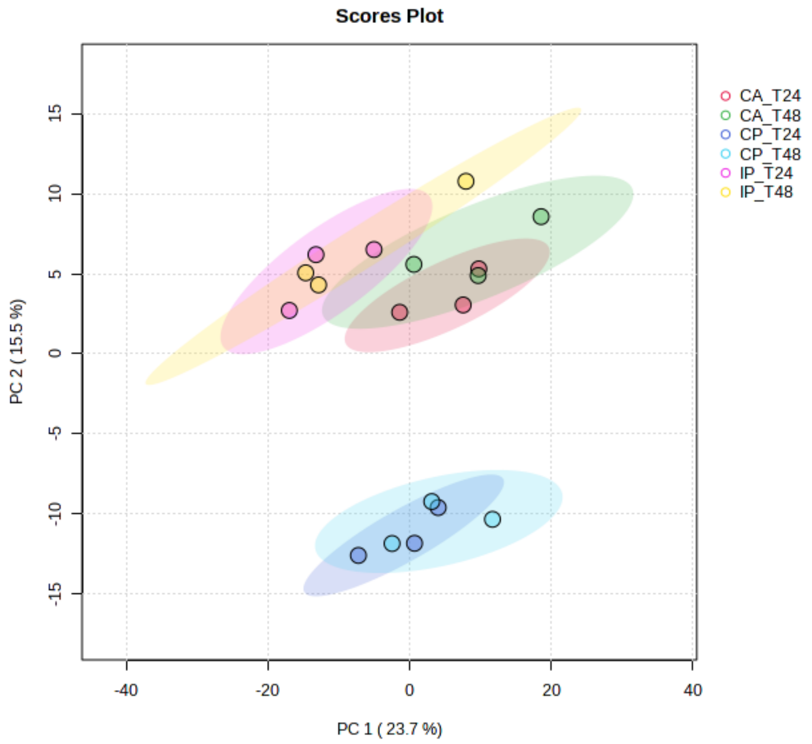


**Figure S20.** Principal component analysis (PCA) based on microbiota gene abundance, with elliptical boundaries representing 95% confidence regions

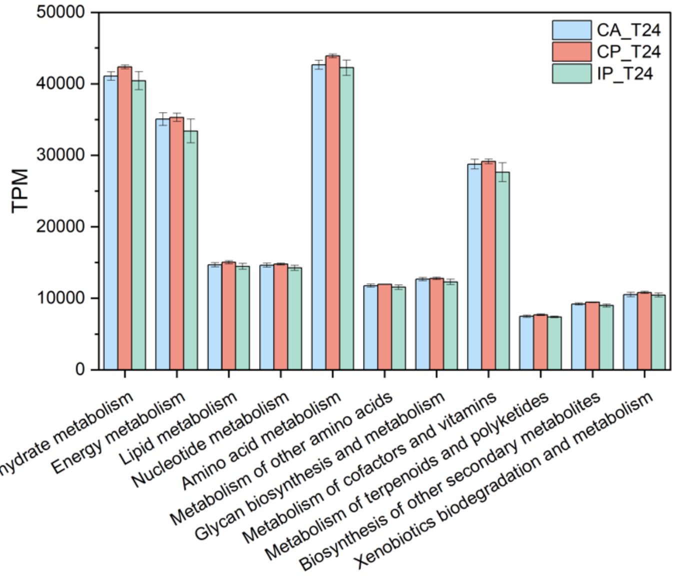


**Figure S21.** The KEGG level 2 functions of activated sludge samples exposed to different aeration conditions after 24 hours. The error bars represent standard deviations (biological triplicates; n=3)

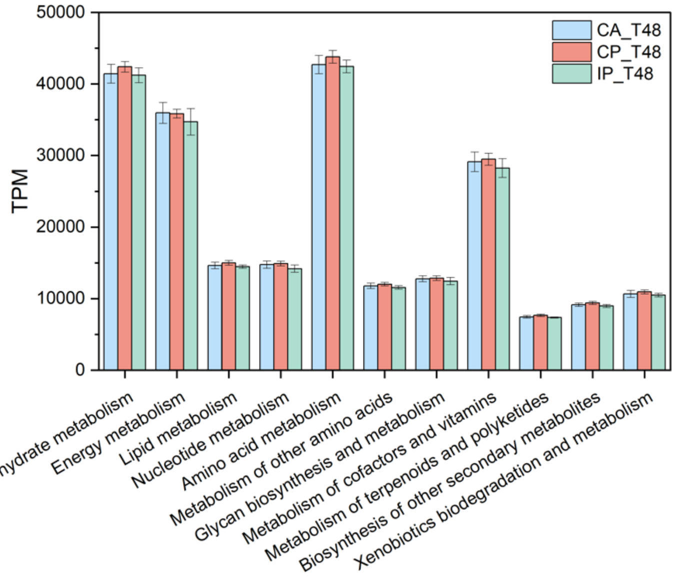


**Figure S22.** The KEGG level 2 functions of activated sludge samples exposed to different aeration conditions after 48 hours. The error bars represent standard deviations (biological triplicates; n=3)

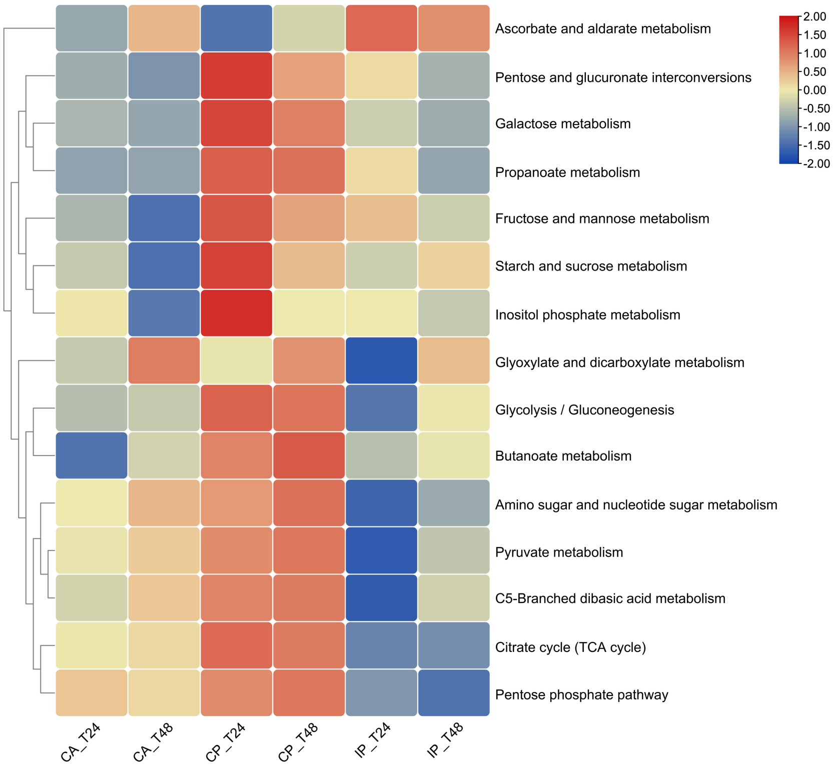


**Figure S23.** Gene abundance in the KEGG level 3 pathways under carbohydrate metabolism. The heatmap shows the average value of the same condition. Row clustering according to 'Ward' (biological triplicates; n=3)

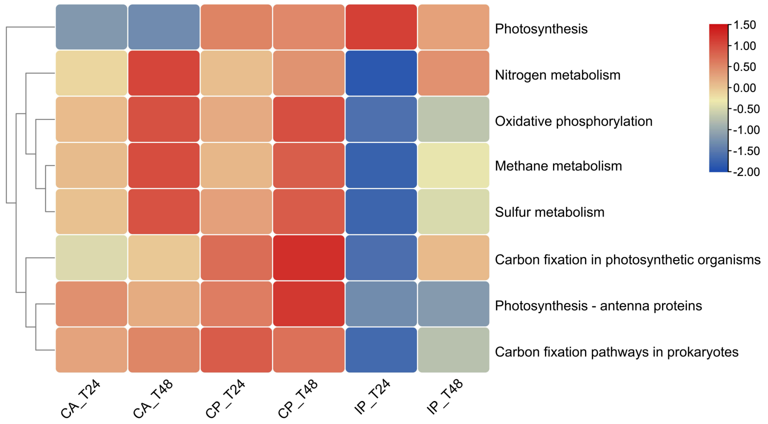


**Figure S24.** Gene abundance in the KEGG level 3 pathways under energy metabolism. The heatmap shows the average value of the same condition. Row clustering according to 'Ward' (biological triplicates; n=3)

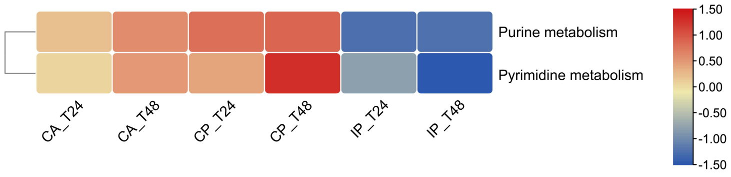


**Figure S25.** Gene abundance in the KEGG level 3 pathways under nucleotide metabolism. The heatmap shows the average value of the same condition. Row clustering according to 'Ward' (biological triplicates; n=3)

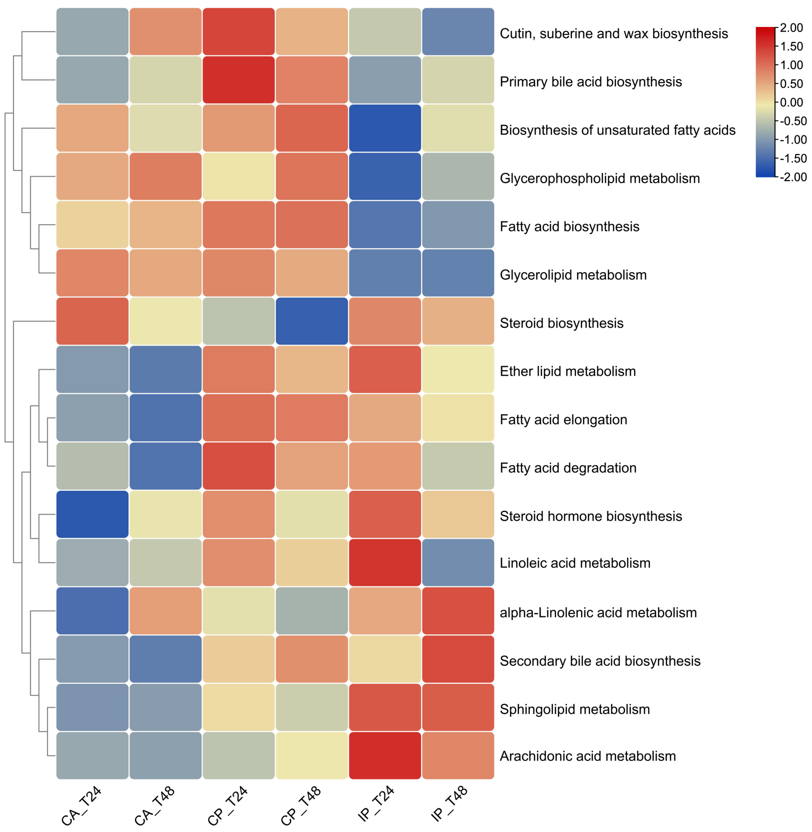


**Figure S26.** Gene abundance in the KEGG level 3 pathways under lipid metabolism. The heatmap shows the average value of the same condition. Row clustering according to 'Ward' (biological triplicates; n=3)

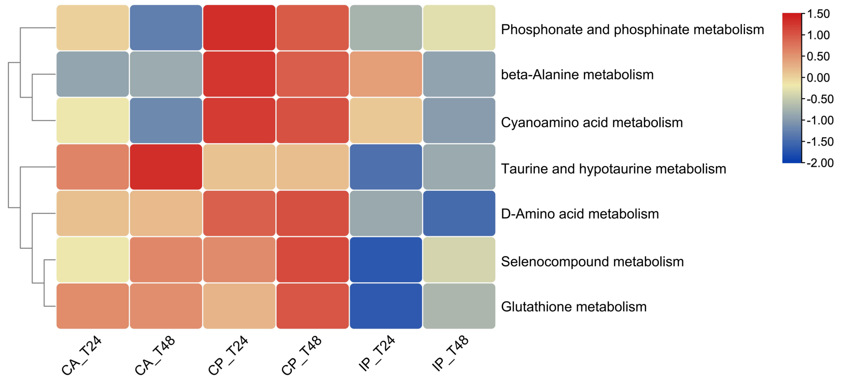


**Figure S27.** Gene abundance in the KEGG level 3 pathways under the metabolism of other amino acids. The heatmap shows the average value of the same condition. Row clustering according to 'Ward' (biological triplicates; n=3)

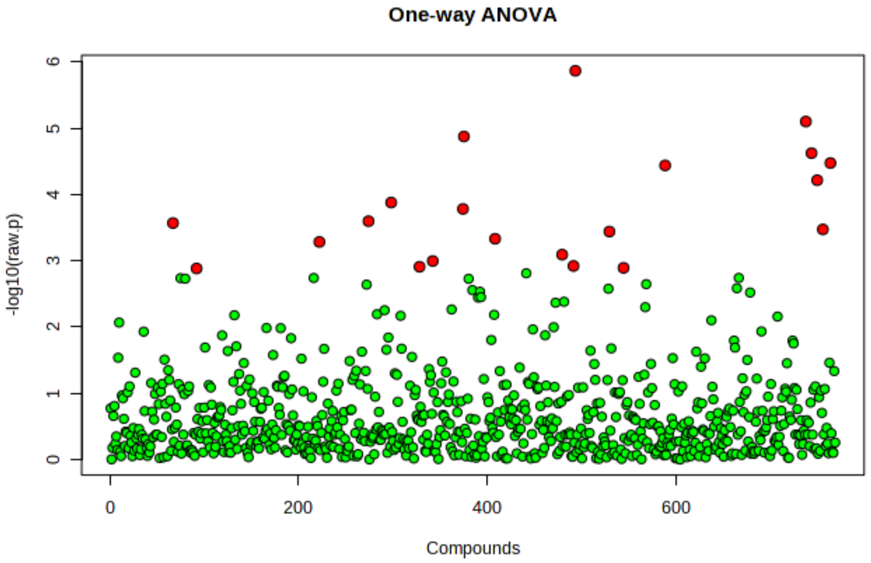


**Figure S28.** Differential abundance analysis of genes on carbohydrate metabolism by ANOVA analysis (21 significant differences out of 761 genes, $P<0.05$)

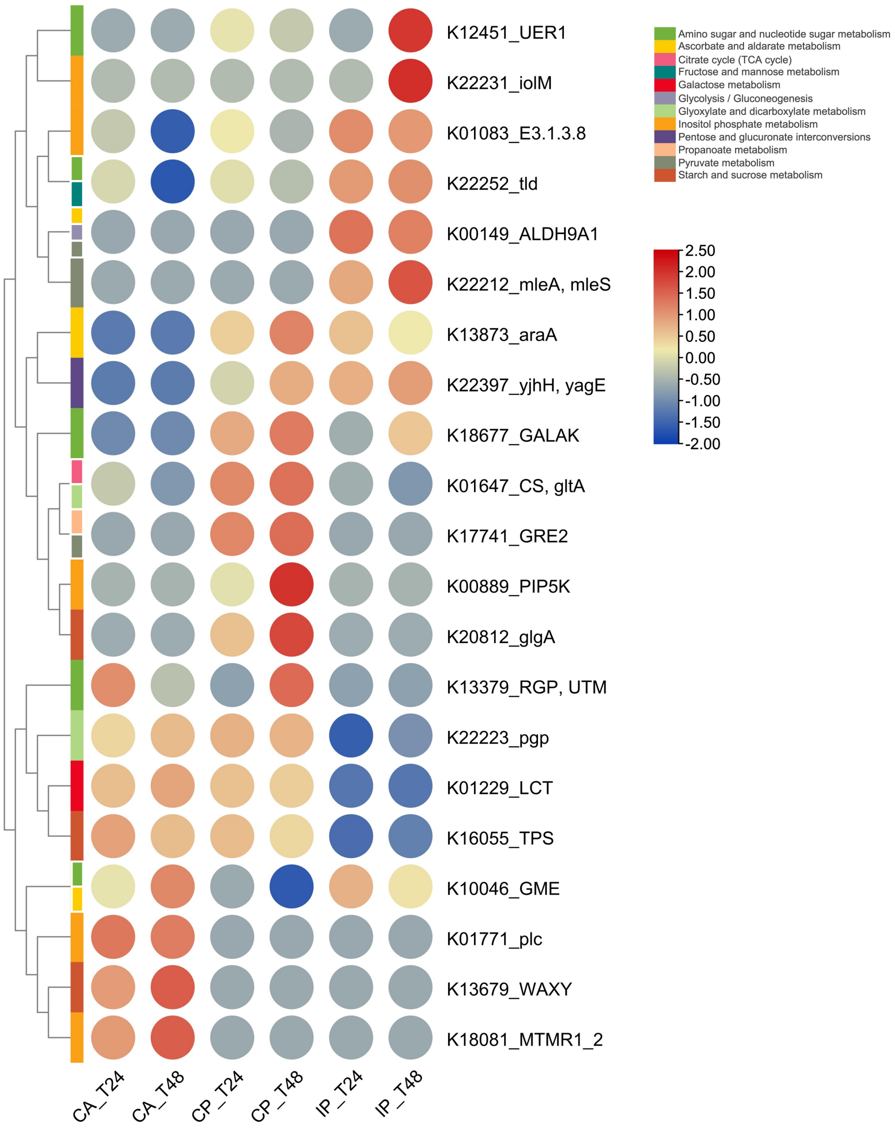


**Figure S29.** Differential abundance of genes on carbohydrate metabolism under different aeration conditions. The heatmap shows the average value of the same condition. Row clustering according to 'Ward' (biological triplicates; n=3)

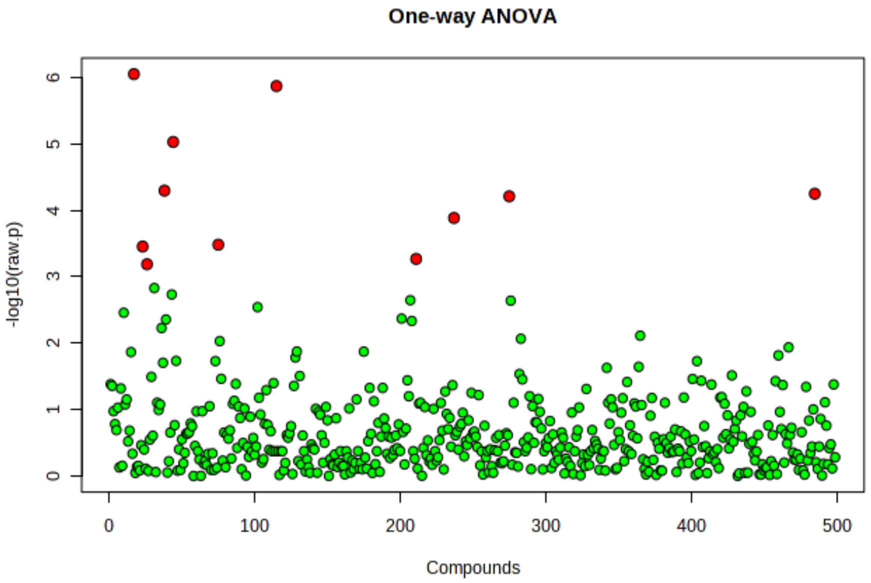


**Figure S30.** Differential abundance analysis of genes on energy metabolism by ANOVA analysis (11 significant differences out of 499 genes, *P*<0.05)

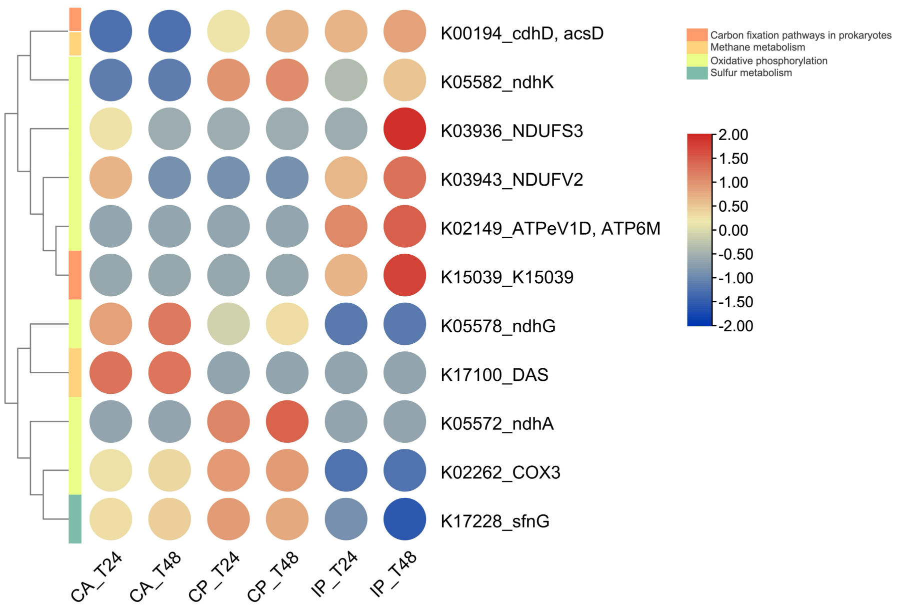


**Figure S31.** Differential abundance of genes on energy metabolism under different aeration conditions. The heatmap shows the average value of the same condition. Row clustering according to ‘Ward’ (biological triplicates; n=3)

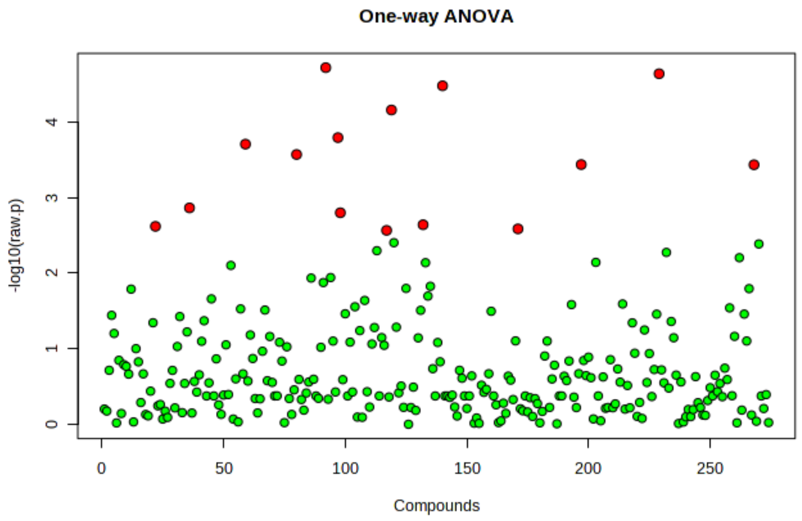


**Figure S32.** Differential abundance analysis of genes on lipid metabolism by ANOVA analysis (15 significant differences out of 274 genes, $P<0.05$)

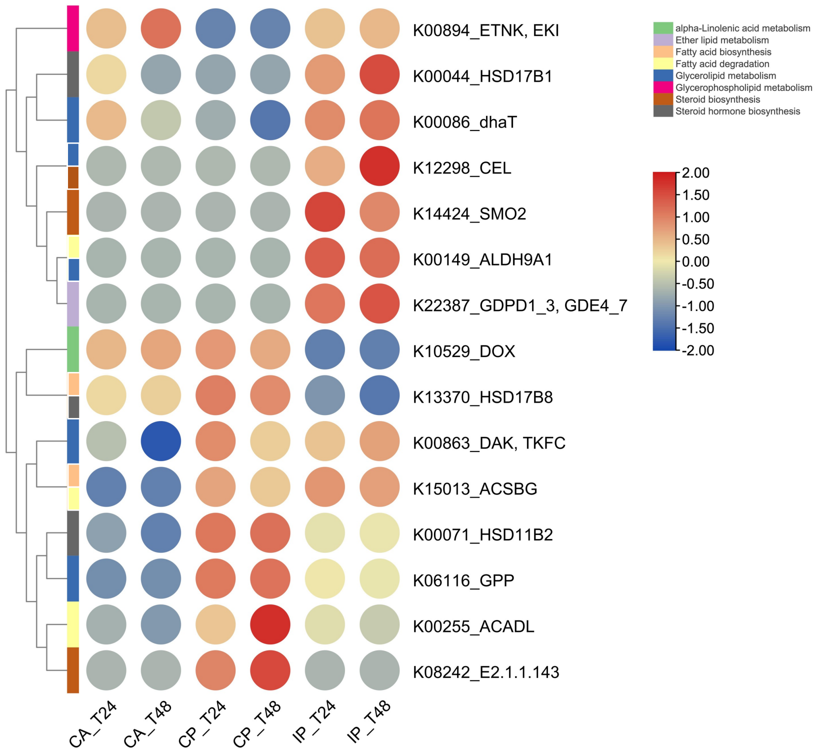


**Figure S33.** Differential abundance of genes on lipid metabolism under different aeration conditions. The heatmap shows the average value of the same condition. Row clustering according to 'Ward' (biological triplicates; n=3)

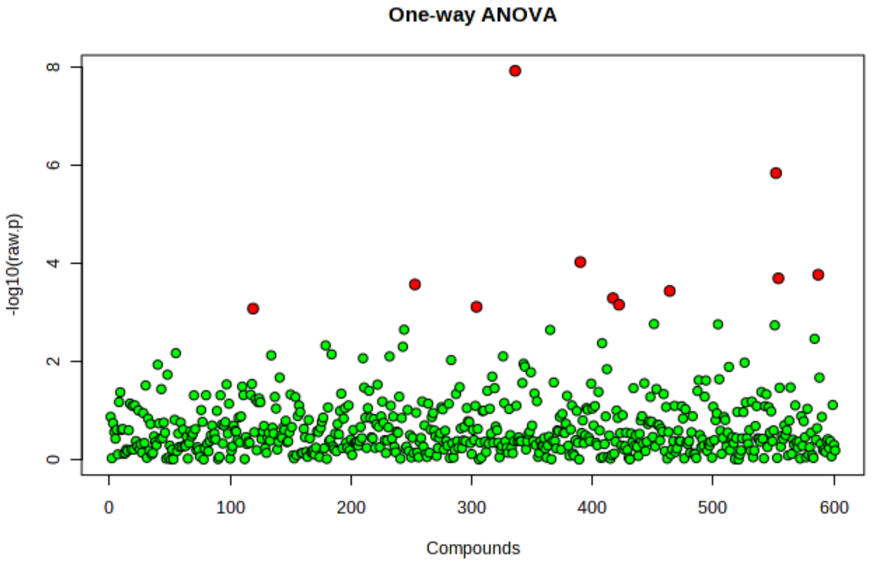


**Figure S34.** Differential abundance analysis of genes on amino acid metabolism by ANOVA analysis (11 significant differences out of 601 genes, $P<0.05$)

#### S2.5. Metabolomics analysis

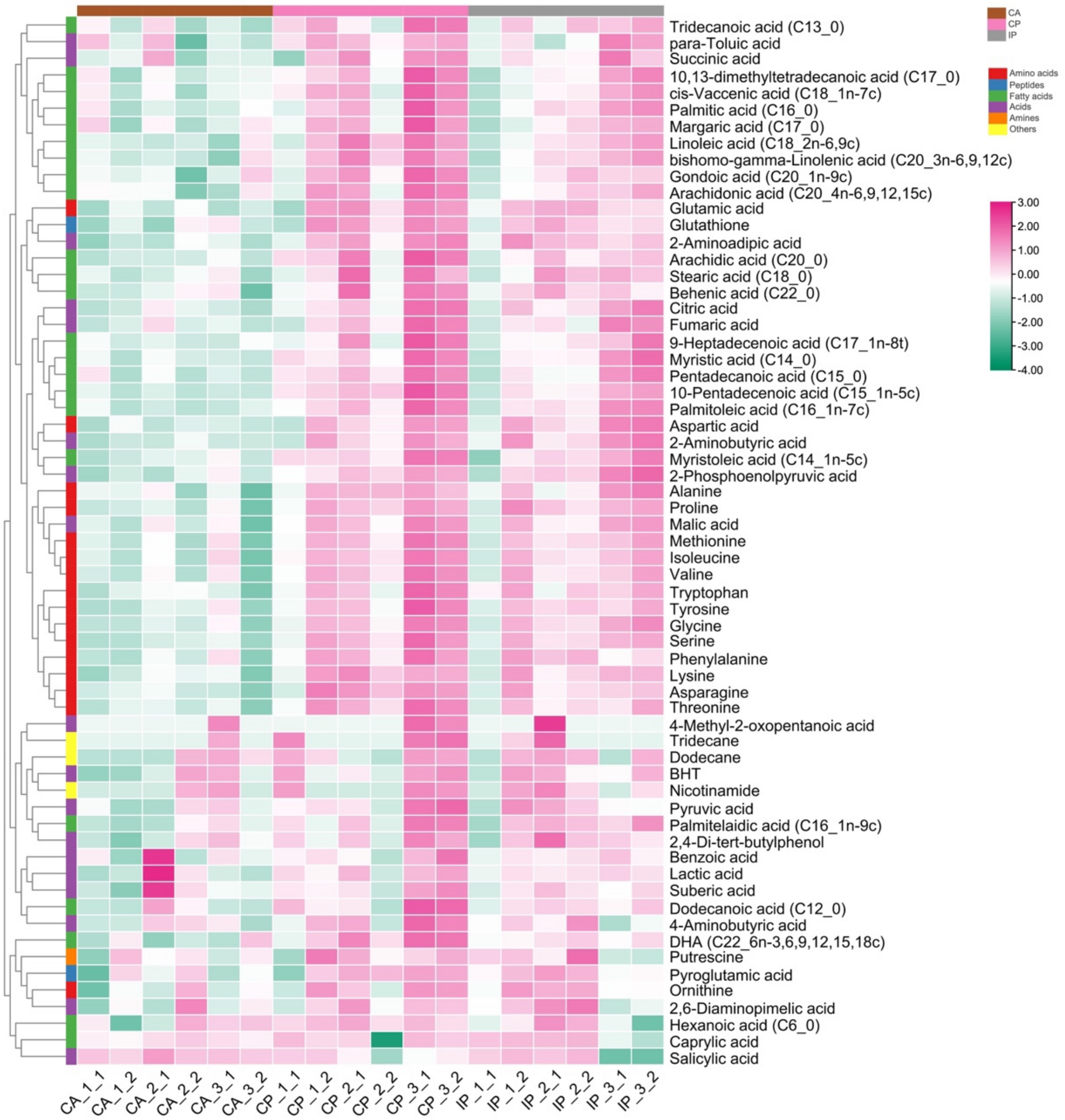


**Figure S35.** Relative abundance of all intracellular metabolites identified in biomass samples from activated sludge exposed to different aeration strategies after 48 hours. The data is normalized by internal standard, Log10 transformation, and Pareto scaling. Row clustering according to ‘Ward’ (technical duplication results for biological triplicates; n=6)

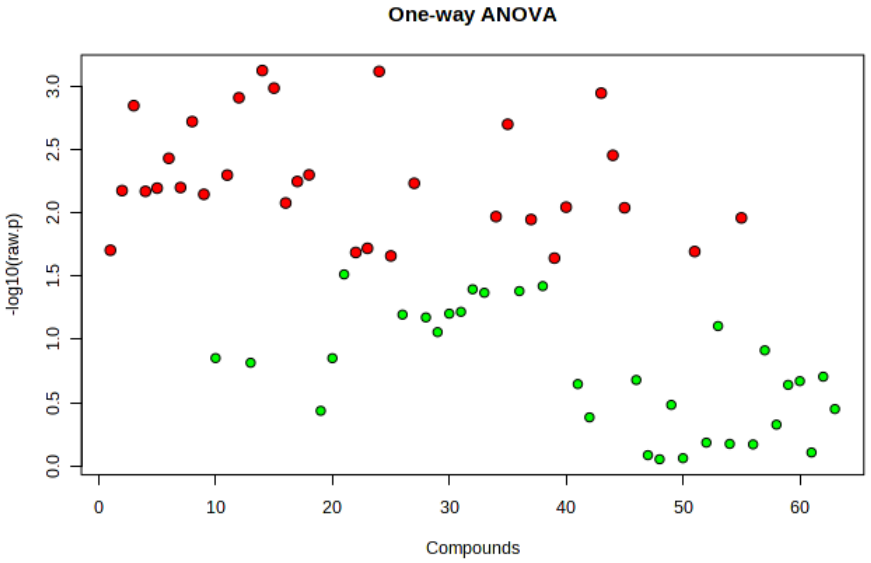


**Figure S36.** Differential abundance analysis of metabolites by ANOVA analysis (31 significant differences out of 63 metabolites, *P*<0.05)

**Table S2.** Metabolites with significant differences in abundance among the three aeration conditions (ANOVA, FDR< 0.05)

| Metabolites | f.value | p.value | -log10(p) | FDR | Tukey's LSD |
|---|---|---|---|---|---|
| Serine | 12.092 | 0.00074538 | 3.1276 | 0.014844 | CP - CA; IP - CA |
| 10-Pentadecenoic acid (C15_1n-5c) | 12.05 | 0.00075741 | 3.1207 | 0.014844 | CP - CA; IP - CA |
| Threonine | 11.272 | 0.001027 | 2.9884 | 0.014844 | CP - CA; IP - CA |
| 2-Aminoadipic acid | 11.045 | 0.0011251 | 2.9488 | 0.014844 | CP - CA; IP - CA |
| Proline | 10.84 | 0.0012234 | 2.9124 | 0.014844 | CP - CA; IP - CA |
| Asparagine | 10.489 | 0.0014137 | 2.8496 | 0.014844 | CP - CA; IP - CA |
| Lysine | 9.8064 | 0.0018897 | 2.7236 | 0.015623 | CP - CA; IP - CA |
| Arachidic acid (C20_0) | 9.6946 | 0.0019839 | 2.7025 | 0.015623 | CP - CA; IP - CA |
| 2-Aminobutyric acid | 8.4454 | 0.0034929 | 2.4568 | 0.02324 | CP - CA; IP - CA |
| Glycine | 8.3298 | 0.0036889 | 2.4331 | 0.02324 | CP - CA; IP - CA |

| | | | | | |
|---|---|---|---|---|---|
| Valine | 7.7055 | 0.0049884 | 2.302 | 0.023577 | CP - CA; IP - CA |
| Phenylalanine | 7.6963 | 0.005011 | 2.3001 | 0.023577 | CP - CA; IP - CA |
| Tyrosine | 7.463 | 0.0056276 | 2.2497 | 0.023577 | CP - CA; IP - CA |
| Palmitoleic acid (C16_1n-7c) | 7.3972 | 0.0058165 | 2.2353 | 0.023577 | CP - CA; IP - CA |
| Isoleucine | 7.2448 | 0.0062829 | 2.2018 | 0.023577 | CP - CA; IP - CA |
| Glutamic acid | 7.223 | 0.0063532 | 2.197 | 0.023577 | CP - CA; IP - CA |
| Alanine | 7.1348 | 0.0066459 | 2.1774 | 0.023577 | CP - CA; IP - CA |
| Aspartic acid | 7.1085 | 0.0067364 | 2.1716 | 0.023577 | CP - CA; IP - CA |
| Methionine | 7.0035 | 0.0071107 | 2.1481 | 0.023578 | CP - CA; IP - CA |
| Tryptophan | 6.7015 | 0.0083262 | 2.0796 | 0.026018 | CP - CA; IP - CA |
| DHA (C22_6n-3,6,9,12,15,18c) | 6.5581 | 0.0089846 | 2.0465 | 0.026018 | CP - CA |
| 2-Phosphoenolpyruvic acid | 6.5372 | 0.0090856 | 2.0416 | 0.026018 | CP - CA; IP - CA |
| Linoleic acid (C18_2n-6,9c) | 6.239 | 0.010673 | 1.9717 | 0.028353 | CP - CA |
| Malic acid | 6.1959 | 0.010928 | 1.9615 | 0.028353 | CP - CA; IP - CA |
| bishomo-gamma-Linolenic acid (C20_3n-6,9,12c) | 6.1428 | 0.011251 | 1.9488 | 0.028353 | CP - CA; IP - CA |
| Myristoleic acid (C14_1n-5c) | 5.2202 | 0.019021 | 1.7208 | 0.044568 | CP - CA |
| Glutathione | 5.1609 | 0.019701 | 1.7055 | 0.044568 | CP - CA; IP - CA |
| Citric acid | 5.1211 | 0.020171 | 1.6953 | 0.044568 | CP - CA; IP - CA |
| Myristic acid (C14_0) | 5.0926 | 0.020516 | 1.6879 | 0.044568 | CP - CA; IP - CA |
| Pentadecanoic acid (C15_0) | 4.987 | 0.021854 | 1.6605 | 0.045893 | CP - CA |
| Behenic acid (C22_0) | 4.9211 | 0.022738 | 1.6432 | 0.04621 | CP - CA; IP - CA |

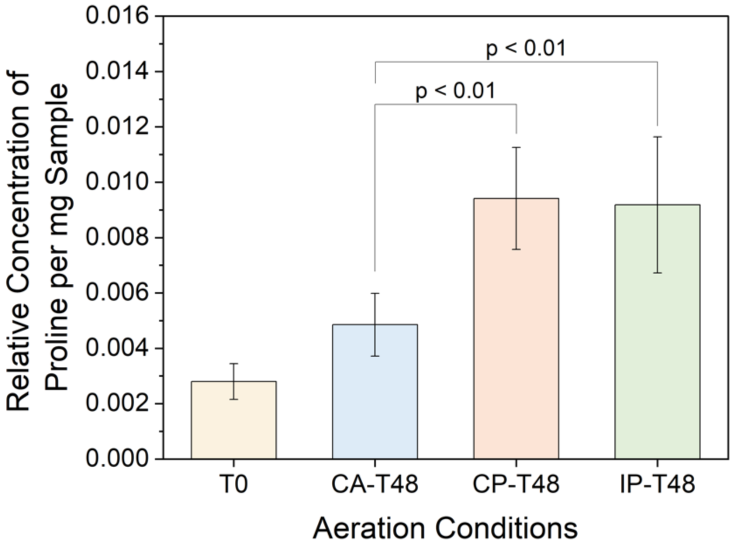


**Figure S37.** Differences in the abundance of proline in activated sludge biomass samples. *P* values obtained from ANOVA Tukey test (*P* values greater than 0.05 between the three aeration conditions are not shown). The error bars represent standard deviations (technical duplication results for biological triplicates; n=6)

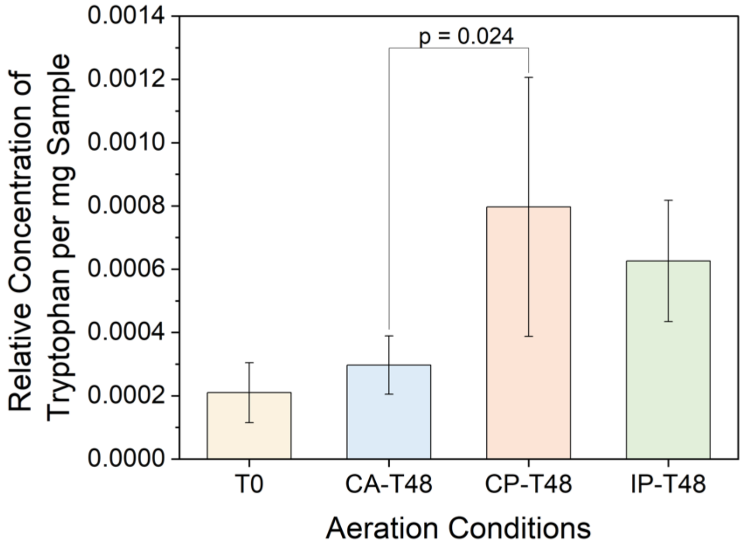


**Figure S38.** Differences in the abundance of tryptophan in activated sludge biomass samples. *P* values obtained from ANOVA Tukey test (*P* values greater than 0.05 between the three aeration conditions are not shown). The error bars represent standard deviations (technical duplication results for biological triplicates; n=6)

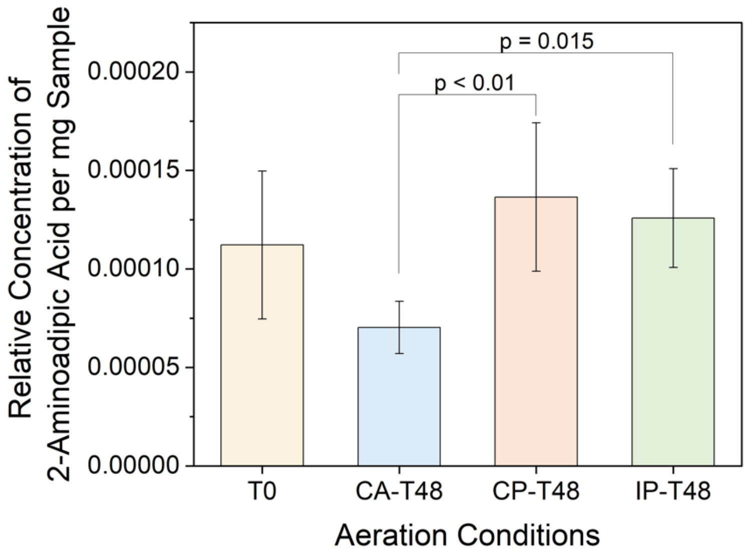


**Figure S39.** Differences in the abundance of 2-aminoadipic in activated sludge biomass samples. *P* values obtained from ANOVA Tukey test (*P* values greater than 0.05 between the three aeration conditions are not shown). The error bars represent standard deviations (technical duplication results for biological triplicates; n=6)

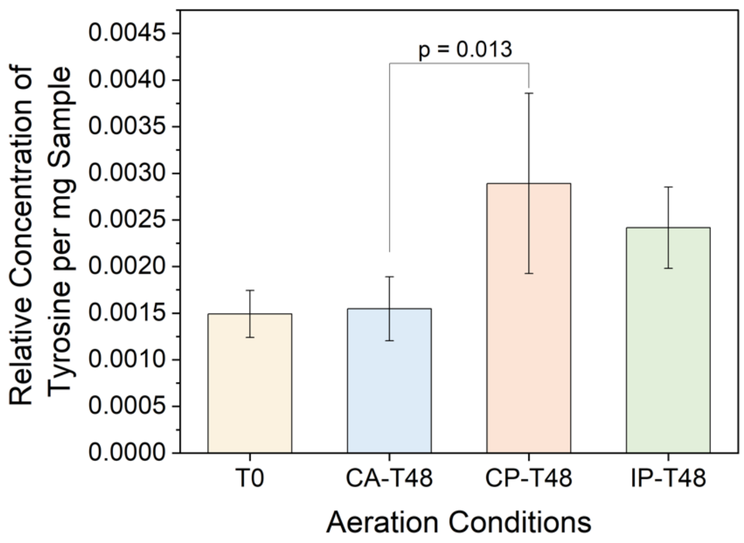


**Figure S40.** Differences in the abundance of tyrosine in activated sludge biomass samples. *P* values obtained from ANOVA Tukey test (*P* values greater than 0.05 between the three aeration conditions are not shown). The error bars represent standard deviations (technical duplication results for biological triplicates; n=6)

##### S2.6. BIO-Sankey network analysis

Note: In BIO-Sankey network diagram, dark red (or green) bars indicate the microbes or metabolites that are significantly higher (or lower) under the CP or IP ($FC>1$ or $FC<1$, $P<0.05$); Light red (or green) bars indicate the microbes or metabolites that are higher (or lower) under the CP or IP ($FC>1$ or $FC<1$, $P\geq0.05$); Black bars indicate the microbes or metabolites in the reference database; Purple bars indicate the metabolic enzymes; Dark red (or green) bands indicate significant positive (or negative) correlations (Spearman correlation test; $R>$ or $R<0$, $P<0.05$); Light red (or green) bands indicate positive (or negative) correlations without statistical significance (Spearman correlation test; $R>$or $R<0$, $P\geq0.05$).

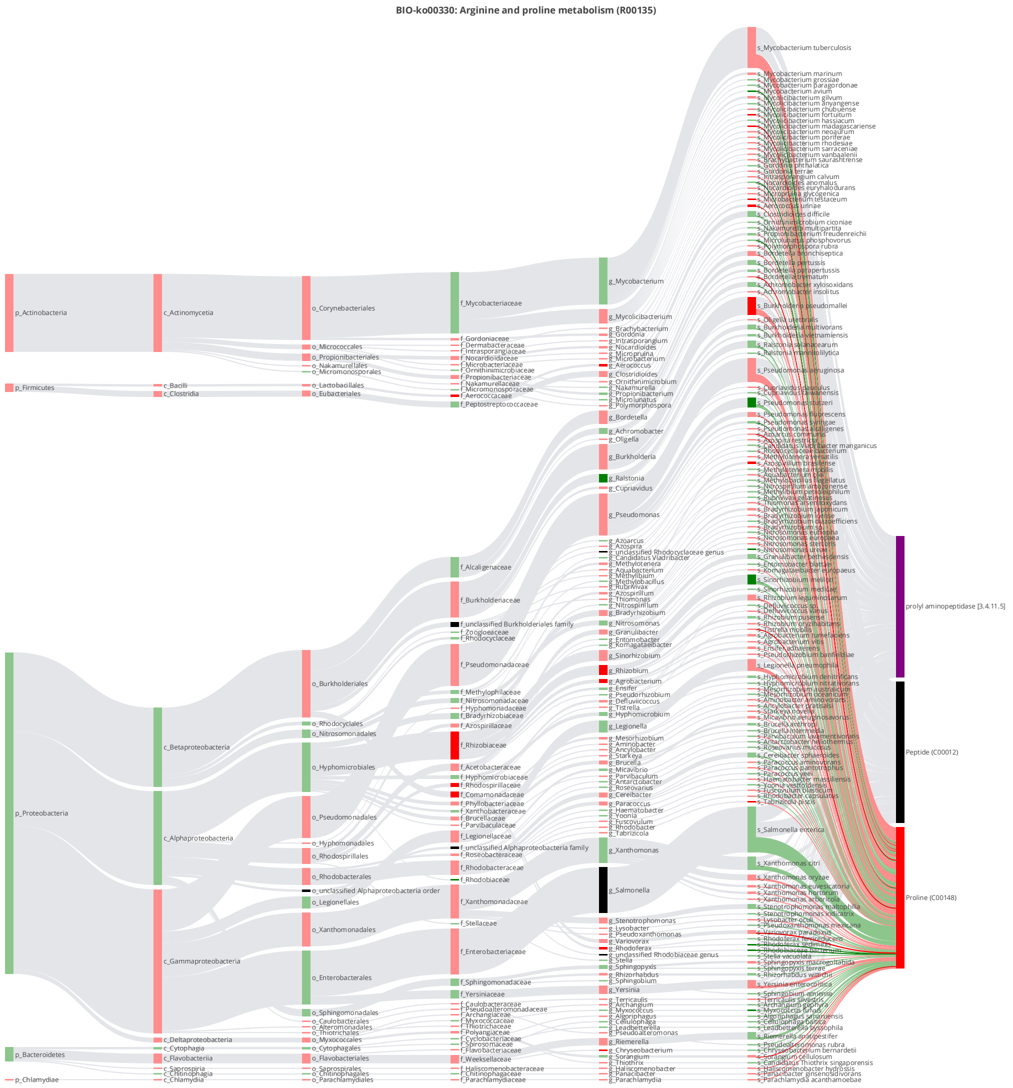


**Figure S41.** BIO-Sankey network diagram (Hypergeometric test, log0.05 p-value > 1) for R00135 of arginine and proline metabolism (ko00330) when comparing CP with CA condition (the reaction involving proline (C00148) and cytosol aminopeptidase [EC:3.4.11.1 3.4.11.5] encoded by K11142)

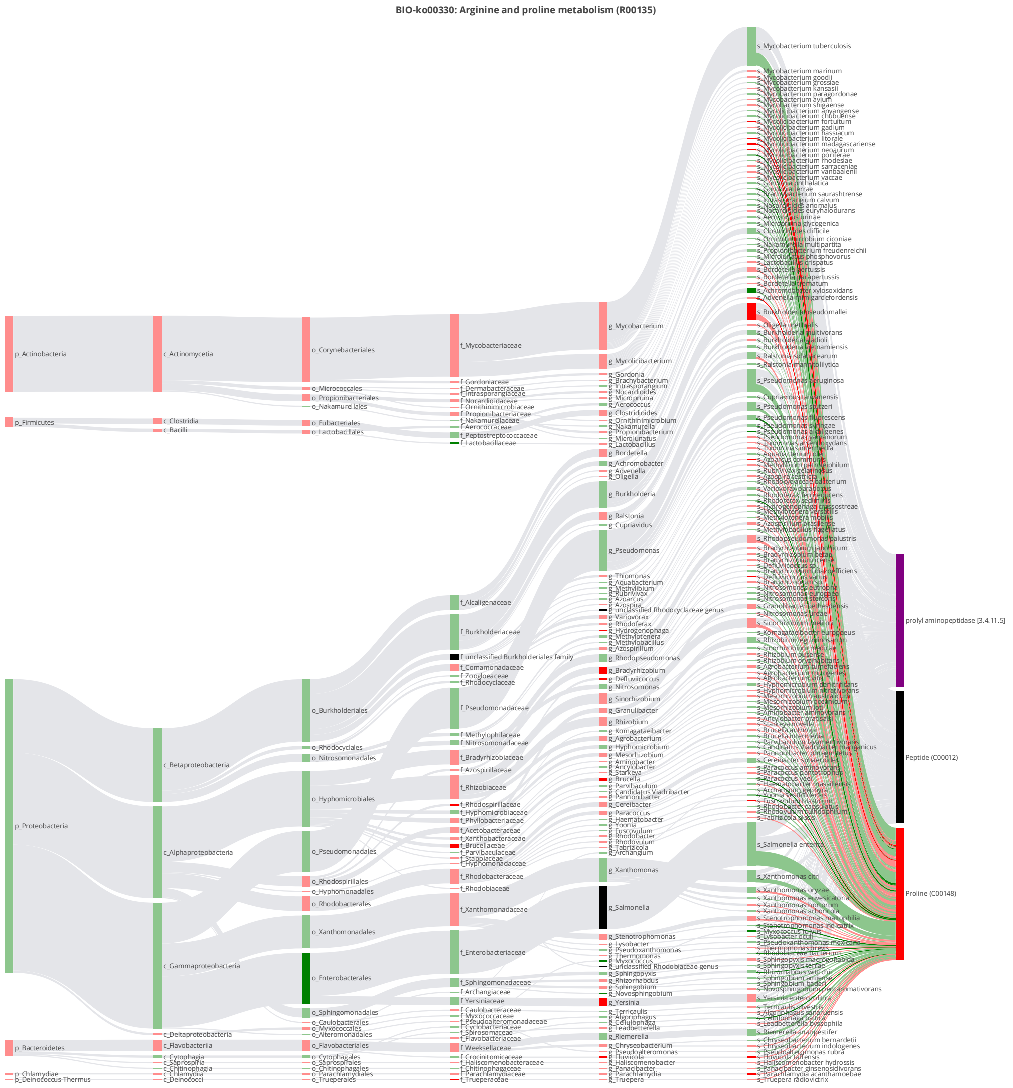


**Figure S42.** BIO-Sankey network diagram (Hypergeometric test, log0.05 p-value > 1) for R00135 of arginine and proline metabolism (ko00330) when comparing IP with CA condition (the reaction involving proline (C00148) and cytosol aminopeptidase [EC:3.4.11.1 3.4.11.5] encoded by K11142)

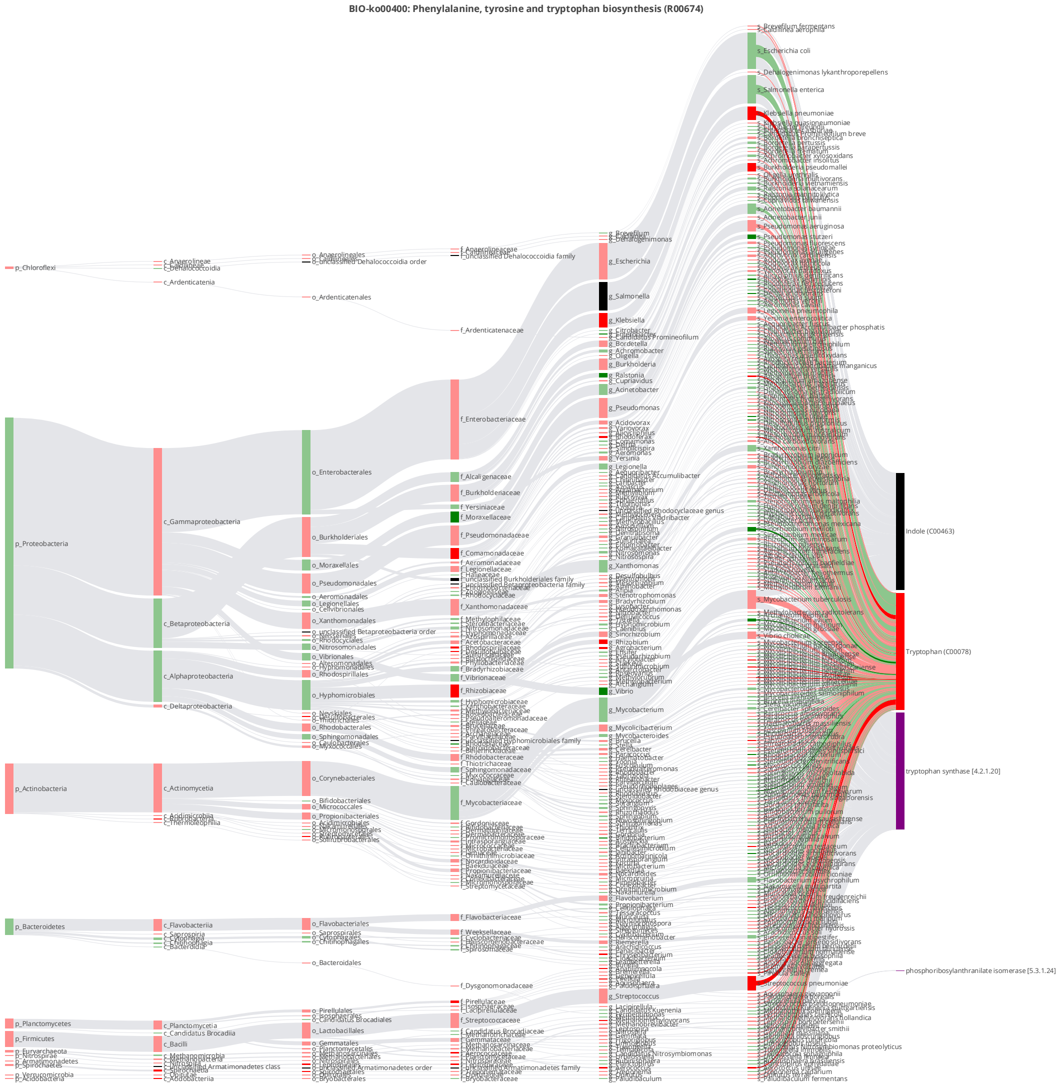


**Figure S43.** BIO-Sankey network diagram (Hypergeometric test, log0.05 p-value > 1) for R00674 of phenylalanine, tyrosine, and tryptophan biosynthesis (ko00400) when comparing CP with CA condition (the reaction involving tryptophan (C00078) and indole-3-glycerol phosphate synthase / phosphoribosylanthranilate isomerase [EC:4.1.1.48 5.3.1.24] encoded by K13498)

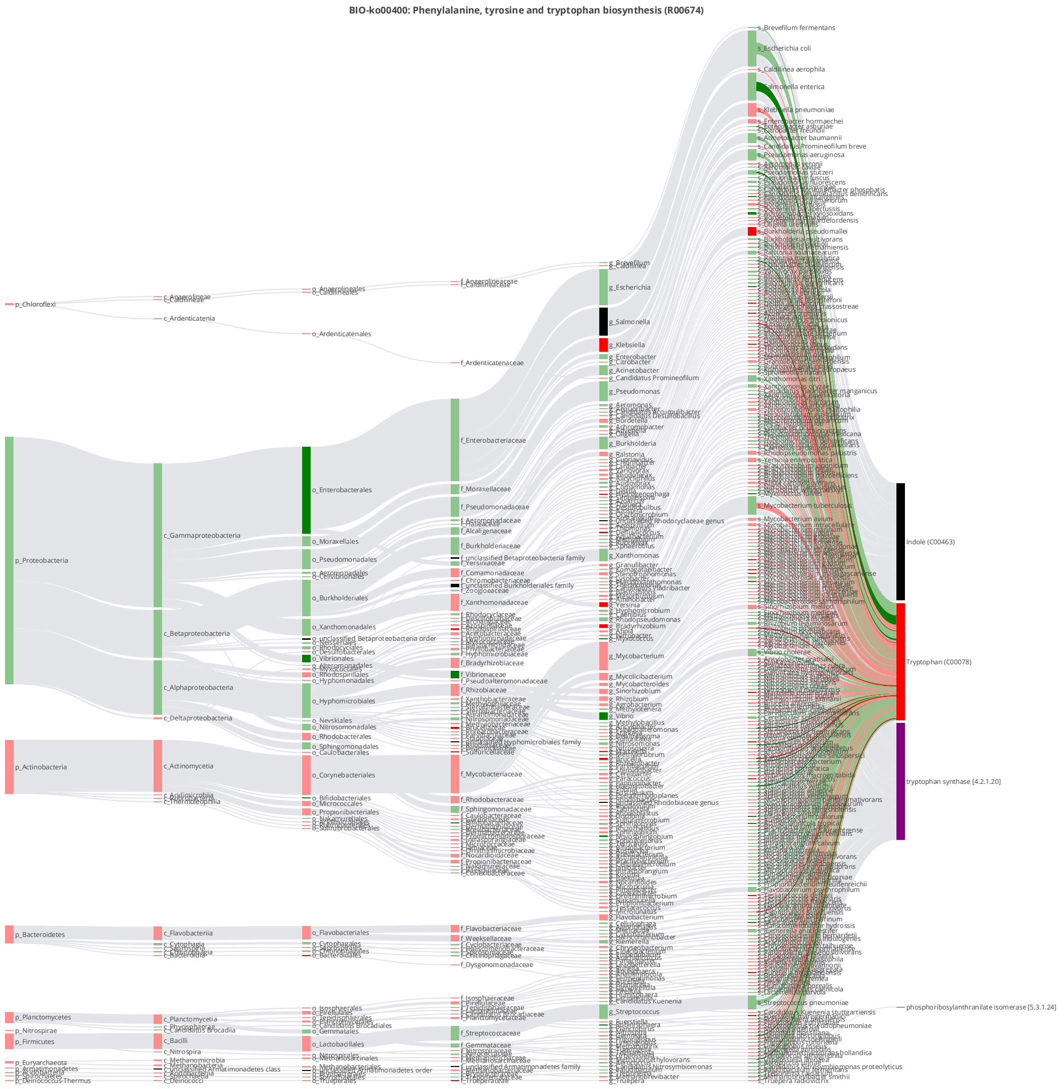


**Figure S44.** BIO-Sankey network diagram (Hypergeometric test, log0.05 p-value > 1) for R00674 of phenylalanine, tyrosine, and tryptophan biosynthesis (ko00400) when comparing IP with CA condition (the reaction involving tryptophan (C00078) and indole-3-glycerol phosphate synthase / phosphoribosylanthranilate isomerase [EC:4.1.1.48 5.3.1.24] encoded by K13498)

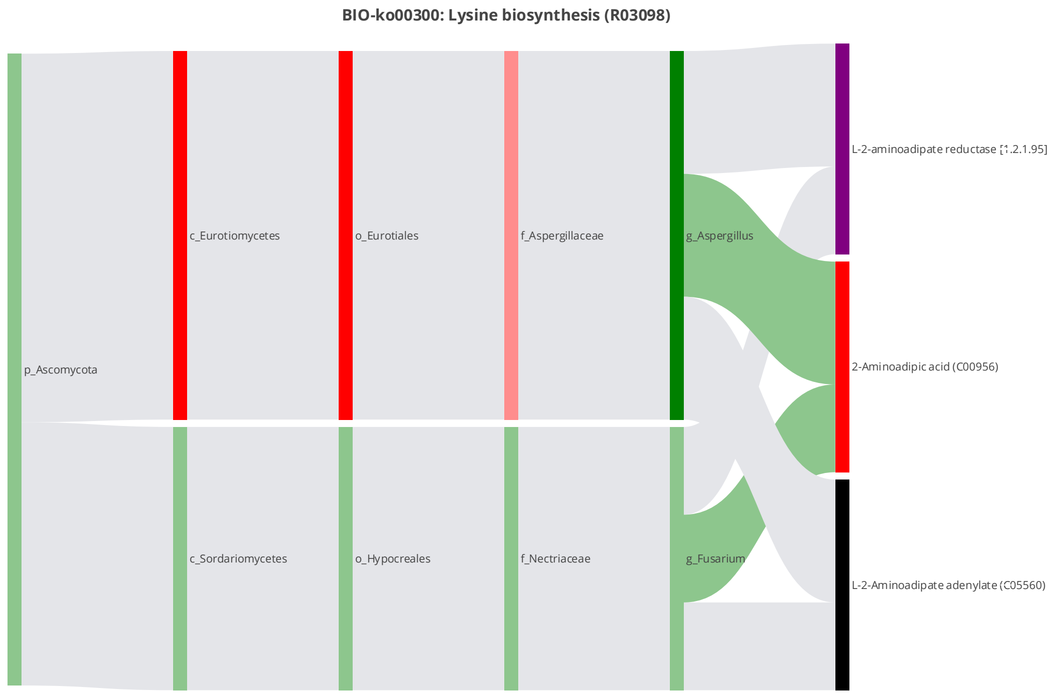


**Figure S45.** BIO-Sankey network diagram (Hypergeometric test, log0.05 p-value > 1) for R03098 of lysine biosynthesis (ko00300) when comparing CP with CA condition (the reaction involving 2-aminoadipic acid (C00956) and L-2-aminoadipate reductase [EC:1.2.1.95] encoded by K00143)

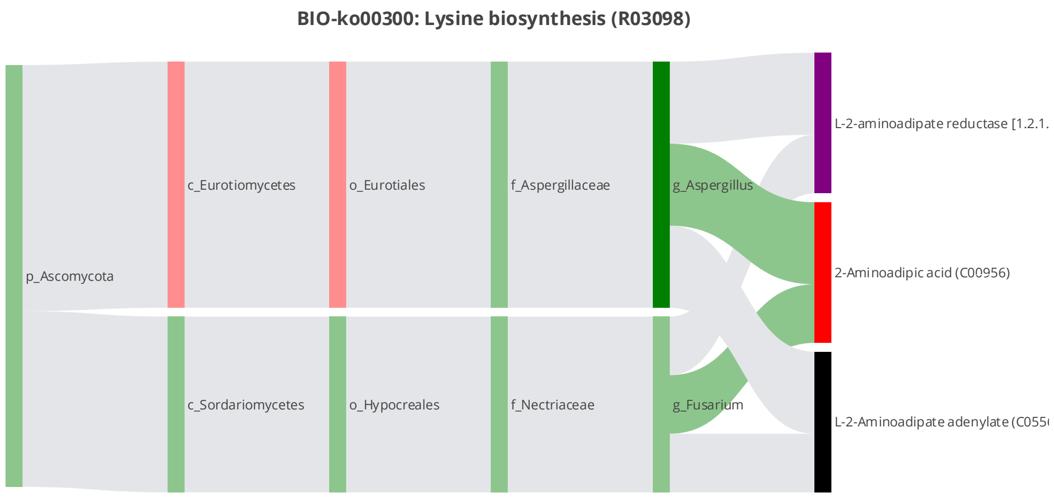


**Figure S46.** BIO-Sankey network diagram (Hypergeometric test, log0.05 p-value > 1) for R03098 of lysine biosynthesis (ko00300) when comparing IP with CA condition (the reaction involving 2-aminoadipic acid (C00956) and L-2-aminoadipate reductase [EC:1.2.1.95] encoded by K00143)

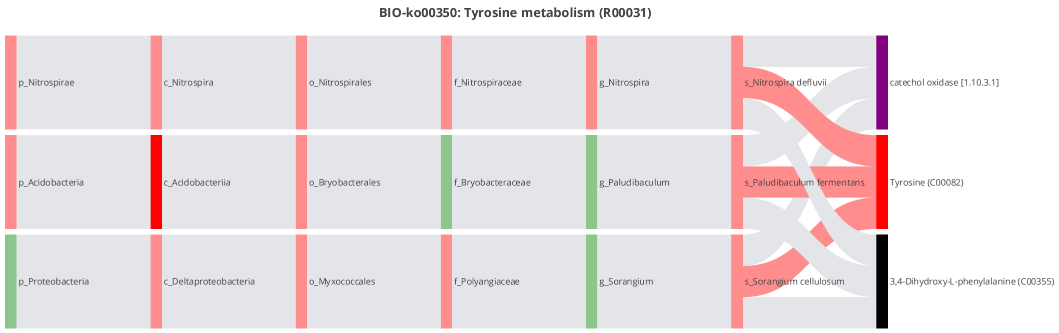


**Figure S47.** BIO-Sankey network diagram (Hypergeometric test, log0.05 p-value > 1) for R00031 of tyrosine metabolism (ko00350) when comparing CP with CA condition (the reaction involving tyrosine (C00082) and polyphenol oxidase [EC:1.10.3.1] encoded by K00422)

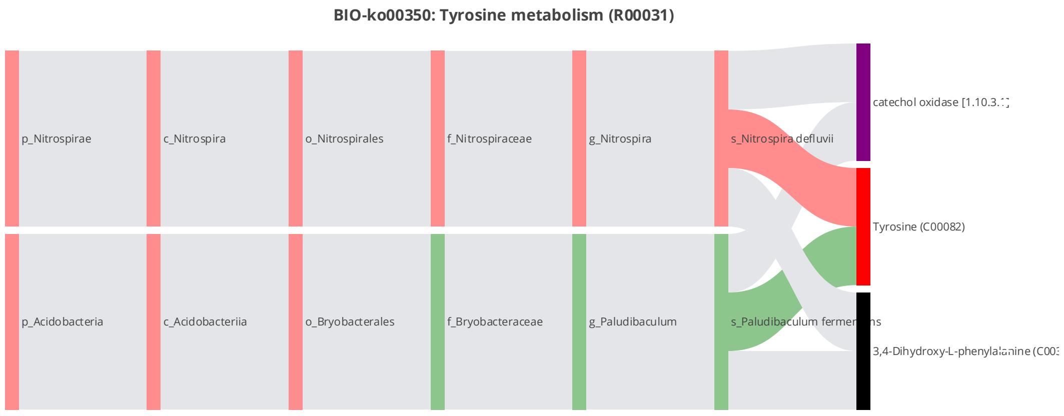


**Figure S48.** BIO-Sankey network diagram (Hypergeometric test, log0.05 p-value > 1) for R00031 of tyrosine metabolism (ko00350) when comparing IP with CA condition (the reaction involving tyrosine (C00082) and polyphenol oxidase [EC:1.10.3.1] encoded by K00422)

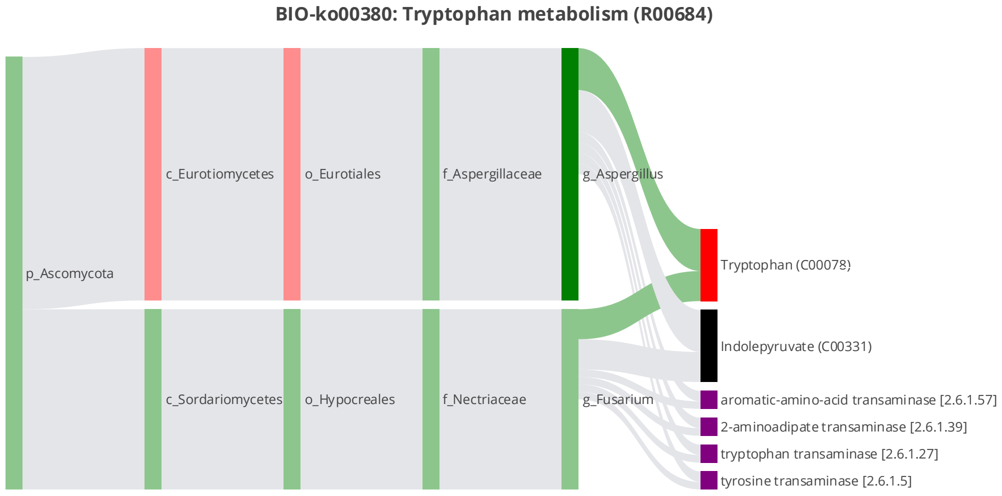


**Figure S49.** BIO-Sankey network diagram (Hypergeometric test, log0.05 p-value > 1) for R00684 of tryptophan metabolism (ko00380) when comparing IP with CA condition (the reaction involving tryptophan (C00078) and tryptophan aminotransferase [EC:2.6.1.27] encoded by K14265)

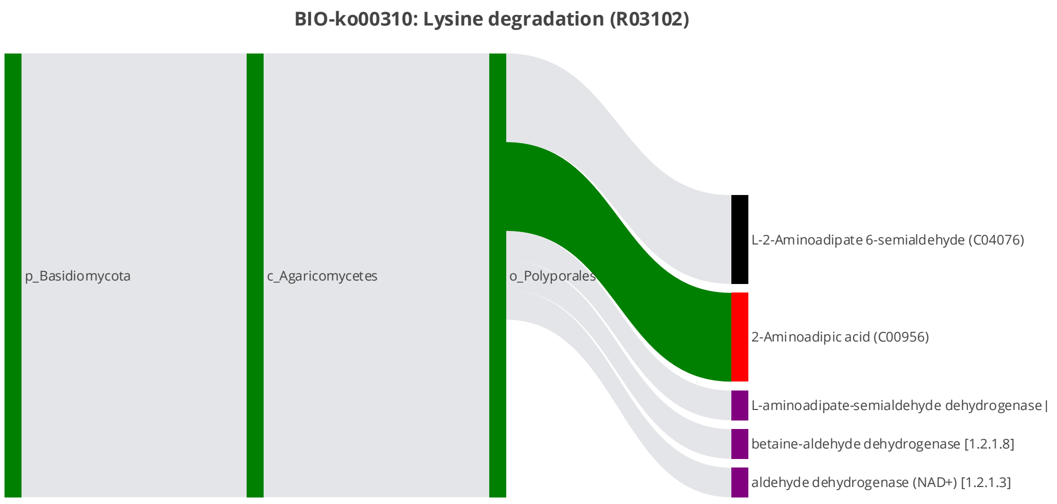


**Figure S50.** BIO-Sankey network diagram (Hypergeometric test, log0.05 p-value > 1) for R03102 of tryptophan metabolism (ko00310) when comparing IP with CA condition (the reaction involving 2-aminoadipic acid (C00956) and aldehyde dehydrogenase family 9 member A1 [EC:1.2.1.47 1.2.1.3] encoded by K00149)

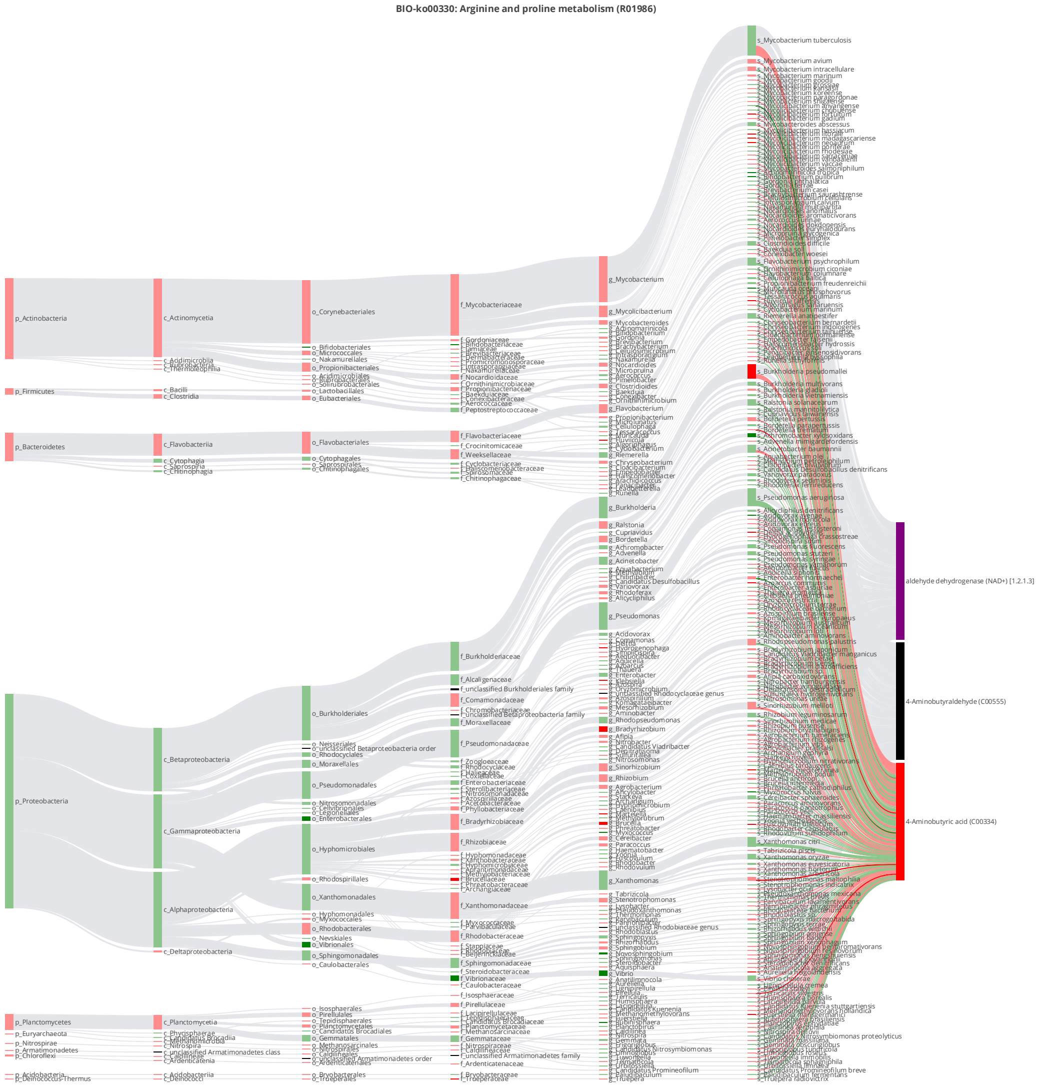


**Figure S51.** BIO-Sankey network diagram (Hypergeometric test, log0.05 p-value > 1) for R01986 of arginine and proline metabolism(ko00330) when comparing IP with CA condition (the reaction involving 4-aminobutyric acid (C00334) and aldehyde dehydrogenase family 9 member A1 [EC:1.2.1.47 1.2.1.3] encoded by K00149)

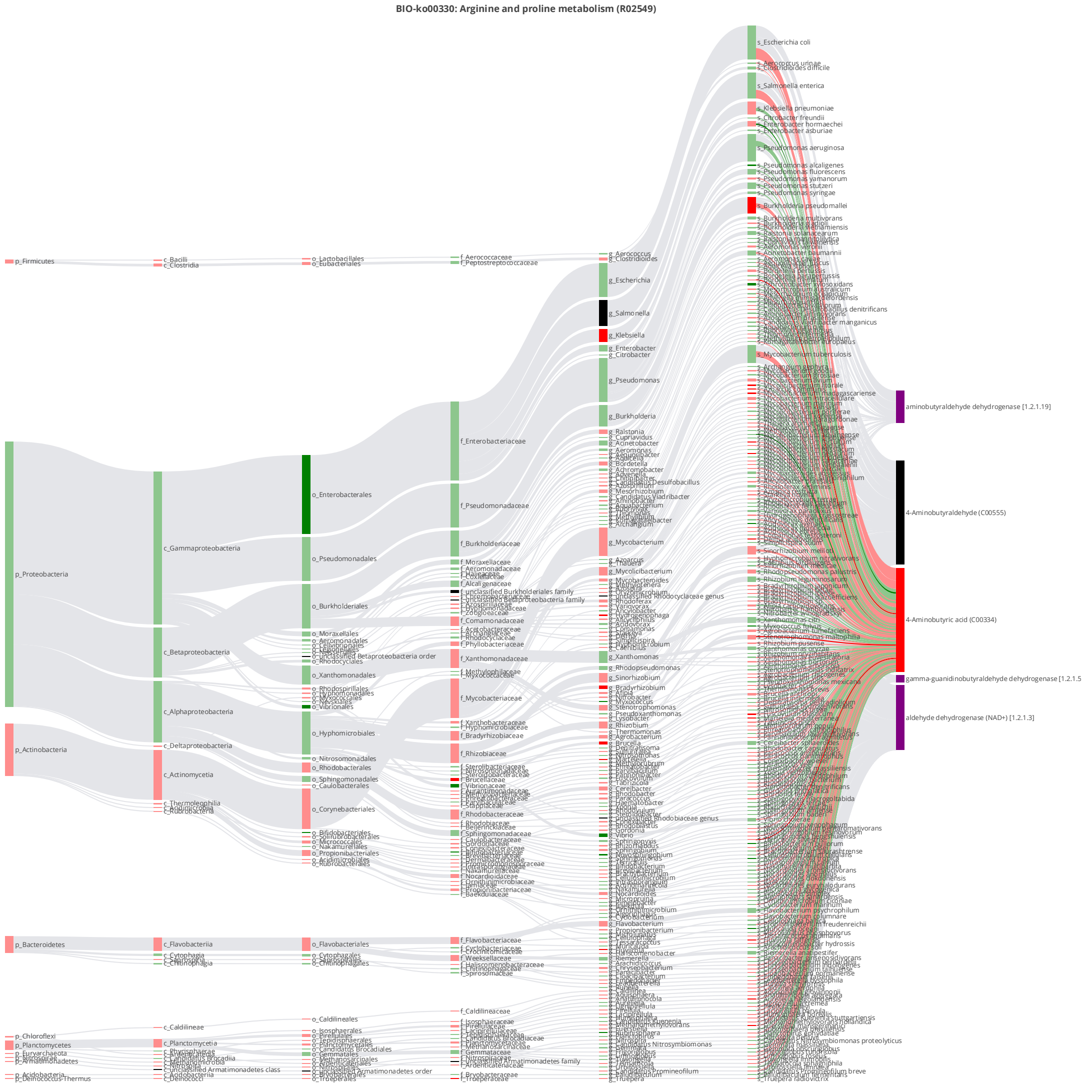


**Figure S52.** BIO-Sankey network diagram (Hypergeometric test, log0.05 p-value > 1) for R02549 of arginine and proline metabolism(ko00330) when comparing IP with CA condition (the reaction involving 4-aminobutyric acid (C00334) and aldehyde dehydrogenase family 9 member A1 [EC:1.2.1.47 1.2.1.3] encoded by K00149)

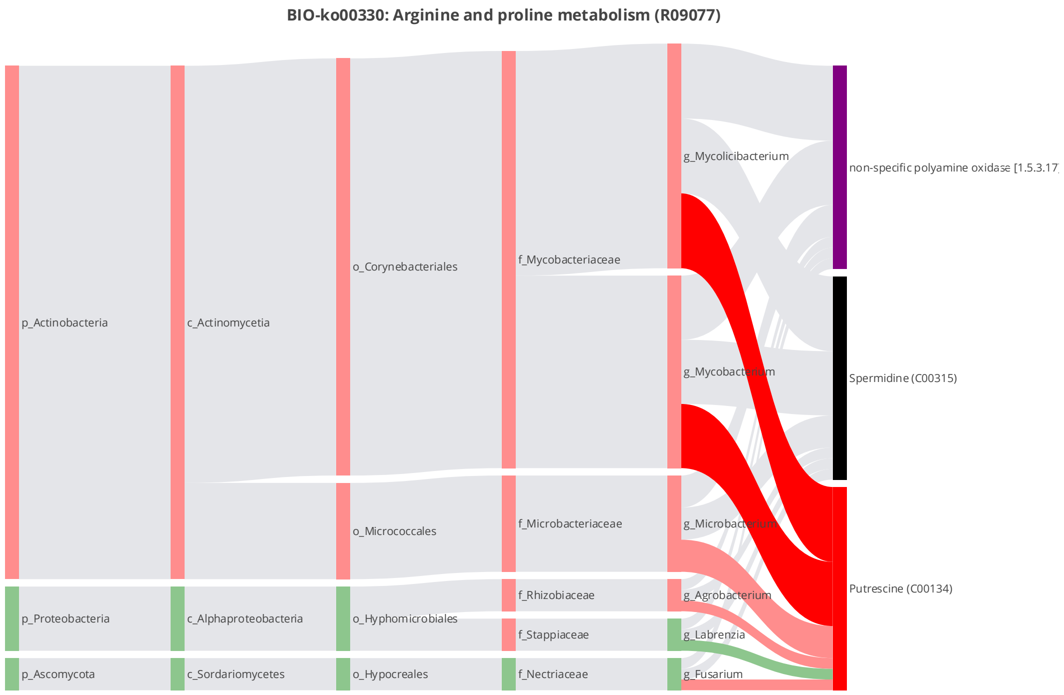


**Figure S53.** BIO-Sankey network diagram (Hypergeometric test, log0.05 p-value > 1) for R09077 of arginine and proline metabolism(ko00330) when comparing IP with CA condition (the reaction involving putrescine (C00134) and polyamine oxidase [EC:1.5.3.17] encoded by K13367)

**S2.7. STA-Sankey network analysis**

Note: The dark red (or green) bars indicate the microbes or metabolites that are significantly higher (or lower) in the high-altitude population (FC>1 or FC<1, $P$<0.05); light red (or green) bars indicate the microbes or metabolites that are higher (or lower) in the high-altitude population (FC>1 or FC<1, $P$≥0.05); dark grey bars indicate the microbes or metabolites with no change (FC = 1); Dark red (or green) bands indicate significant positive (or negative) correlations (Spearman correlation test; R>or R<0, $P$<0.05); Light red (or green) bands indicate positive (or negative) correlations without statistical significance (Spearman correlation test; R>or R<0, $P$≥0.05); Light gray bands indicate reference relationships searched through database. The asterisk (*) indicates an abiologically significant correlation with the metabolite.

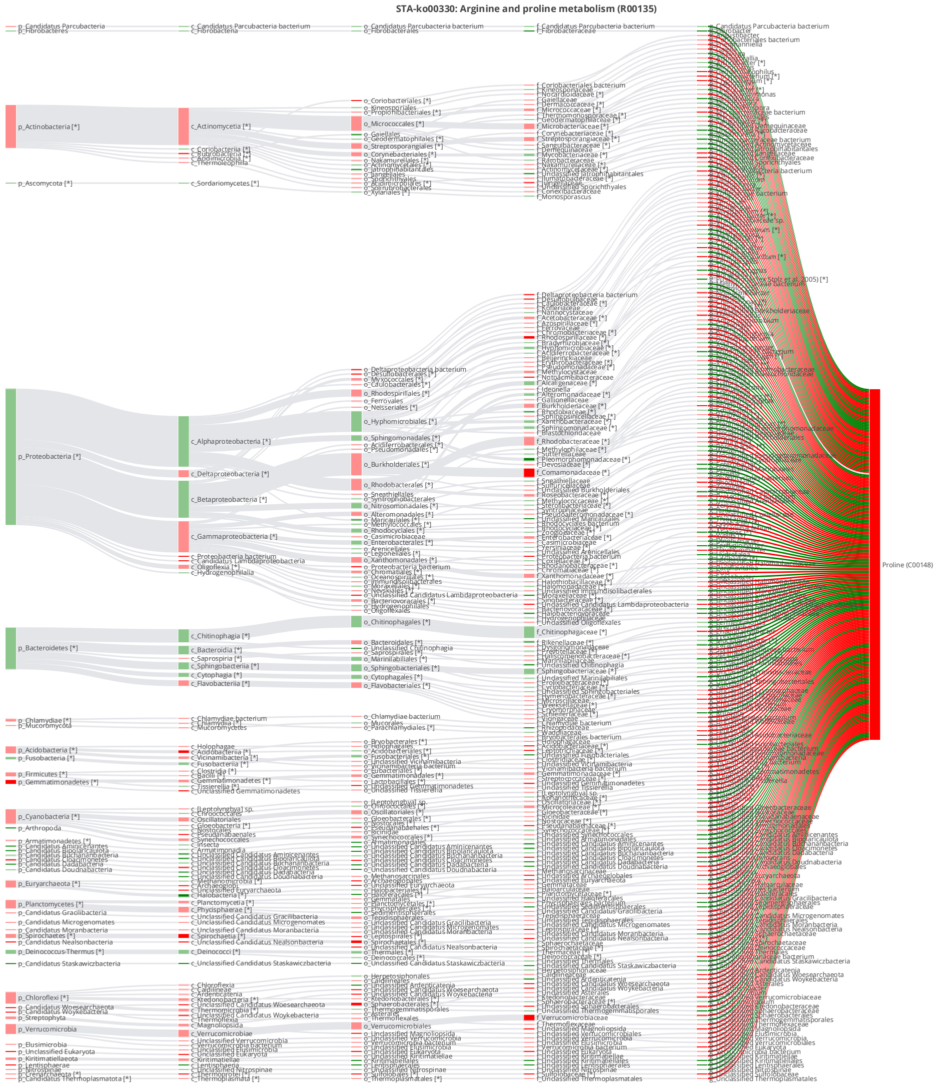


**Figure S54.** STA-Sankey network diagram (Hypergeometric test, log0.05 p-value > 1) for R00135 of arginine and proline metabolism(ko00330) when comparing CP with CA condition (the reaction involving proline (C00148) and cytosol aminopeptidase [EC:3.4.11.1 3.4.11.5] encoded by K11142)

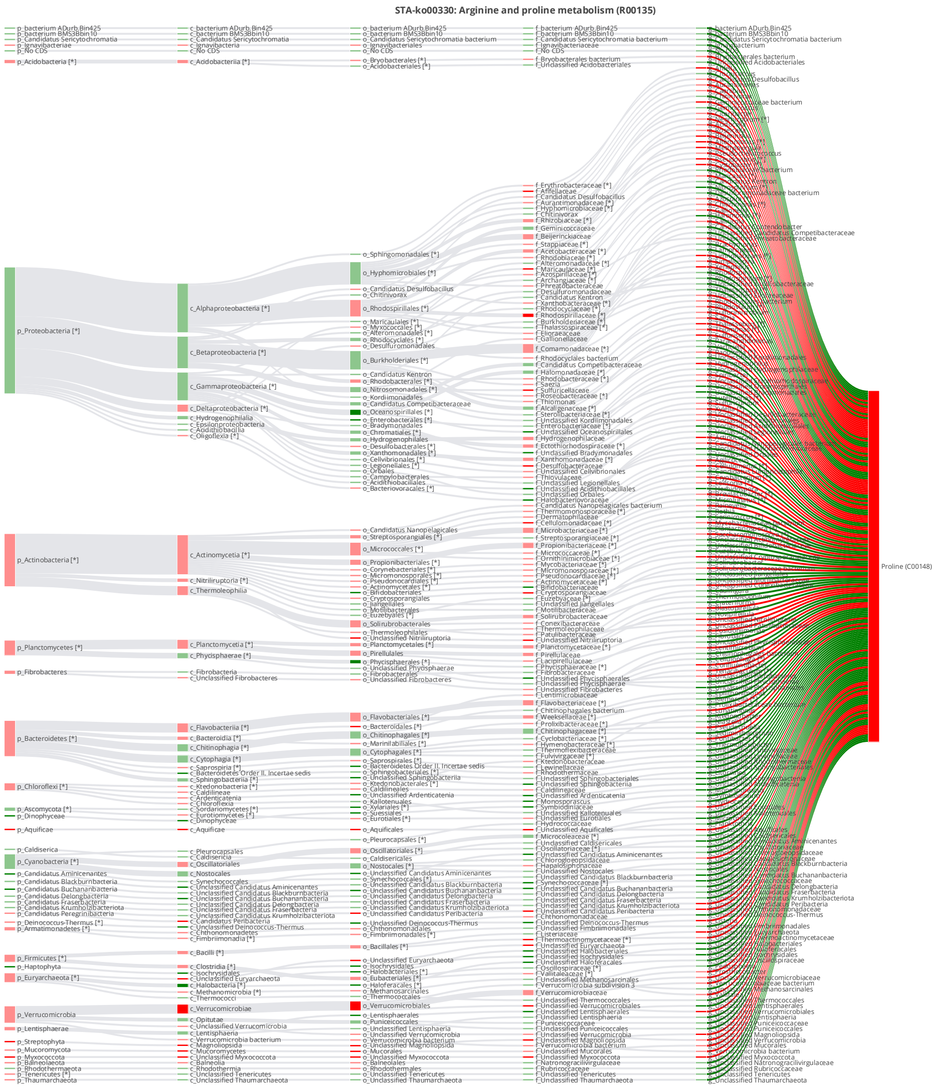


**Figure S55.** STA-Sankey network diagram (Hypergeometric test, log0.05 p-value > 1) for R00135 of arginine and proline metabolism(ko00330) when comparing IP with CA condition (the reaction involving proline (C00148) and cytosol aminopeptidase [EC:3.4.11.1 3.4.11.5] encoded by K11142)

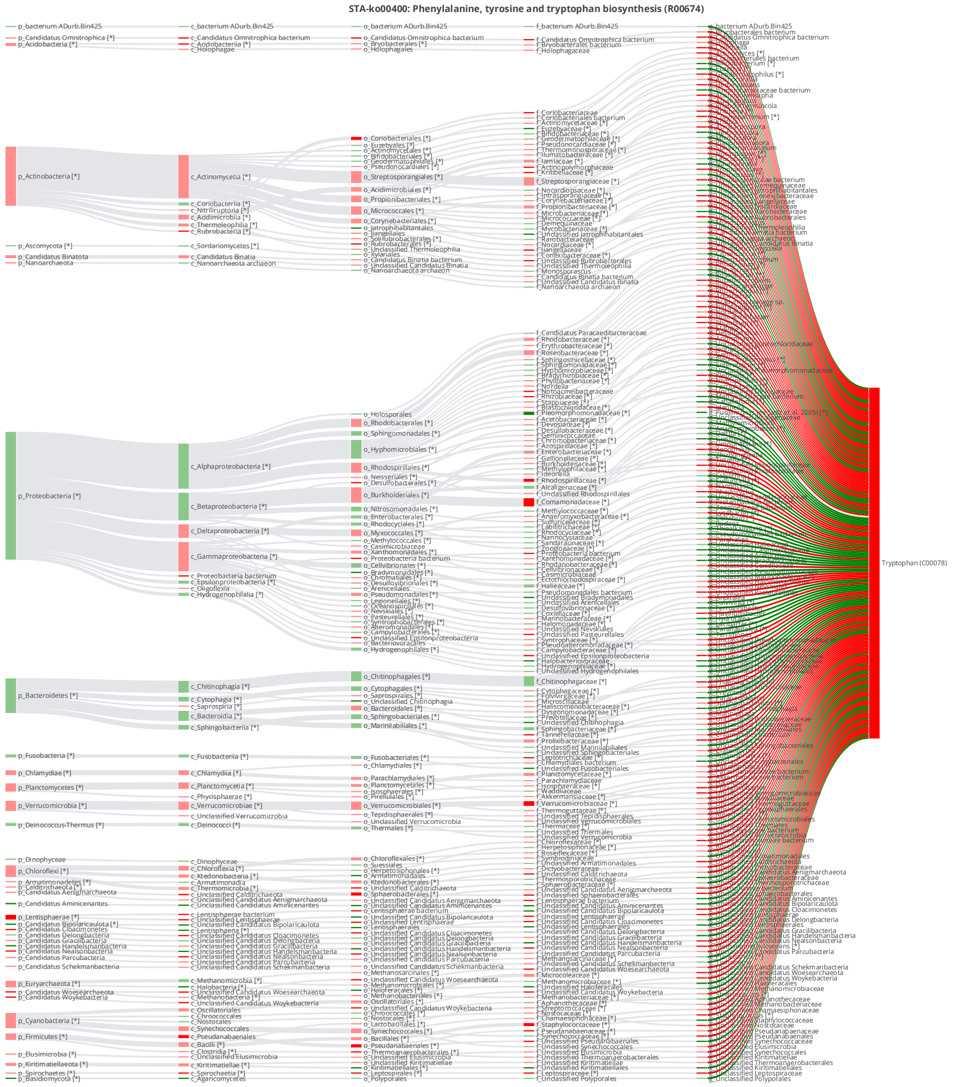


**Figure S56.** STA-Sankey network diagram (Hypergeometric test, log0.05 p-value > 1) for R00674 of phenylalanine, tyrosine, and tryptophan biosynthesis (ko00400) when comparing CP with CA condition (the reaction involving tryptophan (C00078) and indole-3-glycerol phosphate synthase / phosphoribosylanthranilate isomerase [EC:4.1.1.48 5.3.1.24] encoded by K13498)

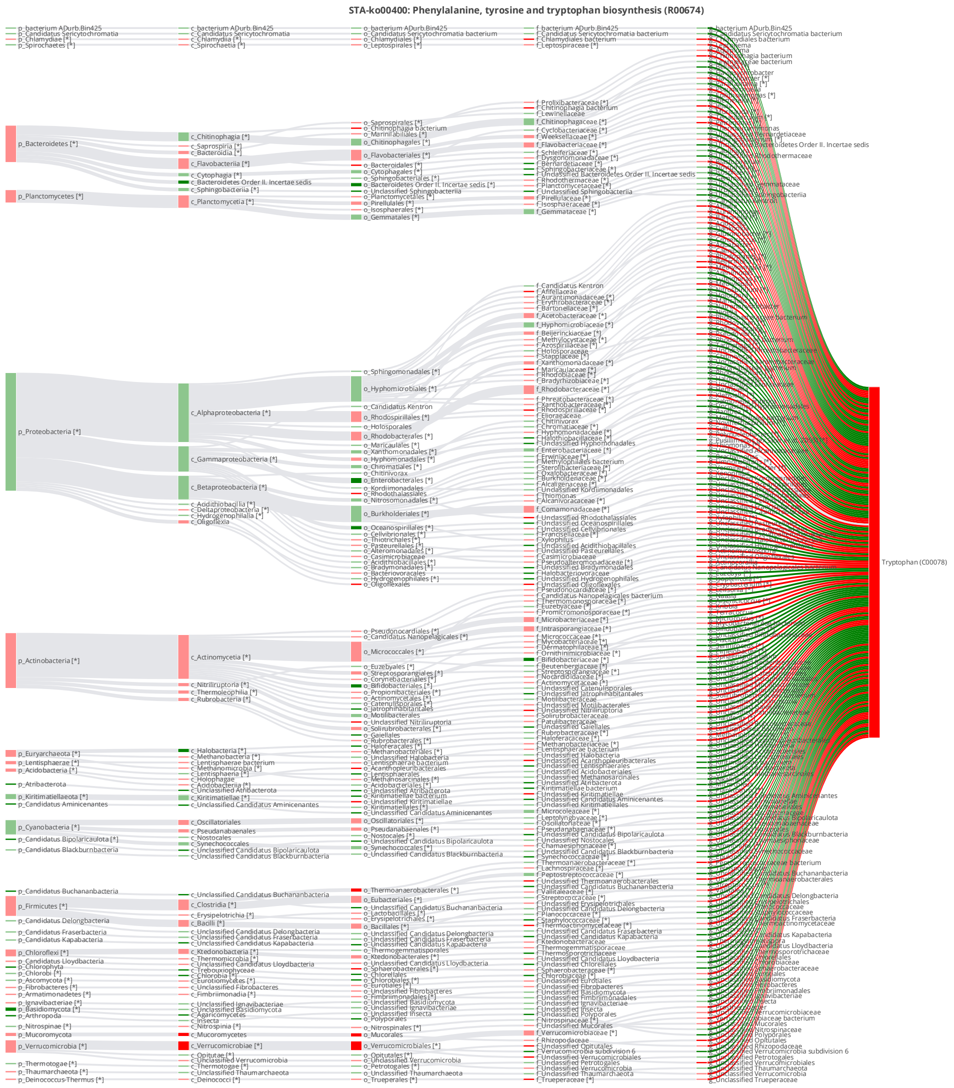


**Figure S57.** STA-Sankey network diagram (Hypergeometric test, log0.05 p-value > 1) for R00674 of phenylalanine, tyrosine, and tryptophan biosynthesis (ko00400) when comparing IP with CA condition (the reaction involving tryptophan (C00078) and indole-3-glycerol phosphate synthase / phosphoribosylanthranilate isomerase [EC:4.1.1.48 5.3.1.24] encoded by K13498)

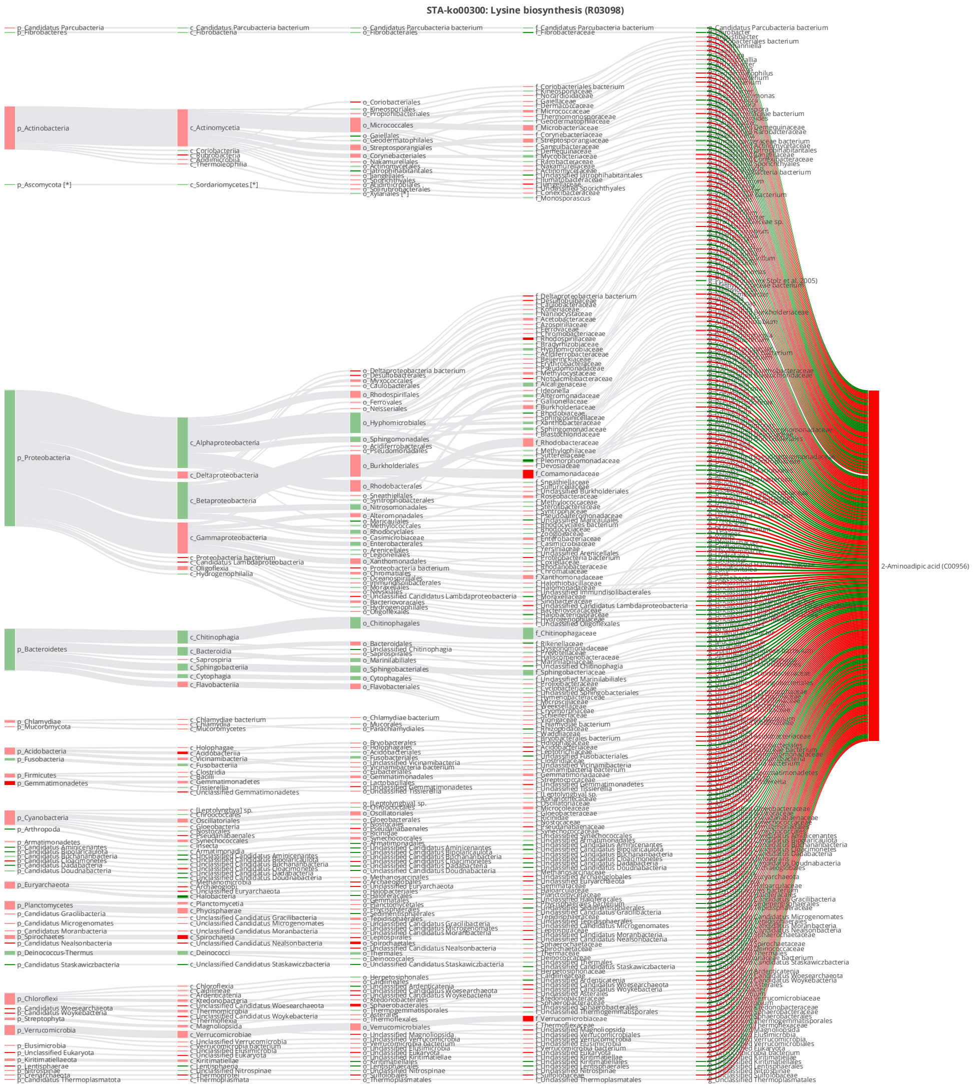


**Figure S58.** STA-Sankey network diagram (Hypergeometric test, log0.05 p-value > 1) for R03098 of lysine biosynthesis (ko00300) when comparing CP with CA condition (the reaction involving 2-aminoadipic acid (C00956) and L-2-aminoadipate reductase [EC:1.2.1.95] encoded by K00143)

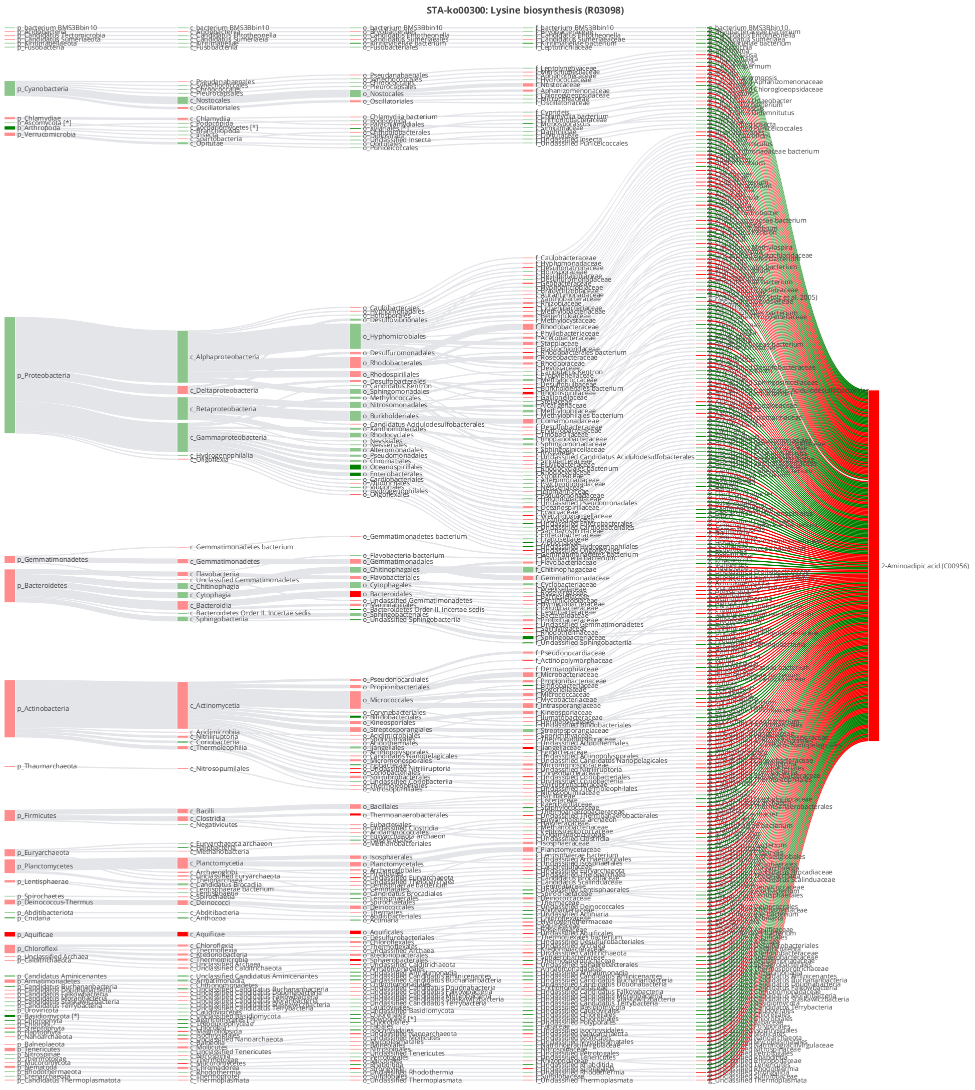


**Figure S59.** STA-Sankey network diagram (Hypergeometric test, log0.05 p-value > 1) for R03098 of lysine biosynthesis (ko00300) when comparing IP with CA condition (the reaction involving 2-aminoadipic acid (C00956) and L-2-aminoadipate reductase [EC:1.2.1.95] encoded by K00143)

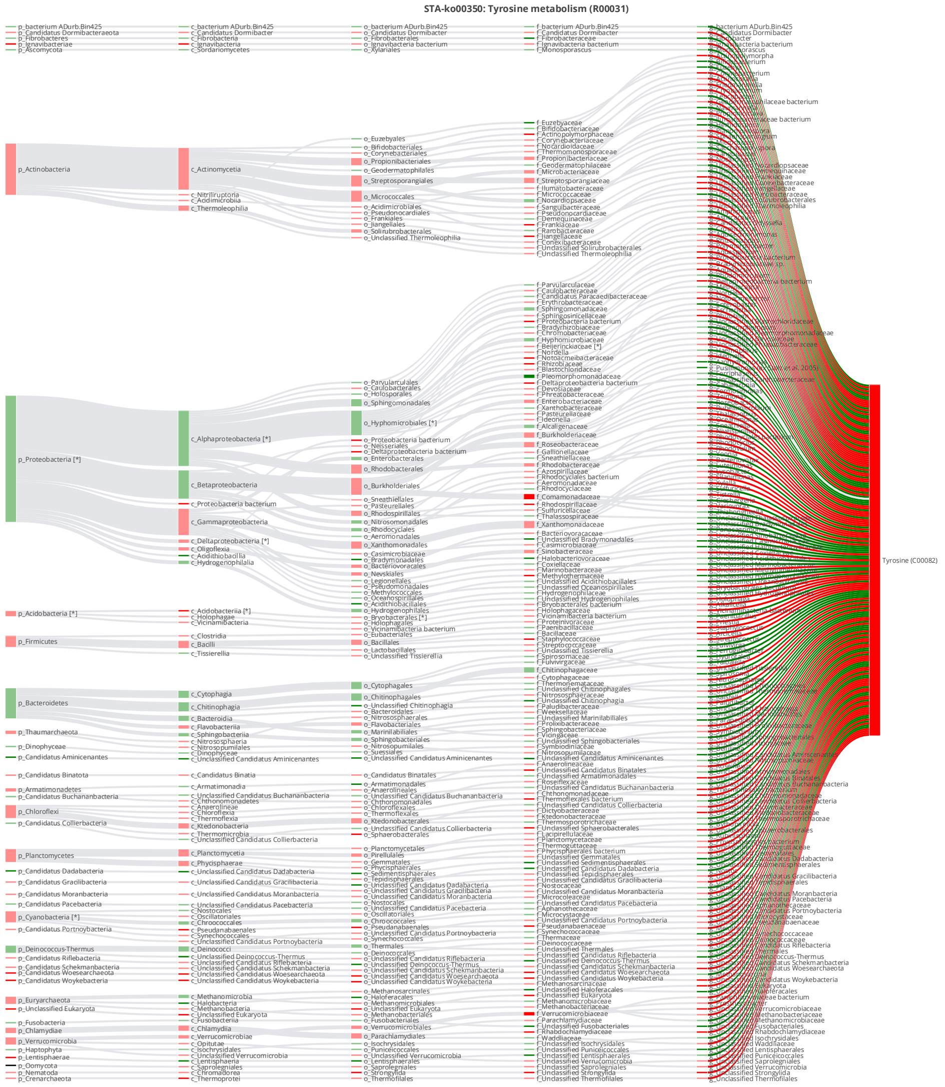


**Figure S60.** STA-Sankey network diagram (Hypergeometric test, log0.05 p-value > 1) for R00031 of tyrosine metabolism (ko00350) when comparing CP with CA condition (the reaction involving tyrosine (C00082) and polyphenol oxidase [EC:1.10.3.1] encoded by K00422)

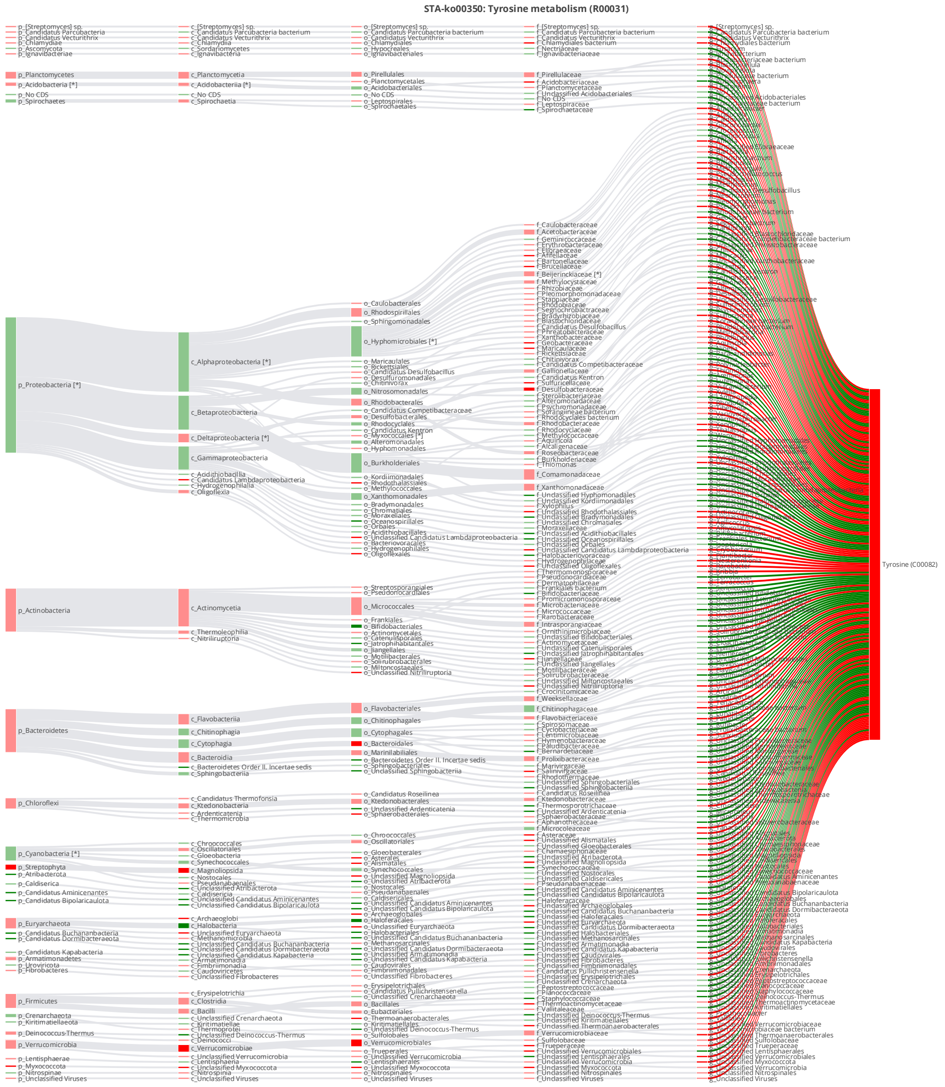


**Figure S61.** STA-Sankey network diagram (Hypergeometric test, log0.05 p-value > 1) for R00031 of tyrosine metabolism (ko00350) when comparing IP with CA condition (the reaction involving tyrosine (C00082) and polyphenol oxidase [EC:1.10.3.1] encoded by K00422)

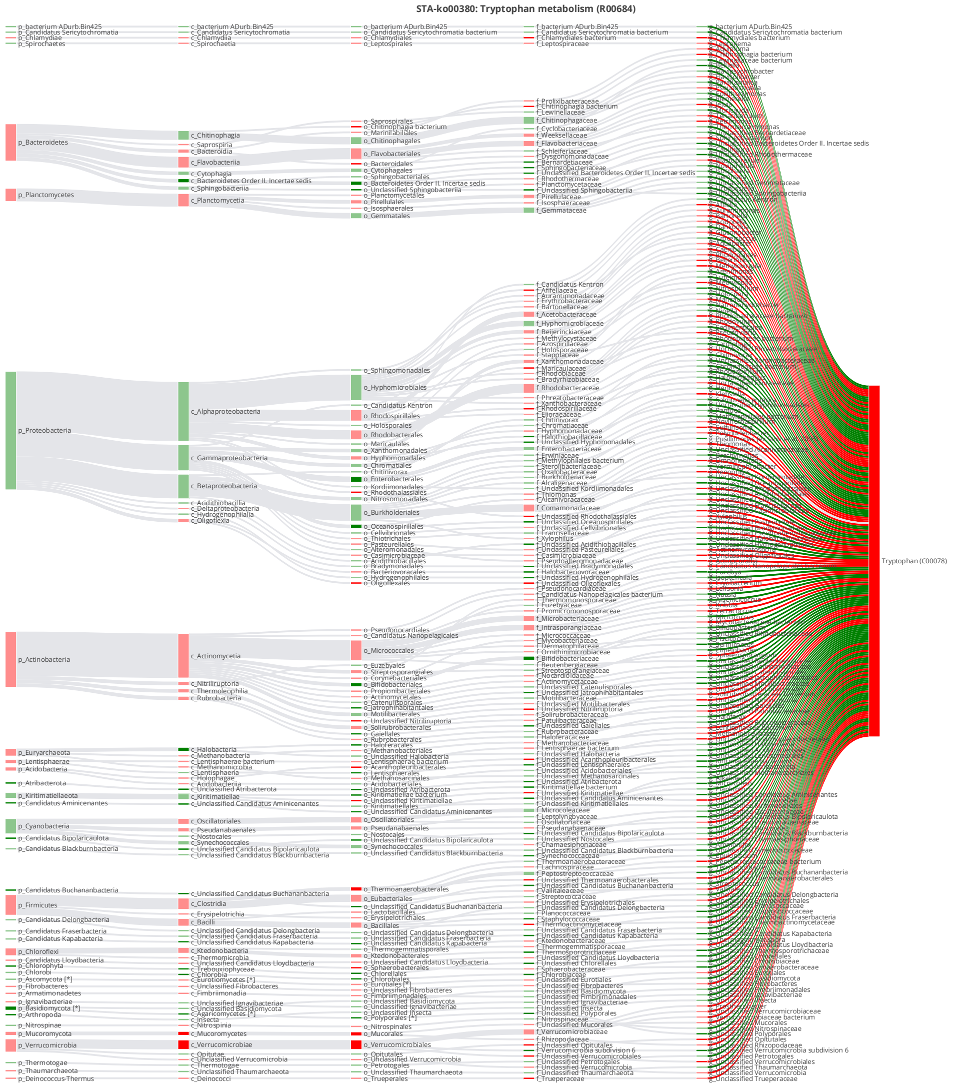


**Figure S62.** STA-Sankey network diagram (Hypergeometric test, log0.05 p-value > 1) for R00684 of tryptophan metabolism (ko00380) when comparing IP with CA condition (the reaction involving tryptophan (C00078) and tryptophan aminotransferase [EC:2.6.1.27] encoded by K14265)

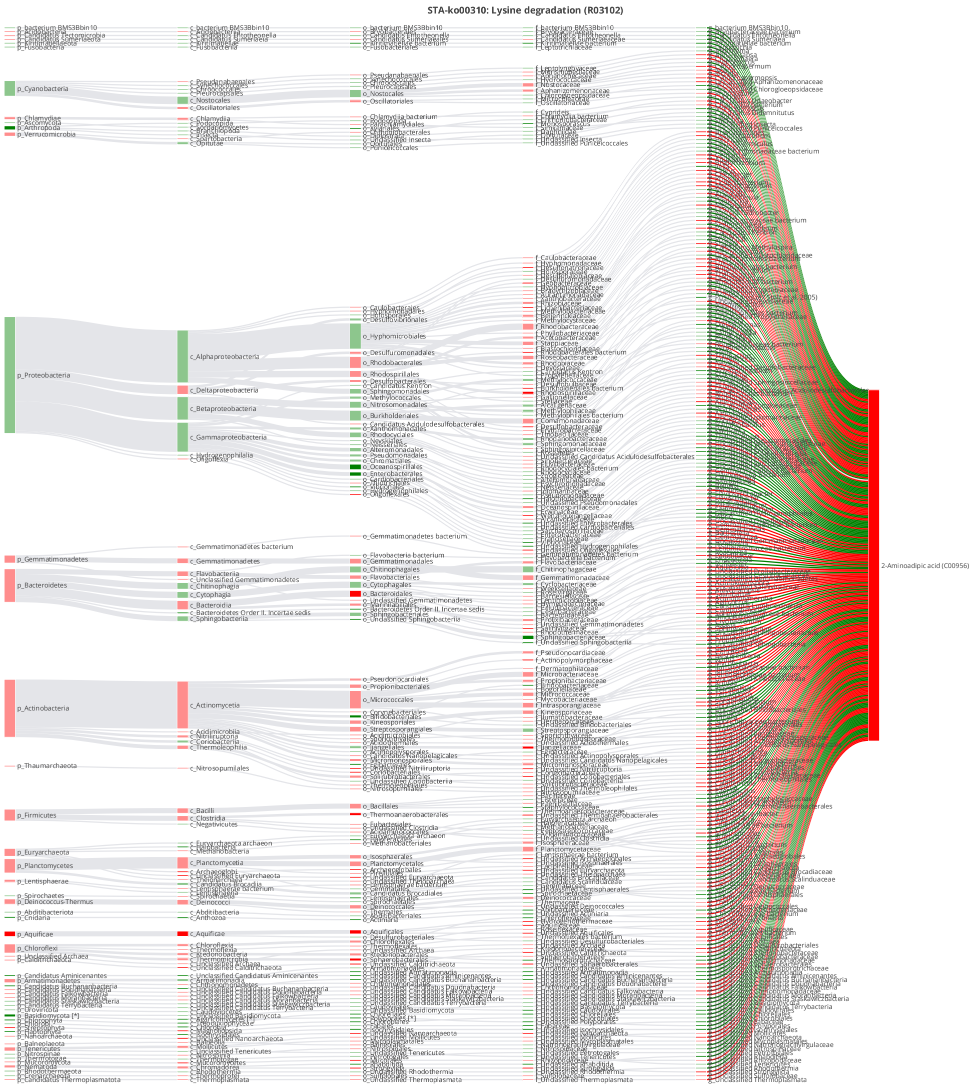


**Figure S63.** STA-Sankey network diagram (Hypergeometric test, log0.05 p-value > 1) for R03102 of tryptophan metabolism (ko00310) when comparing IP with CA condition (the reaction involving 2-aminoadipic acid (C00956) and aldehyde dehydrogenase family 9 member A1 [EC:1.2.1.47 1.2.1.3] encoded by K00149)

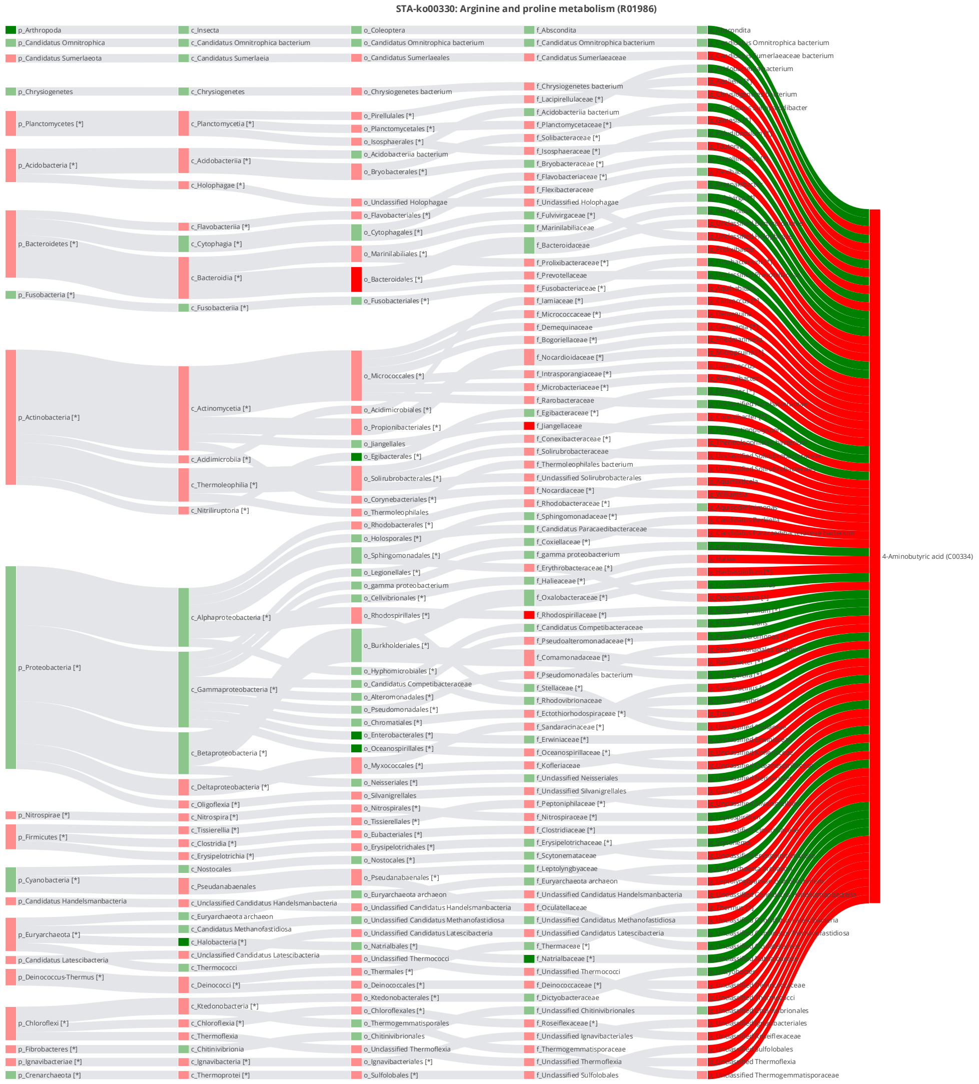


**Figure S64.** STA-Sankey network diagram (Hypergeometric test, log0.05 p-value > 1) for R01986 of arginine and proline metabolism(ko00330) when comparing IP with CA condition (the reaction involving 4-aminobutyric acid (C00334) and aldehyde dehydrogenase family 9 member A1 [EC:1.2.1.47 1.2.1.3] encoded by K00149)

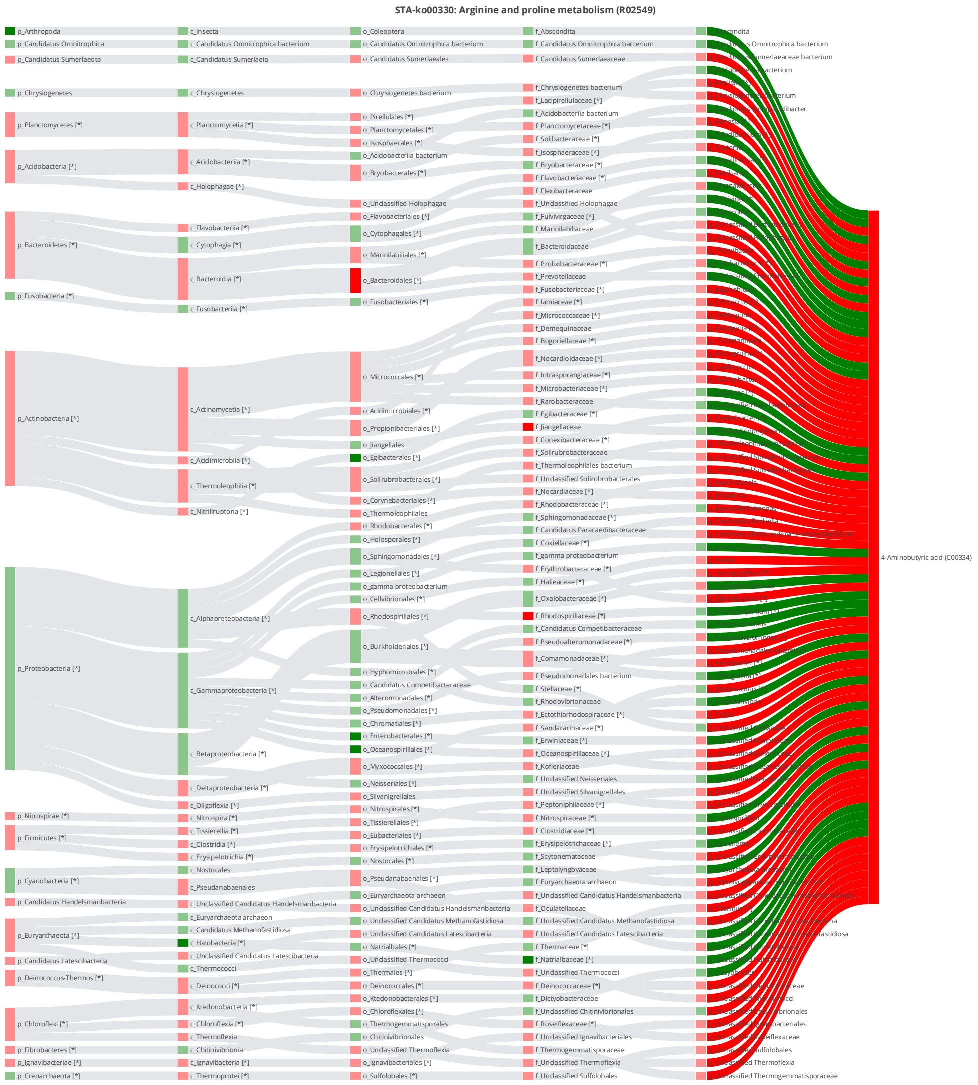


**Figure S65.** STA-Sankey network diagram (Hypergeometric test, log0.05 p-value > 1) for R02549 of arginine and proline metabolism(ko00330) when comparing IP with CA condition (the reaction involving 4-aminobutyric acid (C00334) and aldehyde dehydrogenase family 9 member A1 [EC:1.2.1.47 1.2.1.3] encoded by K00149)

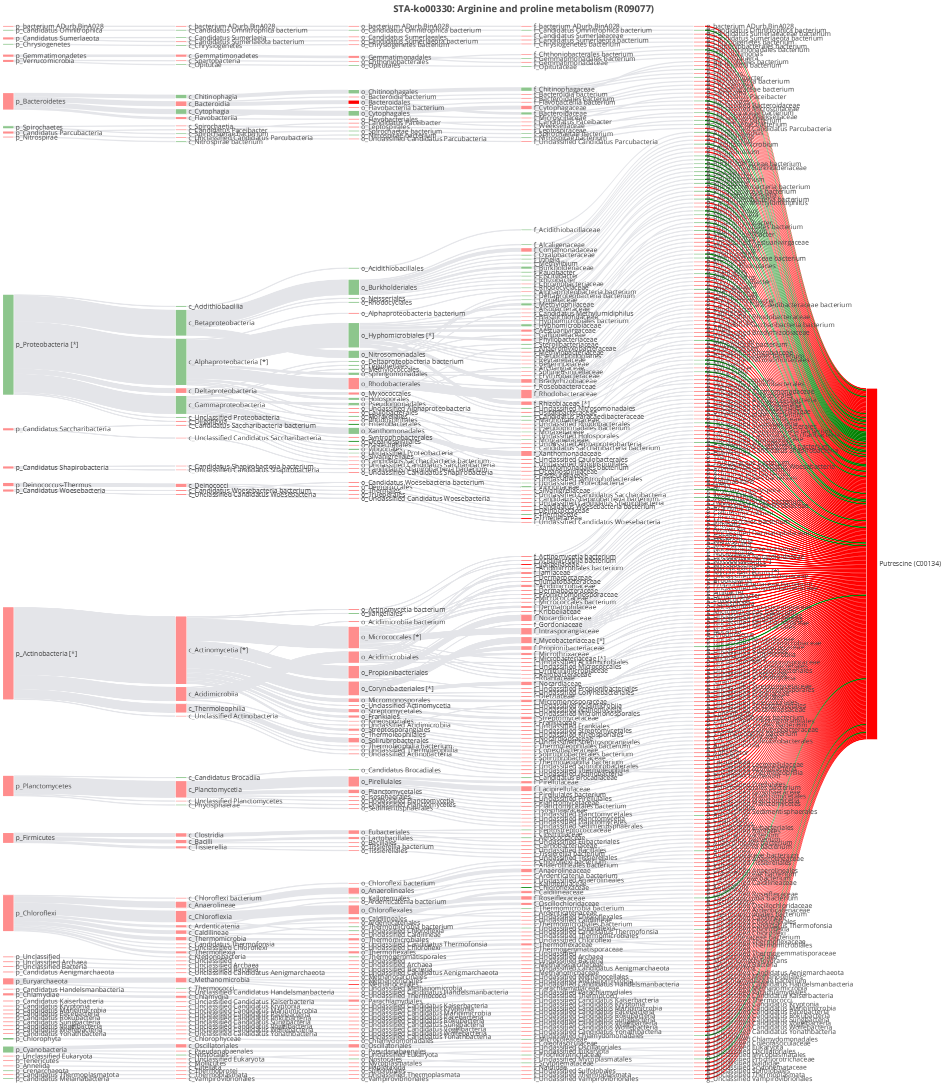


**Figure S66.** STA-Sankey network diagram (Hypergeometric test, log0.05 p-value > 1) for R09077 of arginine and proline metabolism(ko00330) when comparing IP with CA condition (the reaction involving putrescine (C00134) and polyamine oxidase [EC:1.5.3.17] encoded by K13367)

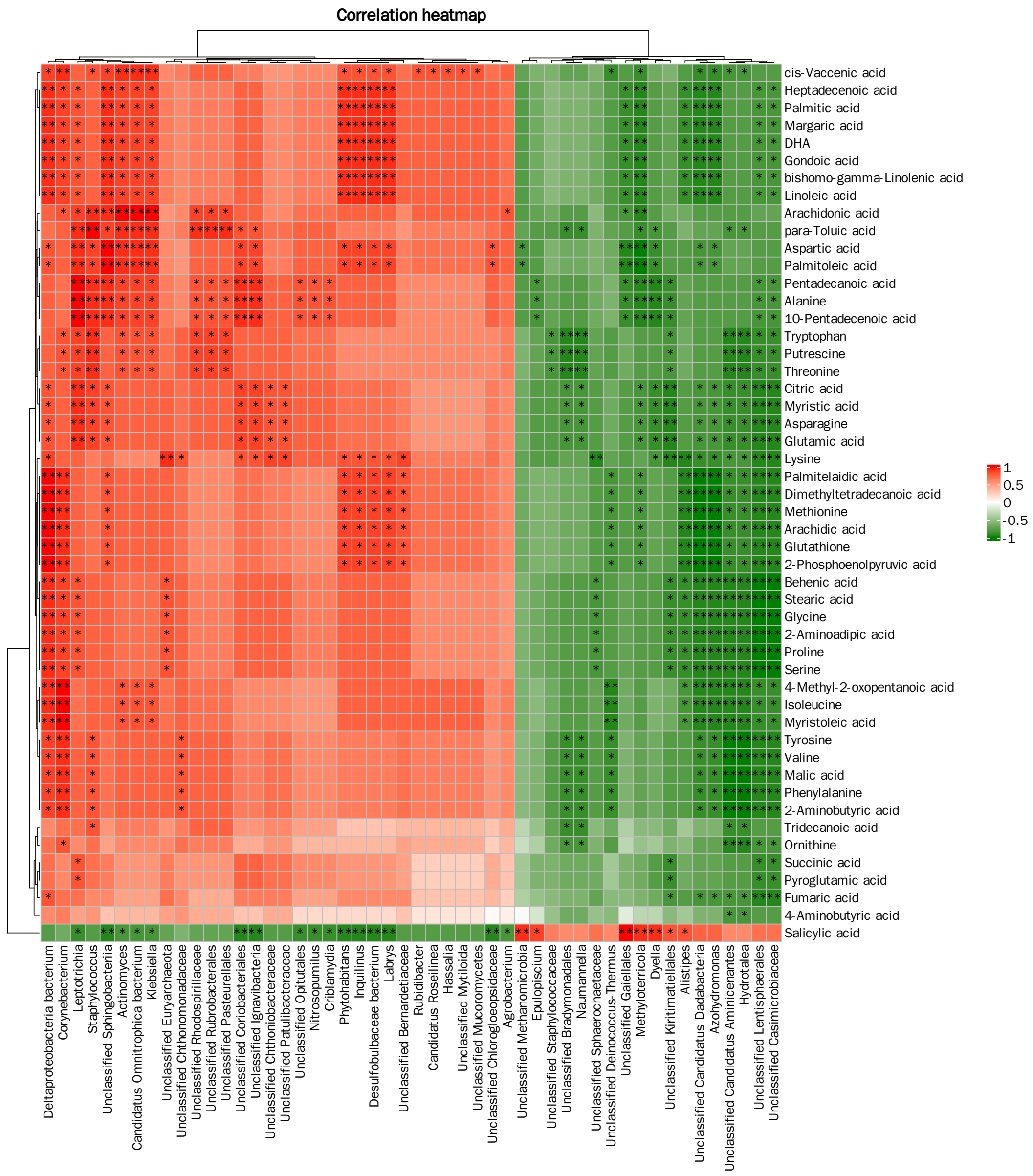


**Figure S67.** Correlation analysis of genera and metabolites in activated sludge samples based on the Spearman correlation method when comparing CP with CA condition (* indicates *P*<0.05, ** indicates *P*<0.01)

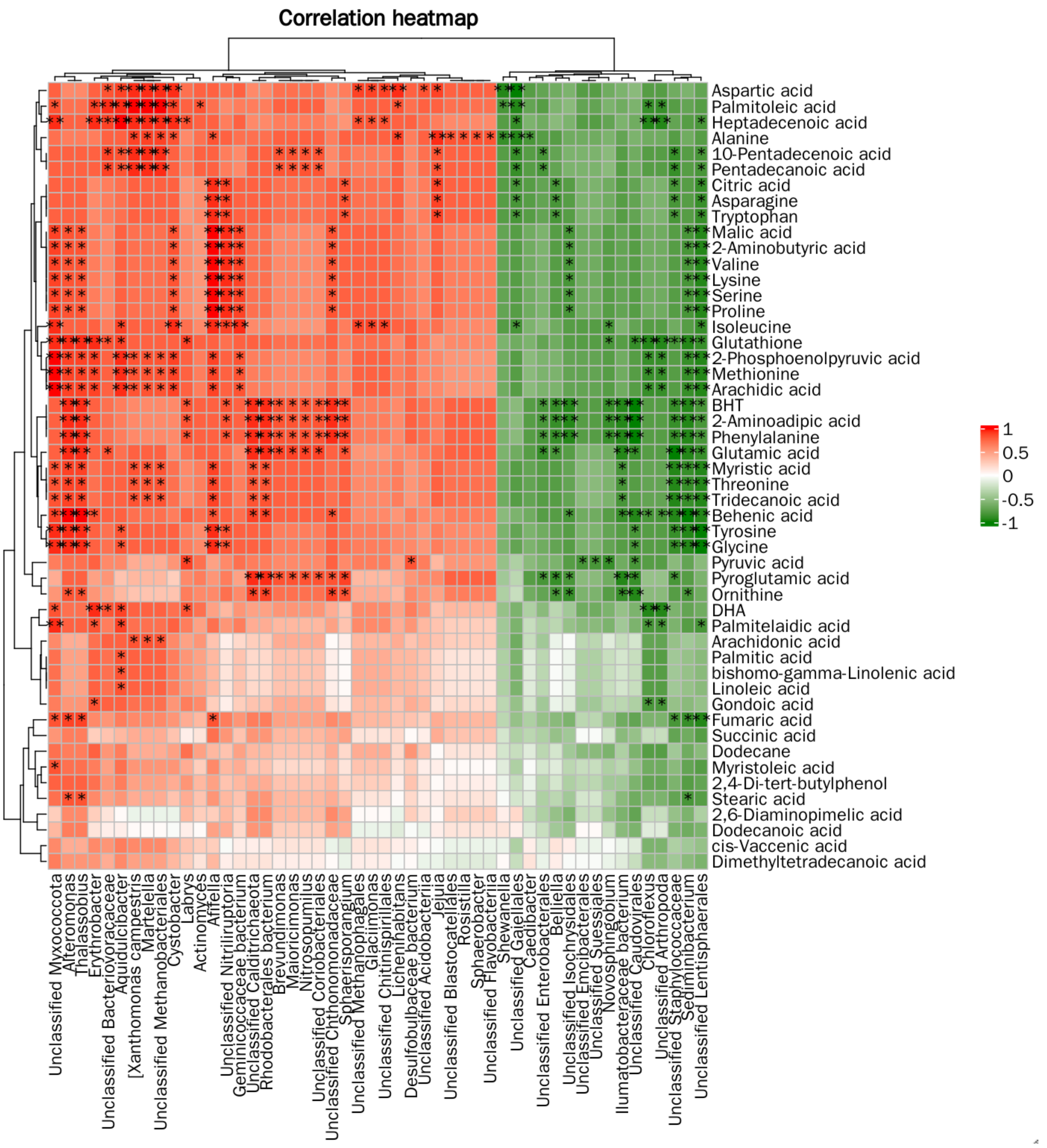


**Figure S68.** Correlation analysis of genera and metabolites in activated sludge samples based on the Spearman correlation method when comparing IP with CA condition (* indicates *P*<0.05, ** indicates *P*<0.01)
